# The molecular aetiology of tRNA synthetase depletion: induction of a *GCN4* amino acid starvation response despite homeostatic maintenance of charged tRNA levels

**DOI:** 10.1101/610790

**Authors:** Matthew R. McFarland, Corina D. Keller, Brandon M. Childers, Stephen A. Adeniyi, Holly Corrigall, Adélaïde Raguin, M. Carmen Romano, Ian Stansfield

**Affiliations:** Institute of Medical Sciences, School of Medicine, Medical Sciences and Nutrition, University of Aberdeen, Aberdeen, AB25 2ZD, UK; Institute of Complex Systems and Mathematical Biology, University of Aberdeen, Aberdeen, AB24 3UE, UK; MRC Protein Phosphorylation and Ubiquitylation Unit, School of Life Sciences, University of Dundee, Dundee, DD1 5EH, UK; Institut für Quantitative und Theoretische Biologie, Heinrich-Heine-Universität, 40225 Düsseldorf, Germany

**Keywords:** translation, tRNA synthetase, *Saccharomyces cerevisiae*, GCN4, Totally Asymmetric Simple Exclusion Process

## Abstract

During protein synthesis, charged tRNAs deliver amino acids to translating ribosomes, and are then re-charged by tRNA synthetases (aaRS). In humans, mutant aaRS cause a diversity of neurological disorders, but their molecular aetiologies are incompletely characterised. To understand system responses to aaRS depletion, the yeast glutamine aaRS gene (*GLN4*) was transcriptionally regulated using doxycycline by *tet-off* control. Depletion of Gln4p inhibited growth, and induced a *GCN4* amino acid starvation response, indicative of uncharged tRNA accumulation and Gcn2 kinase activation. Using a global model of translation that included aaRS recharging, Gln4p depletion was simulated, confirming slowed translation. Modelling also revealed that Gln4p depletion causes negative feedback that matches translational demand for Gln-tRNA^Gln^ to aaRS recharging capacity. This maintains normal charged tRNA^Gln^ levels despite Gln4p depletion, confirmed experimentally using tRNA Northern blotting. Model analysis resolves the paradox that Gln4p depletion triggers a *GCN4* response, despite maintenance of tRNA^Gln^ charging levels, revealing that normally, the aaRS population can sequester free, uncharged tRNAs during aminoacylation. Gln4p depletion reduces this sequestration capacity, allowing uncharged tRNA^Gln^ to interact with Gcn2 kinase. The study sheds new light on mutant aaRS disease aetiologies, and explains how aaRS sequestration of uncharged tRNAs can prevent *GCN4* activation under non-starvation conditions.

## INTRODUCTION

During gene expression, the translation of mRNA transcripts to produce a polypeptide chain involves the coordinated activities of an estimated repertoire of 200,000 ribosomes and 3 million tRNAs with associated translation factors, together representing a cellular production line that manufactures 70% of the dry weight of the cell in the form of protein (1). The delivery of amino acids to the ribosome by aminoacyl tRNAs, bound to GTP-charged elongation factor 1 (eEF1A in eukaryotes) is central to the translation process (2). Deposition of the aminoacyl tRNA in the ribosomal acceptor ‘A’ site by eEF1A is followed by the ribosome-mediated movement of the nascent polypeptide from the peptidyl ‘P’ site tRNA, to the newly-delivered amino-acyl tRNA in the ‘A’ site, thus lengthening the polypeptide chain by one amino acid (reviewed in; (2)). Coincident with ribosomal translocation, a now uncharged tRNA is released from the ribosomal exit ‘E’ site.

The released tRNAs must then be re-charged by one of a population of amino-acyl tRNA synthetases which covalently ligate amino acid to the correct tRNA in an ATP hydrolysis-linked reaction; most species have 20 different varieties of synthetase, one for each type of amino acid. (3). Following charging of the tRNA with amino acid, it can then again bind to eEF1A for delivery to the ribosomal A site. tRNAs thus cycle between charged and uncharged states, driven by the respective activities of tRNA synthetase and translating ribosome populations. Balancing of the demand of the translating ribosomes for aminoacyl tRNAs, with the capacity of the tRNA synthetase population to supply them, is vital to ensure translation can proceed without pausing. A mismatch between charged tRNA supply, and translational demand can cause accumulation of uncharged tRNA, with consequent activation of the Gcn2 kinase, triggering a *GCN4* amino acid starvation response (4).

Slow delivery of aminoacyl tRNAs to the ribosome triggers translational pausing, and a range of potentially detrimental outcomes. Failure to deliver a cognate tRNA to the A site can allow noncognate species to bind instead, with misincorporation of the amino acid (5). Extended translational pauses may trigger +1 ribosomal frameshifts, where the ribosome slips forwards by one nucleotide, before resuming triplet translation (6)(7). Specific ribosomal mechanisms exist to deal with pause events. In prokaryotes, tmRNA aided by the relE ribonuclease, triggers mRNA and nascent polypeptide degradation in response to pausing (8). In eukaryotes, no-go mRNA decay is triggered by the binding of the Dom34 and Hbs1 release factor paralogues to the paused ribosome, and release of the nascent peptide (9). Linked to the activity of Dom34 is a ribosome quality control pathway (RQC) that functions to resolve ribosomes stalled in hybrid states with occupied ‘A’ and ‘P’ sites (10). In yeast, proteins including Hel2 and Rqc2 are involved in detecting the stalled ribosome (11), while Rqc1 and Cdc48 participate in release of the nascent peptide from the large ribosomal subunit (12)(13)(14), with the nascent peptide being ubiquitinated as a result of the action of Ltn1 (13)(10). In some cases, degradation targeting is augmented by the Rqc2-mediated, non-ribosomal addition of C-terminal alanine and threonine (CAT) tails (15).

The adverse consequences of ribosomal pausing underscore the importance of the tRNA synthetase population in supplying aminoacyl tRNAs at the required rate. In humans, a number of Mendelian diseases are associated with tRNA synthetase mutations. Of the 37 amino-acyl tRNA synthetase genes (17 cytosolic, 17 mitochondrially-targeted, 3 bi-located), mutations in at least 28, many of them recessive, cause pathologies ranging from Charcot-Marie-Tooth (CMT) disease, through hypomyelination and epileptic encephalopathy, to cardiomyopathy and lactic acidosis (16). Semi-dominant mutations in cytosolic tRNA synthetases (GARS, YARS, AARS, MARS and KARS) have also been shown to result in various forms of Charcot-Marie-Tooth neuronal disorders (17)(18)(19)(20). Mutations in the human glutamine tRNA synthetase, QARS, can cause brain developmental disorders such as microcephaly (21)(22). The mechanisms behind these various conditions are still unclear, though *in vitro* biochemical assays, together with yeast complementation assays of several mutant tRNA synthetases, have indicated loss of aminoacylation activity (19)(23). This would be expected to result in a reduction in the charging level of the synthetase’s cognate tRNAs, and consequently impact on the translation of all mRNAs containing their cognate codons. Intriguingly, some of the aforementioned diseases are thought to be the result of mutations in protein domains responsible for noncanonical functions of the tRNA synthetase (24). Again, such mechanisms are still under investigation, but tRNA synthetases have been implicated in processes such as vascular development and mTOR signalling (25)(26). Almost all the tRNA synthetase Mendelian diseases are thus poorly understood in terms of the consequences for the translation system.

In this work we have established a model system to better understand the molecular consequences of loss of tRNA synthetase activity for translation. We analysed translation system and stress response pathways to define how the translation system reacts when the ribosomal delivery of tRNAs is impeded through use of a glutaminyl tRNA synthetase shut-off system, mimicking aspects of the human QARS gene loss of function mutations. A combination of experimental investigation and mathematical modelling was used to investigate the extent to which these mutations can have system-wide consequences for tRNA charging and stress responses. We show that the primary response of the cell to restriction of tRNA synthetase activity is a simultaneous slow-down of translation and growth rate, which homeostatically limits the accumulation of uncharged tRNA species, and prevents the formation of ribosome queues on the mRNA. Despite the homeostatic response, there is nevertheless an induction of the *GCN4* response, triggered by the activation of the Gcn2 kinase by uncharged tRNA. System modelling reveals that the reduced abundance of tRNA synthetase can interfere with the normal ability of the synthetase to sequester the uncharged tRNA pool. This failure to sequester allows uncharged tRNAs to interact with Gcn2 and induce an amino acid starvation stress response via the *GCN4* pathway. The study thus identifies a key sequestration role for the tRNA synthetase population in preventing unwanted triggering of amino acid starvation responses even while active translation continues to produce uncharged tRNA. The combined modelling and experimental analyses also define the potential range of mechanisms underpinning different molecular phenotypes of human tRNA synthetase mutations.

## MATERIAL AND METHODS

### *S. cerevisiae* strains and growth conditions

Yeast strains were grown at 30°C in YPD (2% glucose, 2% peptone, 1% yeast extract, all w/v) or synthetic defined (SD) media (2% glucose, 0.67% yeast nitrogen base (w/v), supplemented with required nucleotides and/or amino acids at 20 mg l^-1^ or 60 mg l^-1^ [leucine]). *E.coli* strains were grown at 37°C in Luria-Bertani (LB) media supplemented with 100μg/ml ampicillin to maintain plasmid selection. Where yeast cultures were treated with doxycycline to deplete production of a tRNA synthetase, doxycycline was added to the required concentration to YPD or SD medium and the culture grown over-night at 30°C, with 200 rpm shaking, until an optical density at 600 nm of between 0.5 and 0.8 was reached, representing mid-log phase. Inoculations were adjusted to ensure culture harvest was always at this point in the batch growth cycle to accommodate differences in growth rate caused by doxycycline.

To construct strains containing a regulatable *GLN4* gene, a *KanMX4-tTA-PtetO* cassette was amplified from pCM225 (27) using primer A1 (Table S1) with either primer A2 or A3, each including 45 nt of flanking sequence homology to the 5’ (A1) or 3’ (A2 and A3) of the target integration site upstream of the *GLN4* open reading frame; primer sequences are listed in Table S1. Primer A3 also encoded an N-terminal haemagluttinin (HA) sequence to tag to the *GLN4* coding sequence. Following amplification, KanMX4-tTA-PtetO cassettes were separately transformed into yeast strain BY4742 (*MATα his3-1 leu2-0 lys2-0 ura3-0),* and geneticin-resistant integrants selected (28). The same approach was used to place the genomic lysine tRNA synthetase gene *KRS1* under tet-off regulation using primers A15 and A16 (Table S1) to amplify the tet-off cassette from plasmid pCM225, and direct its integration immediately upstream of the genomic copy of *KRS1.*

The ablation of the *GCN2* gene was achieved using CRISPR-Cas9 (29). *tetO-GLN4* yeast were transformed using standard methods (28) with 125 ng SwaI/BclII-linearised pML104, (directing expression of Cas9, and the guide RNA) (29) and 70 ng of plasmid repair DNA fragment (Life Technologies, Table S1, sequence A8; this directs gap repair of pML104 by homologous recombination, inserting the gRNA sequence). Alongside these DNA fragments, 250ng of doublestranded genome repair DNA fragment (Life Technologies, Table S1, sequence A9) was cotransformed. This DNA fragment contained >75 nucleotides of sequence homologous to the genome immediately 5’, and 75 nt immediately 3’, to the *GCN2* gene, directing repair of the Cas9-induced double strand break and deletion of the *GCN2* gene by homologous recombination. Successful editing was confirmed in individual transformants using PCR sequencing with primers A10 and A11.

To construct strains containing translational fusion of ribosomal protein L25 (uL23 in the systematic nomenclature (30)) with C-terminal GFP, CRISPR-Cas9 was again used (29) as before. *tetO-GLN4* yeast were transformed with gapped pML104, repaired by homologous recombination in yeast during the transformation process, using a gRNA DNA fragment (sequence A14: Table S1). Alongside these DNA fragments, 3 μg of double-stranded genome repair DNA fragment (GFP open reading frame, PCR amplified using primers A12, A13, Table S1) was co-transformed to drive homologous recombination-directed integration of GFP in-frame with the YOL127W (uL23/RPL25) gene. Successful editing was confirmed in individual transformants using PCR sequencing of amplified genomic DNA.

### Plasmids

The activation of the *GCN4* response was assayed in yeast using plasmids p180, p226 and p227, which express β-galactosidase under the translational control of the yeast *GCN4* gene 5’ leader (31); a kind gift of Prof A. Hinnebusch). β-galactosidase assays were carried out using standard methods (32). Plasmids were transformed into *S. cerevisiae* using standard methods (28). Plasmid pET19b-*GLN4* was used to express the yeast *GLN4* gene, and GFP gene in *E. coli.* The HA-tagged *GLN4* gene (PCR-amplified using primers A4 and A5; Table S1) and His_6_-tagged GFP gene (primers A18 and 19; Table S1) were separately cloned using InFusion (Clonetech) methods into NdeI/BamHI-linearised pET19b (Novagen).

### Ribosome quantification using uL23-GFP

Actively growing mid-log phase 25 ml yeast cultures expressing GFP-tagged ribosomal protein uL23 (RPL25) were harvested at 3000 x *g* for 5 min. Washed cells were re-suspended in 0.5 ml lysis buffer [25 mM Tris-HCl (pH 7.2), 50 mM KCl, 5 mM MgCl_2_, 1 mM 2-mercaptoethanol with protease inhibitors, containing 50 ng of zymolyase to improve cell breakage. Cells were lysed with glass beads, centrifuged for 15 min at 13,000 rpm, and the supernatant retained to assay GFP fluorescence (485 nm excitation and 520 nm emission), using a FLUOstar Omega multi-well plate reader (LabTech International). The percentage of cells remaining unbroken (typically 25%) was calculated by measuring the fluorescence of the initial cell suspension before breakage, and again of the unbroken cells in the cell pellet following glass bead lysis. Using this estimate of the number of lysed cells, the uL23-GFP fluorescence units released per yeast cell were estimated, and this value converted to uL23-GFP mass using a standard curve derived from purified GFP expressed in *E. coli.* From this value, moles, and thus, uL23-GFP molecules per cell were calculated. All GFP was ribosome-associated in these strains (data not shown), thus measurements of uL23-GFP molecules per cell are a direct proxy for ribosomes per cell.

### Western blot analysis

Yeast protein extracts were prepared as described previously from three independent yeast cultures (33). For Western blots, following electrophoresis, proteins were transferred to Immobilon FL PVDF membrane (Merck-Millipore) using semi-dry transfer (25 V for 30 min.). HA-tagged Gln4 protein was detected using an anti-HA antibody (HA.11 clone 16B12, Cambridge Biosciences) and SuperSignal West Femto substrate (Thermo Scientific) using standard Western blot protocols (34). The resulting light output was quantified using an Alpha Innotech Multi-Image II camera. eIF2-α phosphorylation was detected using a phospho-eIF2α (Ser52) antibody (Thermo-Fisher; 44-728G; 1:10,000 dilution). Non-specific antibody detection of non-eIF2α proteins was reduced by pre-adsorbing the primary antibody to nitrocellulose blots of yeast extracts derived from cells grown in amino acid replete conditions (YPD medium), which do not contain phosphorylated eIF2-α. A glycolytic protein, phosphoglycerate kinase (Pgk1p) was immune-detected to control for lane loading (Abcam ab113687; 1:10,000 dilution).

### Expression of purified Gln4p and GFP

Recombinant HA-tagged Gln4 protein for use as a standard was produced by transforming *E.coli* strain Rosetta-gami B (DE3; Novagen) with pET19b-*GLN4*. Protein expression was induced in pET19b-*GLN4* transformants using 1 mM IPTG for 3 hours. Following harvest and lysis by sonication, TALON cobalt affinity resin (Clontech) was used to purify the His-tagged protein using manufacturer’s protocols, Samples were dialysed to remove excess salt and imidazole before mass spectrometry analysis using a Q-Exactive Mass Spectrometer (Thermo-Fisher) to confirm protein identity. Recombinant GFP for use as a standard for the determination of ribosome content was similarly produced using pET19b-GFP.

### Northern blot measurement of tRNA charging levels

tRNAs were extracted from yeast cultures using standard methods (35) with some modifications (36). tRNA samples were resuspended in sodium acetate buffer (pH 4.6) and stored at −80°C.

Northern blot analysis was carried out by separating 5 μg tRNA per lane on a 25 cm-long 8 M urea, 0.1 M sodium acetate (pH 5) denaturing acrylamide gel (10%: 19:1 acrylamide: bisacrylamide). Gels were run at 80 mA for 36 hours at 4°C with 7-hourly buffer re-circulation and semi-dry blotted onto GE-Healthcare Nylon N+ membrane using 1 x tris borate-EDTA (TBE) buffer at constant current (1 mA/cm^2^) for 1 hour at 4°C. tRNAs were crosslinked using UV (120,000 μJ/cm^2^), before probing using 3’ end biotin-labelled oligonucleotide probes; tRNA^Gln^_UUG_, tRNA^Lys^_CUU_ and tRNA^Arg^_UCU_ were detected using primers A6, A7 and A17 respectively (Table S1). Probe labelling was carried out using the Pierce Biotin 3’ End DNA Labelling Kit. Probing of the blot was performed using the North2South™ Chemiluminescent Hybridization and Detection Kit (Thermo-Fisher) according to the manufacturer’s instructions. Probe-tRNA hybridisations were carried out overnight at 42 °C, except the tRNA^Arg^_UCU_ probe hybridisation, which was carried out at 70 °C, reduced in 10 °C increments per hour to 40 °C after overnight incubation, before then washing and detection.

### RNA sequencing

Three independent yeast colonies were grown in 50 ml YPD media ± doxycycline as per the conditions detailed above. Total RNA was extracted from the cultures using a Nucleospin RNA kit (Macherey-Nagel) and RNA sequencing was carried out by the Centre for Genome Enabled Biology and Medicine at the university of Aberdeen using Illumina NextSeq 500. The RNA-sequencing data discussed in this publication have been deposited in NCBI’s Gene Expression Omnibus (37) and are accessible through GEO Series accession number GSE126435 (https://www.ncbi.nlm.nih.gov/geo/query/acc.cgi?acc=GSE126435). Library preparation, basecalling and quality control methods are described fully in the GEO database entry. Following quality control, the reads were mapped to the sequenced *S. cerevisiae* genome using TopHat for Illumina (38). Differential expression between test and control samples was assessed using CuffDiff (39).

### SILAC mass spectrometry

Stable isotopic labelling of amino acids in cell culture (SILAC) was carried out as previously described (40)(41). Triplicate yeast cultures were grown in ‘light’ (containing 20 mg/l ^12^C arginine and lysine) in the presence of 1μg/ml doxycycline to deplete the Gln4 tRNA synthetase, or using ‘heavy’ (containing 20 mg/l L-arginine-^13^C-HCl (CLM-2265-H, Cambridge Isotope Labs) and 20 mg/l L-lysine-^13^C-HCl [CLM-2247-H, Cambridge Isotope Labs]) SD media. Cultures were grown for at least 8 doublings to ensure steady-state ^13^C labelling of proteins, and harvested at approximately OD_600_ 0.5-0.6. Equal numbers of cells from heavy and light cultures were mixed, harvested and resuspended in lysis buffer [25 mM Tris-HCl (pH 7.2), 50 mM KCl, 5 mM 2-mercaptoethanol, 1 x cOmplete mini protease inhibitor (Roche)]. Yeast were glass bead-lysed and the lysate clarified using low speed centrifugation (10 min at 12,000 x rcf).

For SILAC mass spectrometry, 25 μg total protein was separated on a 12% SDS-PAGE gel, and 12 gel slices of equal size subjected to in-gel tryptic digestion and LC-MS/MS performed using a Q-Exactive Mass Spectrometer (Thermo-Fisher). Subsequent analysis of the mass spectra was carried out using MaxQuant software. Only protein groups with two or more unique peptides recorded were analysed. The mass spectrometry proteomics data have been deposited to the ProteomeXchange Consortium (http://proteomecentral.proteomexchange.org) via the PRIDE partner repository [77] with the dataset identifier PXD017140.

### Polysome profiling

Cells were grown to mid-log phase in YPD as described above and incubated with 200 μg/ml cycloheximide for 15 minutes prior to harvesting. Polysome analysis was performed according to standard methods (42). Gradients were unloaded using a BR-184 density fractionator (Brandel) connected to a UV monitor (254 nm; UV-1, Pharmacia Biotech) and a chart recorder. The area under each peak was quantified using a mass-based integration method.

### A stochastic model of the yeast translation system incorporating tRNA charging dynamics and ribosome competition

A global mathematical model of translation of mRNAs by ribosomes was developed, referred to as Global Translation Model (GTM) from now on available through Biomodels (76) with reference MODEL2001080004. The model is based on the Totally Asymmetric Simple Exclusion Process (TASEP) and it includes competition for ribosomes and tRNAs among the mRNAs, as well as competition among tRNAs for the aminoacyl-tRNA synthetases. The model also incorporates ribosome drop-off, where with a given probability the ribosomes detach from the mRNA before reaching a stop codon (43). mRNAs are represented by one-dimensional lattices, with each site of the lattice being a codon. Ribosomes are represented by particles with a footprint of 9 codons (44) that enter the lattice (5’ end) at a specific initiation rate and hop from one codon to the next until they reach the stop codon, at which point they leave the lattice, thereby producing a protein (Fig. 5A).

The model considers a virtual transcriptome comprising different types of mRNAs representative of the *S. cerevisiae* transcriptome. For each of the 93 gene ontology (GO-Slim) categories (gene assignments downloaded from the *Saccharomyces Genome Database* at https://downloads.yeastgenome.org/curation/literature/, 1.7.2017), for all mRNAs in each GO-Slim category, codon frequencies for each mRNA were summed, and multiplied by abundance of the parent mRNA to yield the GO-Slim category codon composition. Note that codon frequencies of genes assigned to more than one category were weighted accordingly. An mRNA of average length of codons for its GO-Slim class was then generated. Finally, the set of codons obtained in this way was randomised to produce 93 representative mRNA codon sequences. The abundance of each GO-Slim category within the cell, (see Table S2), determined its mRNA copy number within the simulation environment.

mRNAs compete for free (non-mRNA bound) ribosomes. Each different type of mRNA is associated with an initiation rate, which is dependent on the number of free ribosomes in the cytoplasm at any time point. In particular, we assume that the initiation rate is a stepwise linear function of the number of free ribosomes, similar to the approach used in (45). A ribosome either joining or leaving the mRNA respectively decrements or increments the pool of free ribosomes by one; the total ribosome pool size is fixed. The initial conditions are chosen so that the averaged initiation rate yields 〈*α*〉 = 0.15/*s* following the derivation in (46). Ribosomes exit the lattice at the stop codon, i.e., the last lattice site, with a constant termination rate identical for all mRNAs, that is considered to be not limiting (46). The initiation rate constant *α*_0,*i*,_, *i* = 1,…,93 for each of the GO-Slim category representative mRNAs was determined by taking the average over all initiation rate constants from (46) that are assigned to each single mRNA belonging to that category, weighted by the corresponding mRNA abundance (47). Other parameter values for the Global Translation Model are defined in the Supplementary Material.

The translocation to the next codon is not allowed if it is covered by another ribosome. The hopping rate along the lattice is codon-dependent, and is proportional to the number of cognate charged tRNAs available at that time point (48). The factor of proportionality is set to yield an average translation elongation rate (hopping rate) of 10 codons/s (49)(50)(51) and an average tRNA charging level 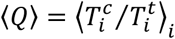 of 80%, as observed under physiological conditions (52)(36)(53)(54)(55)(see Fig. S1 for model sensitivity to this charging level). Wobble base-pairing is considered, so that synonymous codons can have different hopping rates (56). Details of derivation and parameters are provided in Supplementary Materials.

Every translation elongation event uses one charged tRNA 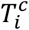, so that the corresponding charged tRNA (aa-tRNA) number with *i* = 1, …, 41 is decreased by one, and the number of uncharged tRNA 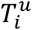 is increased by one. Note that the total number of tRNAs (defined in Table S3) of each type is fixed.

The change in the number of charged tRNA of each type 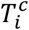 is governed by the main balance equation between supply and usage 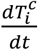 = *recharging rate — usage rate*.

The usage rate is proportional to the ribosomal current *k_j_ρ_j_*(l – *ρ*_*j*+1_) along the mRNA, since each elongation event utilises a charged tRNA (see Supplementary Materials for complete expression). The recharging rate 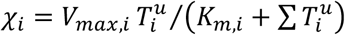 follows Michaelis-Menten dynamics with competitive inhibition, as isoacceptor tRNAs compete for the same tRNA-synthetase. Here *V_max,i_* represents the maximum charging rate and *K_m,i_* the Michaelis constant of the corresponding tRNA-synthetase. Uncharged tRNAs are then recharged with amino acids by the corresponding tRNA synthetase at the recharging rate *χ_i_*.

To mimic the inhibitory effect of doxycycline on the expression of Gln4 (glutaminyl tRNA synthetase) in the doxycycline-responsive Gln4 yeast strain, a range of Gln4 concentrations is considered, from 100% (corresponding to no doxycycline), to 3.5% of the wild type concentration (0.5 μg/ml doxycycline). This inflicts a change in the glutamine charging rate via 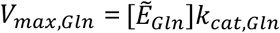, see Table S4 in Supplementary Materials. In the following we use 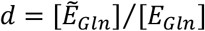, where 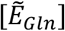 is the altered and [*E_Gln_*] is the wild-type synthetase concentration, to express the Gln4 protein ratio used in the simulations. With this procedure we can also overexpress Gln4, thereby yielding a Gln4 protein ratio larger than 1 (see Fig. 5D-G), which corresponds to a concentration of Gln4 of up to 200% of wild-type, achieved experimentally because the doxycycline-responsive promoter is stronger than the native *GLN4* promoter (data not shown).

## RESULTS

### Gln4p expression becomes growth rate limiting below physiological levels

Mutations in a wide range of aminoacyl tRNA synthetases cause human Mendelian disorders with neurological phenotypes. To better understand how mutations in tRNA synthetase genes cause tissue-specific human disease, we sought to establish the range of molecular phenotypes triggered by loss of tRNA synthetase activity, using yeast as a model system. We chose to deplete the *S.cerevisiae* glutamine tRNA synthetase protein, encoded by the *GLN4* gene. The native *GLN4* promoter was replaced with a tet-off regulatory cassette (27), allowing its transcription to be repressed through the addition of doxycycline. As expected for an essential gene, the tet-off *GLN4* strain shows dose-dependent sensitivity to doxycycline and reductions in growth rate when grown both on solid and in liquid media supplemented with the antibiotic (Fig. 1A-B).

**Figure 1:**
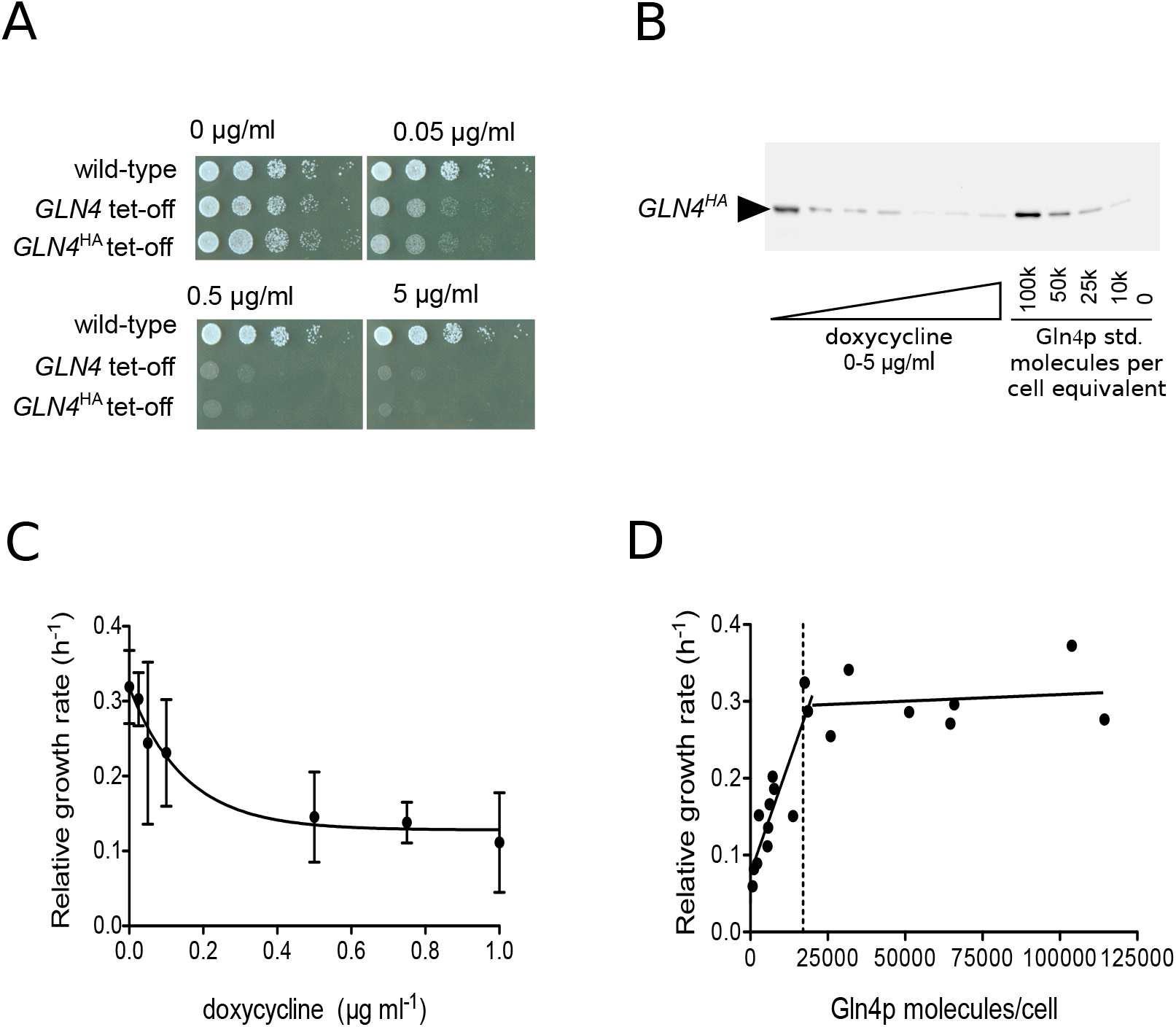
Negative growth regulation by a doxycycline-controlled glutamine tRNA synthetase gene *GLN4*. *Panel A:* The growth of a wild-type strain on a range of doxycycline-containing agar was compared to that of a tet-off *GLN4* integrant, and a tet-off *GLN4* integrant C-terminally tagged with HA epitope. *Panel B:* Using the engineered C-terminal epitope tag on Gln4p, Western blot band intensities were quantitatively compare to an HA-tagged Gln4p standard expressed in *E. coli,* using a standard curve (not shown). *Panel C:* The growth rate constant for a tet-off *GLN4* integrant C-terminally tagged with HA epitope was measured across a range of doxycycline concentrations. [D] Using quantitative Western blots, the cellular content of Gln4p was calculated allowing growth rate to be fitted biphasically to Gln4 content. The vertical dashed line indicates the cellular content of Gln4p estimated in a proteomic meta-study (58).

Previous research has shown that the *S.cerevisiae* translation system is robust to reductions in the abundance of tRNA synthetases, since for the majority of the 20 synthetases, heterozygous deletions are tolerated (57). However, here we show that in response to doxycycline, inhibition of *GLN4* transcription significantly impacts cellular growth (Fig. 1). To establish the cellular requirement for glutamine tRNA synthetase, we quantified the relationship between cellular content of Gln4p, and growth rate. Using purified HA-tagged (His)_10_-Gln4p expressed in *E. coli,* we carried out quantitative Western blot analysis on mid-log phase lysates of P_tet-off_ *GLN4* strain yeast grown at different concentrations of doxycycline. The Gln4p content of each sample was calculated using a recombinant Gln4p standard curve (Fig. 1), and plotted against the relative growth rate of that sample at the time of harvest (Fig. 1D).

The resulting plot shows that the cellular growth rate remained relatively consistent until Gln4p abundance reduced below approximately 20,000 molecules per cell, after which growth rate decreased markedly, with a biphasic response. Intriguingly, the threshold value below which the growth rate is reduced is close to estimates for the physiological Gln4p abundance of 17,000 molecules per cell (58)(59), suggesting that normally the level of glutamine tRNA synthetase is maintained at level just above that which is rate-limiting for growth.

### *GCN4* translation levels are enhanced by Gln4p depletion

Previous research on the effects of a small molecule inhibitor of the human prolyl tRNA synthetase PARS revealed that it caused induction of the mammalian GCN2-ATF4 pathway, a cellular response to amino acid starvation triggered by the accumulation of uncharged tRNA (60). It was therefore important to establish whether a similar *GCN4* amino acid starvation response was triggered by depletion of the glutamine tRNA synthetase in response to doxycycline, due to an expected increase in uncharged glutamine tRNA in the cell. This should activate Gcn2 kinase activity, phosphorylate translation initiation factor eIF2, and thus de-repress translation of *GCN4* (ATF4 in mammals) by bypassing inhibitory uORF sequences in its 5’UTR.

This hypothesis was tested through use of a reporter assay in which the 5’ UTR and first 55 codons of *GCN4* are translationally fused to the lacZ gene, allowing β-galactosidase activity to report Gcn4p expression (Fig. 2). Wild-type yeast, and tetO-*GLN4* yeast, were transformed with plasmid p180, carrying uORF1 and uORF4, which reports *GCN4* activation, or p226, a negative control carrying the translation-attenuating uORF4 only. Each of these constructs was grown in a range of doxycycline concentrations and assayed for β-galactosidase activity. The results were expressed relative to those obtained for a 100% expression control (p227), which lacks any uORFs (Fig. 2).

**Figure 2:**
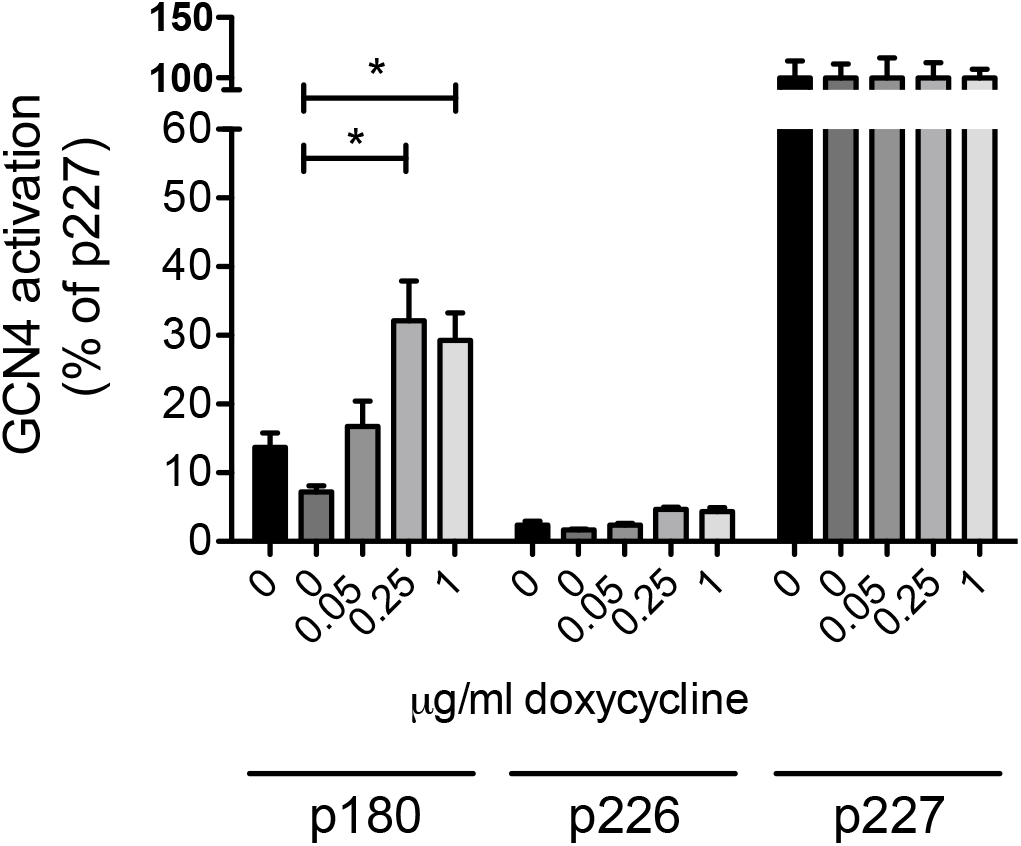
A *GLN4* tRNA synthetase shut-off induces a reporter of *GCN4* translation. The effect of *GLN4* tRNA synthetase shut-off, using doxycycline, on uncharged tRNA accumulation and thus *GCN4* activation was explored using the *GCN4-lacZ* reporter plasmids. The effect of doxycycline on *GCN4* induction was tested in a wild-type cell (BY4742, black bars, left-most in each block), and in a tetO-*GLN4* strain (remaining bars). Error bars represent ± 1 standard deviation, n=3. Significance at the *p* < 0.05 level is indicated by an asterisk. Plasmid p180 measures a *GCN4* response, relative to the negative (p226) and positive (p227) controls.

In response to increasing concentrations of doxycycline, the level of β-galactosidase activity detected in the tetO-*GLN4* yeast increased significantly from 7% in the absence of doxycycline, to 30% of positive control (p227) levels at 0.25 μg ml^-1^, beyond which the *GCN4* induction response reached a plateau. This indicated that following doxycycline treatment, *GCN4* translation was induced, presumably in response to accumulation of uncharged glutamine tRNAs. We confirmed this result by creating another tRNA synthetase doxycycline shut-off strain, this time involving the lysyl-tRNA synthetase *KRS1.* Confirming the phenotype of the *GLN4* shut-off, the tet-off *KRS1* strain also exhibited doxycycline-dependent decreases in growth rate, coincident with *GCN4* induction (Fig. S2). In fact *KRS1* was originally identified as *GCD5,* a *general control derepressed* gene, mutations in which exhibit constitutive expression of *GCN4* (61). Using immuno-blot detection, we additionally confirmed the *GLN4*-tet-off mediated *GCN4* induction was occurring as expected via phosphorylation of phospho-Ser51 of eIF2α (Sui2p in yeast; Fig. S3). Together, the data clearly shows that tRNA synthetase depletion induces Gcn2p-dependent *GCN4* responses.

### Depletion of Gln4p activates Gcn4p-mediated transcriptional starvation responses

In order to confirm the gene regulatory phenotype triggered by tRNA synthetase depletion we carried out a transcriptomic analysis of the tetO-*GLN4* yeast in the presence and absence of doxycycline. Following the discovery that a *GCN4* reporter was induced by doxycycline treatment (Fig. 2) we expected to see a transcriptional response involving the amino acid biosynthetic genes.

RNA sequencing was carried out on wild-type and tetO-*GLN4* yeast grown in the presence or absence of doxycycline in order to investigate changes in gene expression in response to Gln4p depletion. For the wild-type strain, no significant differences in expression were detected for any gene, confirming doxycycline itself caused no transcriptomic changes in wild-type yeast in our hands (data not shown). In the case of the tetO-*GLN4* strain dataset, approximately two-thirds of the genes tested showed significant differences in expression in response to doxycycline treatment (*false discovery rate;* q < 0.05); of these, approximately 1200 genes were either 2-fold up-, or down-regulated.

These transcriptional response to Gln4 shut-off was compared against a previously published dataset in which *GCN4* was induced by amino acid starvation (62). Following merging of the datasets and the removal of non-significant values (where p>0.05), and instances where the induction or repression ratio was less than 2-fold, expression data for 534 genes matched across the two datasets was available. Overall the *GLN4* shut-off and *GCN4* induction datasets were positively correlated (R^2^=0.38), and when the subset of 2-fold induced/repressed and significant genes were alone considered, this correlation index was further increased (R^2^=0.595; Fig. 3A) indicating a strong similarity in transcriptional response.

**Figure 3:**
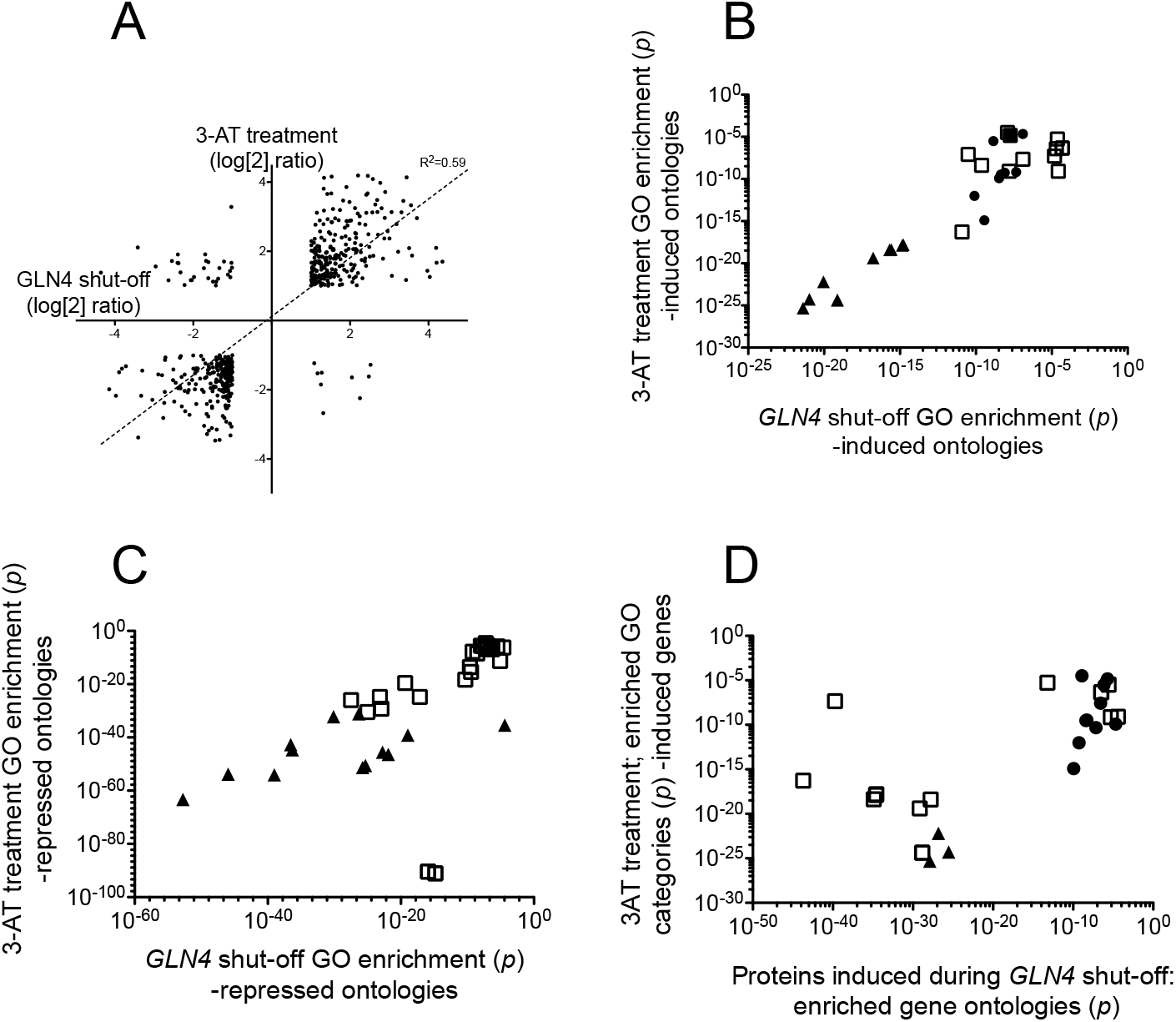
tRNA synthetase shut-off induces a *GCN4* amino acid starvation phenotype. *Panel A:* Transcript profile analysis of the *GLN4*-shut-off response; comparison of effects of Gln4p depletion and 3AT-induced starvation response on gene expression. Gene expression responses to depletion of Gln4p in the presence and absence of doxycycline was compared with that of a wildtype yeast strain in response to treatment with 100 mM 3AT, which induces an amino acid starvation response through inhibition of histidine biosynthesis. Log2 transforms of mRNA induction ratios were plotted for all genes that were common to both datasets in being; (i) statistically significant, and (ii) either >2-fold induced or <2-fold repressed. A line-of-best fit regression analysis is presented. *Panel B:* Gene ontology terms (i) over-represented among *upregulated* genes in response to glutamine tRNA synthetase depletion and (ii) which were also shared with GO classes enriched among genes upregulated in response to 3AT treatment (*p* <0.05). The corresponding *p*-values of each GO enrichment (*GLN4* and 3-AT) were plotted for every shared GO class. Filled triangles; ontologies including; cellular amino acid biosynthesis, amino acid metabolism ontologies. Filled circles; specific amino acid biosynthetic ontologies (Asp, Met, Arg, sulfur-containing amino acids). A list of all plotted ontologies is presented in Table S5. *Panel C:* Gene ontology terms (i) over-represented among *downregulated* genes in response to glutamine tRNA synthetase depletion and (ii) which were also shared with GO classes enriched among genes downregulated in response to 3AT treatment (*p* <0.05). The corresponding *p*-values of each GO enrichment (*GLN4* and 3-AT) were plotted for every shared GO class. Triangle symbols; ontologies including; translation, peptide biosynthetic processes, structural constituent of ribosome, cytosolic small and large ribosomal subunit, peptide metabolic process, ribosome. A full list of all plotted ontologies is presented in Table S6. *Panel D:* Gene ontology terms (i) over-represented among *upregulated* proteins (SILAC analysis) in response to glutamine tRNA synthetase depletion and (ii) which were also shared with GO classes enriched among genes upregulated in response to 3AT treatment (*p* <0.05). The corresponding pairwise *p*-values of each GO enrichment (*GLN4* and 3-AT) were plotted for all shared GO classes. Filled triangles; ontologies including; cellular amino acid biosynthesis, amino acid metabolism ontologies. Filled circles; specific amino acid biosynthetic ontologies (Asp, Met, Arg, sulfur-containing amino acids). A full list of all plotted ontologies is presented in Table S7.

These lists of genes were analysed for gene ontology (GO) term enrichment (63). Among upregulated genes in each dataset, approximately 50 statistically significant GO terms were enriched in the *GCN4* and *GLN4* data sets. Of these, 31 were common to both datasets, with the majority corresponding to amino acid biosynthetic processes. This corresponded to a Jaccard (*J*) overlap index of 0.42; Jaccard indices range from 1 (complete overlap) to 0 (no overlap of significant GO categories) where 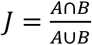 i.e. intersection over union for sets *A* and *B.* The 31 GO categories of over-expressed genes common to the *GCN4* and *GLN4* datasets also showed highly correlated *p* values for GO enrichment (R^2^ = 0.88; not shown).

In a similar manner, the list of genes downregulated in both *GLN4* and *GCN4* experiments (Fig. 3C) yielded 41 over-represented and significant GO terms common to both datasets that centred on translation and ribosome biogenesis. Together these results support the hypothesis that depletion of Gln4p triggers the induction of a *GCN4*-mediated amino acid starvation response, in agreement with the observed increased activation of *GCN4* mRNA translation exhibited in response to doxycycline treatment (Fig. 2).

To further corroborate these results, and to attempt to identify particular proteins whose expression is particularly affected by compromised glutamine tRNA charging, changes in the tetO-*GLN4* proteome in response to Gln4p depletion were examined via SILAC mass spectrometry, with the tetO-*GLN4* strain grown in “light” (+ 1 μg/ml doxycycline) or “heavy” (0 μg/ml doxycycline) minimal media. Heavy/light ratios were recorded for approximately 785 proteins. As before, a threshold log2 (fold change) value of ± 1 was set to indicate potential biological relevance.

Analysis of the heavy/light ratios of 15,000 peptides revealed 215 proteins showing increased expression while just 21 were detected with reduced expression at 2-fold reduction. Reducing this stringency to 1.4-fold then included a larger group of proteins in which was a significant number of ribosomal proteins (*p*=1.7E-10), consistent with the *GCN4* response which defines a shut-down of ribosomal synthesis (62) (Table S8). Gene ontology analysis of the proteins SILAC-enriched by doxycycline treatment showed clear enrichment for multiple amino acid biosynthetic pathways, both general (alpha amino acid biosynthesis) and specific (biosynthesis of Arg, Lys, Gln and Asp respectively). GO class enrichment in the SILAC *GLN4* shut-off data showed a Jaccard overlap index of 0.29 with the *GCN4* transcriptomic dataset over 28 gene ontology classes, mirroring the results of the *GLN4* shut-off transcriptomics. Similar ontological classes were over-represented in the SILAC protein analysis during *GLN4* depletion as during 3-AT-treated transcriptomic analysis (Fig. 3D). The analysis clearly indicates that Gln tRNA synthetase depletion induces transcriptional and proteomic responses mapping closely onto the *GCN4* induction response, with the induction of amino acid biosynthetic pathways.

### Glutamine tRNA synthetase shut-off does not trigger ribosome queues on polysomes

The depletion of glutamine tRNA synthetase is assumed to cause defects in tRNA^Gln^ charging, leading to slow growth. The assumption was that uncharged tRNA was accumulating, causing Gcn2p kinase activation and a Gcn4p transcriptional response (Fig 2, 3). We therefore examined whether a shortage of glutamine tRNAs was causing a ribosomal queuing phenotype, leading to an altered polysome profile indicative of the accumulation of large polysomes.

Accordingly, ribosomal extracts from cycloheximide-treated cells in mid-log phase were prepared. TetO-*GLN4* yeast were grown in doxycycline overnight, establishing a steady state of reduced Gln4p. We also treated the tetO-*GLN4* yeast strain with a short, defined doxycycline time-course. We reasoned that if there were homeostatic cellular responses to Gln4p depletion that allowed cells in some way to adjust to the imposed restriction, treating cells for a short period with doxycycline may reveal ribosome queuing effects that would otherwise be masked by steady-state doxycycline exposures.

Cells were therefore grown overnight to steady state in medium containing either 0, 1μg ml^-1^ or 5μg ml^-1^ doxycycline. Cycloheximide was added to freeze the polysomes on the mRNA, and polysomes analysed using sucrose gradients (Fig. 4). The results showed clearly that there was no accumulation of large polysomes in the doxycycline-treated cultures, and in fact the ratio of polysomes to monosomes decreased, indicative of a weak initiation block (Fig. 4A, B). We therefore used a time-course protocol, where cells at early log phase (OD_600_ =0.1), were exposed to doxycycline for either 4 or 8 hours, and polysomes analysed. However, the results showed only a mild reduction in the ratio of polysomes to monosomes, again indicative of a weak initiation block (Fig. 4A).

**Figure 4:**
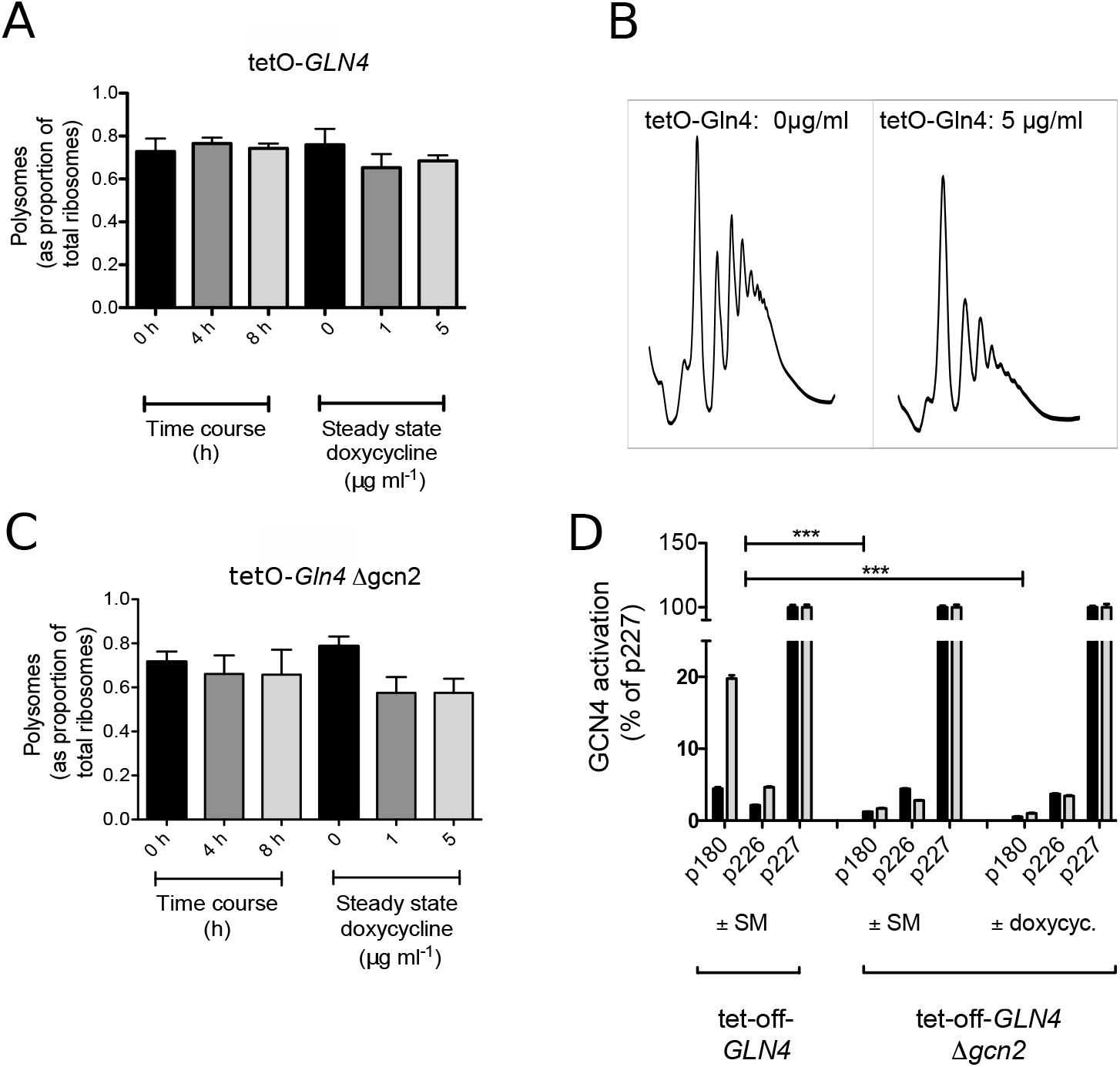
tRNA synthetase depletion does not cause the accumulation of larger polysomes. *Panel A:* Polysomal ribosomes inhibited with cycloheximide were extracted from a tetO-*GLN4* yeast strain either grown overnight to steady state in either 0, 1 or 5 μg ml^-1^ doxycycline, or exposed to 5 μg ml^-1^ for 0, 4h or 8h. Triplicate gradients were run for each condition, error bars represent ± 1 standard deviation, n=3. Significance at the p<0.05 level is indicated by an asterisk. *Panel B:* typical polysome traces are shown for a tetO-*GLN4* wild-type untreated with doxycycline, (left) or treated with 5 μg ml^-1^ doxycycline overnight (right), showing some loss of polysomes but no accumulation of large polysomes. *Panel C:* Polysomal ribosomes inhibited with cycloheximide were extracted from a tetO-*GLN4* Δ*gcn2* yeast strain either grown overnight to steady state in either 0, 1 or 5 μg ml^-1^ doxycycline, or exposed to 5 μg ml^-1^ for 0, 4h or 8h. Triplicate gradients were run for each condition, error bars represent ± 1 standard deviation, n=3. Significance at the *p* < 0.05 level is indicated by an asterisk. *Panel D:* The non-*GCN4* inducing phenotype of the *Dgcn2* mutant was tested using the *GCN4*-response lacZ reporter plasmids p180, p226 and p227. The response of the tetO-*GLN4* p180 transformant to SM reveals the expected *GCN4* induction response of approximately 20%. This response is lacking in the tetO-*GLN4 Dgcn2* deleted strain (p180 bars) treated with either SM or 5 μg ml^-1^ doxycycline. Control lacZ measurements (grey bars) are compared with those in the presence of either the isoleucine synthesis inhibitor sulfometuron methyl (SM), or 5 μg ml^-1^ doxycycline (black bars), as indicated.

We reasoned that since the doxycycline-induced Gln4p depletion caused a *GCN4* response, it was possible that Gcn2 kinase phosphorylation of the eIF2 initiation factor would cause the initiation block indicated by the polysome profiles. The *GCN2* gene was therefore deleted using CRISPR-Cas9. To confirm the effect of the deletion, we showed it was no longer possible to induce a *GCN4* response by treatment with sulfometuron methyl, an inhibitor of amino acid biosynthesis, nor by treatment with doxycycline (Fig. 4D). The latter experiment confirms that the induction of Gcn4 through doxycycline treatment in the tet-off *GLN4* strain is mediated via Gcn2 kinase, and validates the results of phospho-eIF2 immuno-blotting (Fig. S2). Repetition of the polysome analyses, using either the steady state or time-course protocols again revealed profiles typical of weak initiation blocks, rather than of ribosomal queuing with accumulation of large polysomes (Fig 4C). Overall the results indicate that in response to tRNA synthetase depletion, despite evidence for the accumulation of uncharged tRNA and *GCN4* induction, there was nevertheless no evidence for ribosome queuing, even when *GCN2* was ablated.

### Simulation of the effects on synthetase depletion using a global model of translation; an autogenous feedback loop links synthetase activity to growth rate and maintains levels of charged tRNA

A global mathematical model of translation (Global Translation Model) of mRNAs by ribosomes was developed, which defines a population of ribosomes, tRNAs and codon biased transcriptome as described (Materials and Methods). The model includes a description of the tRNA charging process by a population of 20 types of tRNA synthetase. We thus deployed this to simulate the effects of Gln4 depletion in yeast. One of the main outputs of the Global Translation Model is the average protein production rate, (essentially, the ribosomal ‘current’ along the mRNA), which can be used as a proxy for the growth rate of the cell (64)(65). Our model correctly predicts a decrease in the global current/protein production rate as the number of Gln4p molecules is decreased in response to doxycycline (Fig. 5D, blue solid line), in accordance with the experimental results in (Fig. 1D). The model also reproduces the plateau obtained in the growth rate as the number of Gln4p molecules is overexpressed. Note that the relatively large error bars arise from the large variability of the protein production rate among different GO-Slim categories (see Fig. 5I).

**Figure 5:**
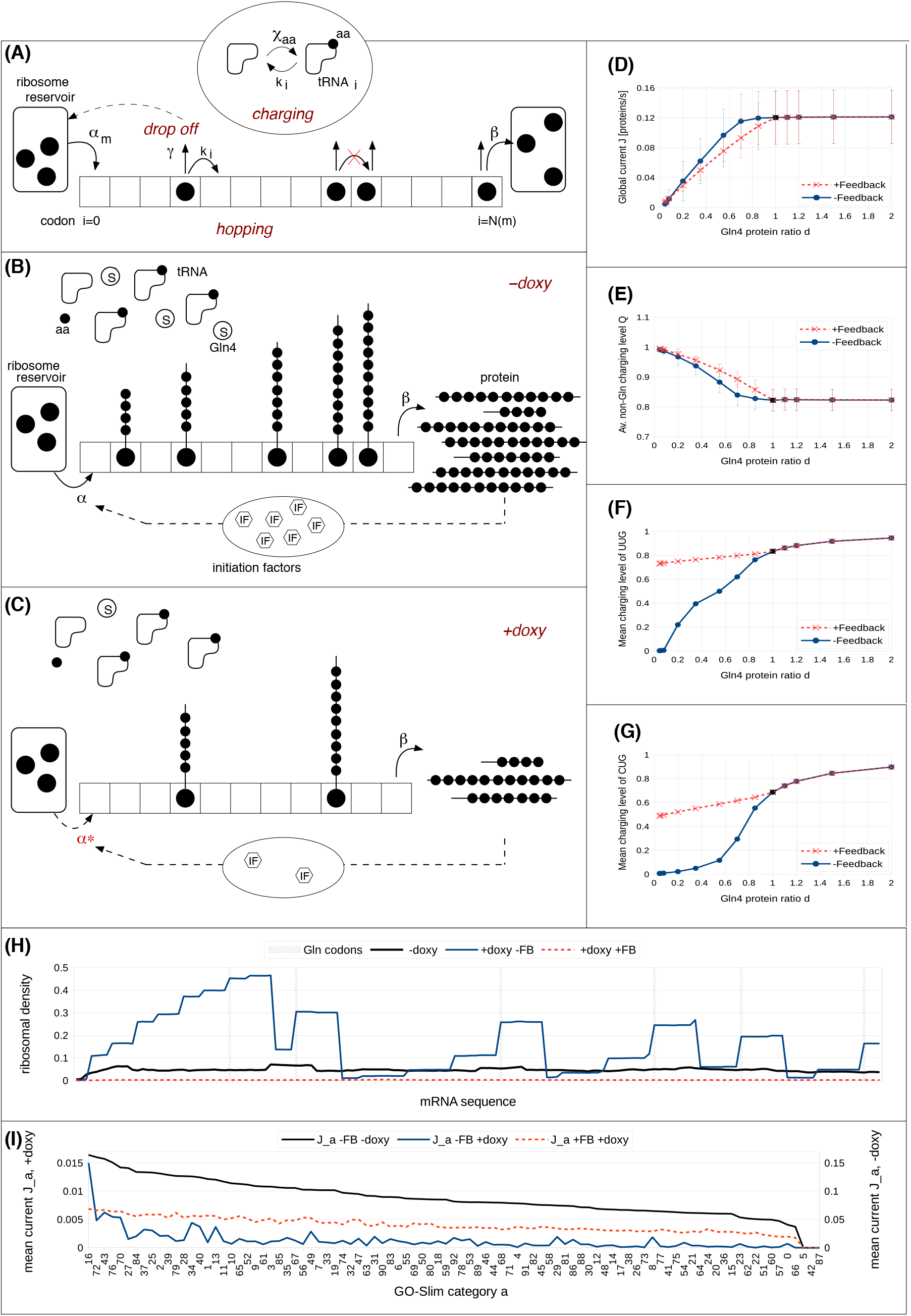
Simulation of a comprehensive translation system comprising tRNAs, ribosomes and codon-biased mRNAs to analyse tRNA synthetase depletion. *Panel A:* Global translation model of mRNAs by ribosomes including competition for tRNAs, tRNA synthetase and ribosomes. The ribosome from the reservoir enters the mRNA-lattice *m* via the initiation rate *α_m_* at the first lattice site, codon *ι* = 0, and terminates with the constant rate *β* at the last lattice site, codon *N(m).* On the lattice, the ribosome translates the codon *ι* with the hopping rate *k(í*) and may unbind from the lattice with the drop off rate *γ*. If the next codon is occupied, translation is not allowed. Note, that the ribosome has a footprint *W* = 9, i.e., the next codon is codon *i* + W (not shown here). Translating a lattice site uses a charged tRNA which then will be recharged via the charging rate *χ_aa_* according to the corresponding amino acid (aa). *Panels B, C:* Link between doxycycline and tRNA charging with and without feedback: Conceptual idea of the feedback loop, model without (B) and with (C) doxycycline: Due to the feedback loop, the initiation rate is reduced proportional to the rate of protein production. In the presence of doxycycline, the number of glutamine synthetases (denoted by S) decreases. As a consequence, the ribosomal speed along the mRNA decreases and less proteins are produced per unit time, and so, less initiation factors. This in turn decreases the initiation rate. Hence, the ribosomal current also decreases, thereby decreasing the demand for glutamine synthetases. This is why in the presence of doxycycline the charging level of glutamine in (F) and (G) is almost the same as in the case without doxycycline. *Panels D-G:* Doxycycline regulation of global protein synthesis and thus growth rate: The global current *J*(D), the average non-glutamine tRNA charging level 〈*Q*〉(E) and the mean charging level *Q* of the two glutamine tRNAs (F) UUG and (G) CUG with (red, dashed lines, crosses) and without (blue, solid lines, circles) feedback loop as functions of the Gln4 protein ratio *d*. Note, that *d* = 1 corresponds to zero doxycycline (black circles and crosses) and, e.g., *d* = 0.035 to a doxycycline concentration of O.5μg/ml. *Panel H:* Ribosomal density profiles along an mRNA from GO Slim category 16 (cytoplasmic translation), which has six glutamine codons, position is highlighted by the grey, dashed lines. The situation without doxycycline is represented by the black line. Note that it is the same for both models, with and without feedback, since the feedback is only effective in the presence of doxycycline. The blue line corresponds the situation with doxycycline, a Gln4 protein ratio of 5%, but without feedback. Here, ribosomal queuing behind the glutamine codons and a highly inhomogeneous profile is visible. In contrast, when the autogeneous feedback mechanism is turned on (red, dashed line) ribosomes are equally distributed in low density all over the mRNA, no queueing visible. For more details of the latter, see Supplementary materials. *Panel I:* The mean current *J_a_* in each GO-Slim category assorted descending (from category 16 to category 66) for the situation without doxycycline (black line). In the presence of doxycycline, the blue line corresponds to the situation without feedback mechanism and the red, dashed line represents the current with feedback loop. Note that the non-doxycycline current is plotted on the secondary axis due to an order of magnitude difference compared to the situation with doxycycline. Note, that the categories 5, 42 and 87 (see Table S2) are not represented within our downscaled system. The category key can be found in Table S11.

As the glutamine tRNA synthetase Gln4p is depleted from the cell, as expected the model predicts that charging levels of both tRNA^UUG^_Gln_ and tRNA^CUG^_Gln_ markedly decrease (Fig. 5F, G, blue solid line). As a consequence, the translation (hopping) rates associated with the corresponding CAA and CAG codons reduce and ribosomal queues form behind those codons (Fig. 5H). The reduced flow of ribosomes along mRNAs causes a global decrease in the protein production rate. Note that as we overexpress Gln4 the charging levels of the glutamine tRNAs increase, since the charging rate increases with the synthetase concentration.

Interestingly, the charging level of all other non-glutamine tRNAs increases to 100% as Gln4p is completely depleted from the system (Fig. 5E; blue solid lines). As ribosomes stall and queues develop, the demand for non-glutamine tRNA decreases dramatically, whereas their supply rate, defined by the corresponding values of *V_max_*, remains constant. As a consequence, their charging levels increase up to 100%. Hence, the model correctly predicts a reduction in the growth rate of the cell upon Gln4p depletion. However, although the model predicts ribosomal queues formation due to depletion of Gln-tRNA^Gln^, polysome analysis revealed no evidence of queues (Fig. 4B).

To resolve this conflict, we considered the fact that initiation rate decreases with decreasing growth rate. We hypothesised that the reduction in the global protein production rate directly feeds back on the initiation factor and ribosome production (66). We therefore integrated a homeostatic negative feedback loop into the model: as the rate of glutamine tRNA charging slows due to Gln4 depletion, so does the initiation rate, and fewer ribosomes enter the mRNAs. Note that in principle, the influence on the translation initiation rate is non-linear due to accumulative effects, but our analysis has shown that the most important influence seems to be the reduction in the initiation factors availability (see Supplementary Materials for a detailed discussion). To implement this negative feedback loop into the model we assumed that the initiation rate constant α_0_ decreases proportional to the growth rate. Therefore, we choose a linear decrease with the amount of Gln4p in the cell. In particular, we used 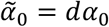, where 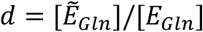 is the Gln4 protein ratio and 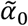 denotes the initiation rate constant in the presence of doxycycline.

To provide experimental evidence to support the introduction of this homeostatic feedback loop in the model as above we confirmed first that the reduction in growth rate accompanying tRNA synthetase depletion causes a concomitant reduction in translation initiation factor expression. Analysis showed that of 24 translation initiation factor genes, 21 were significantly transcriptionally repressed in response to doxycycline treatment in the *GLN4* tet-off strain (Table S9). We also confirmed that the reduction in growth rate accompanying tRNA synthetase depletion causes a concomitant reduction in ribosome content. We created a strain of yeast in which ribosomal protein uL23 (also referred to as RPL25) is tagged with GFP. This allowed us to accurately quantify the numbers of ribosomes per cell based on quantitation of fluorescence in a cell lysate. We grew this strain across a range of doxycycline conditions and measured ribosome content. In the absence of doxycycline, there were 250,000 ribosomes per cell, in excellent agreement with published values (66). This reduced in a dose-responsive manner as doxycycline reduced the cellular content of Gln tRNA synthetase, and thus the growth rate (Fig. 6). We also qualitatively measured GFP fluorescence per cell using cell cytometry, confirming the same doxycycline-dependent decrease (Fig. 6). We thus confirmed that the reduced growth rate caused by doxycycline-shut-off of Gln4p is indeed associated with reduced cellular ribosome content, as well as a reduction in expression of 21 of the 24 translation initiation factors.

**Figure 6:**
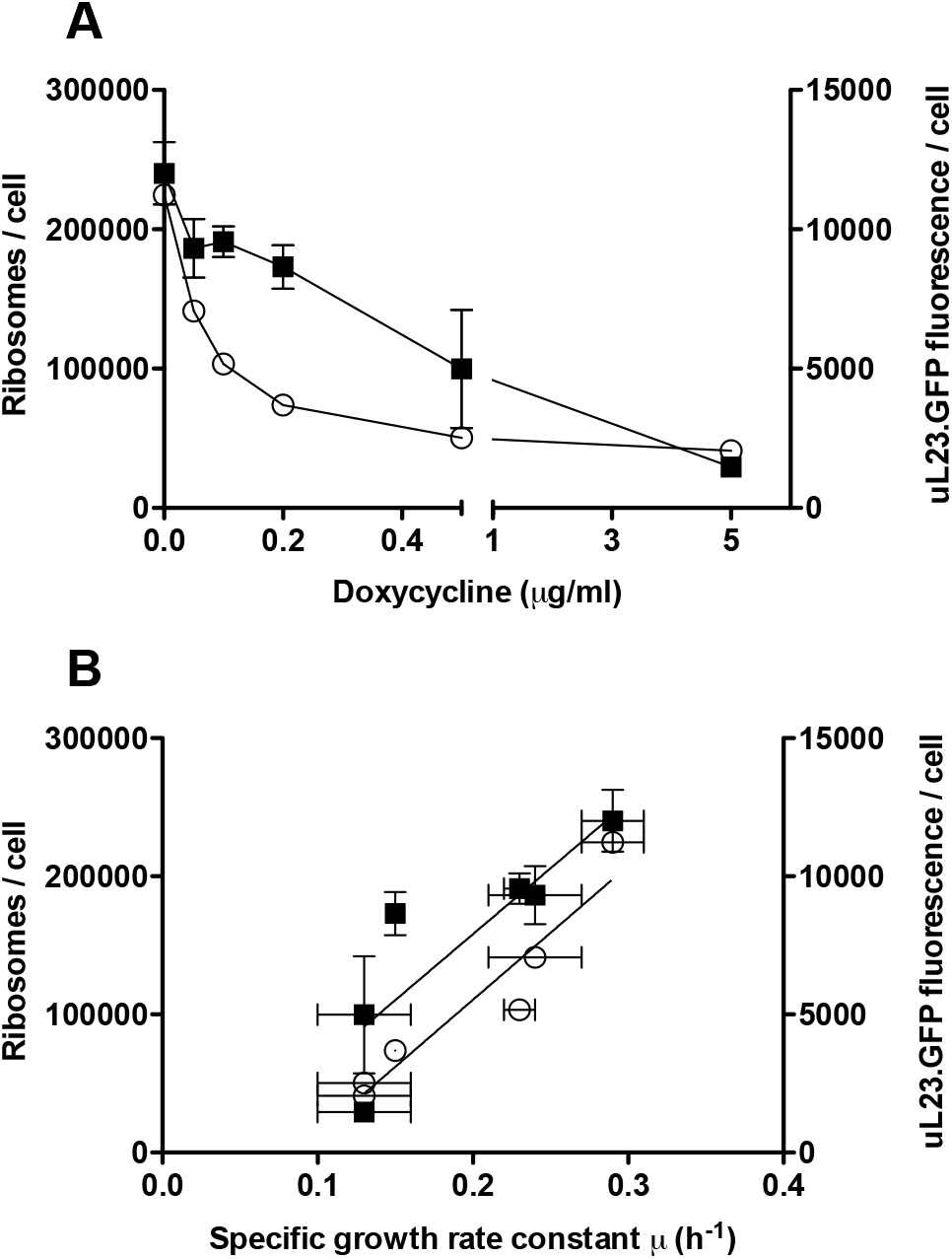
Cellular ribosome content reduces with growth rate as tRNA synthetase is depleted. The cellular content of ribosomes in the tet-off, Gln-tRNA synthetase strain grown to mid-log phase was calculated using fluorescence quantification of GFP-tagged uL23 (RPL25) in cell lysates across a range of doxycycline concentrations (closed squares represent mean ribosome content, n=3, ± standard deviation). In parallel, the geometric mean fluorescence per cell was quantitated in arbitrary fluorescence units per cell using flow cytometry across the same range of doxycycline concentrations and growth rates in 15,000 cells (open circles represent geometric mean ribosome content, n=3, ± standard deviation [horizontal error bars] for the growth rate determination). Results are plotted against doxycycline concentrations (panel A) and the specific growth rate constant (panel B).

Crucially, with the incorporation of the negative feedback loop, simulations of ribosome density profiles no longer exhibit ribosomal queues (Figs. 5H, and Fig. S4 including Table S10), in agreement with the polysome profile experiments. Now we obtain a quite homogeneous density profile (Fig. 5H; red dashed line). Note that ribosomal density on the lattice decreases strongly compared to the situation without doxycycline, with a smaller ribosomal pool causing reduced rates of translation initiation.

The average protein production rate still decreases as a function of the Gln4p abundance (Fig. 5D; red dashed line), in good agreement with the measurements (Fig. 1D). However, at first sight it seems counter-intuitive that the global current is very similar with and without feedback (Fig. 5D, compare blue and red lines), since with feedback the initiation rate is strongly reduced. The explanation is in part that without feedback, flow is reduced because ribosomes queue, and with feedback, flow is reduced because there are fewer ribosomes on the mRNA (including for CAG-rich mRNAs; see Figs. S5 and S6). However, there are also mRNA-specific effects. To understand this, we analysed how each GO-Slim category is affected (Fig. 5I). Here, the black line represents the mean current for each GO-Slim category without doxycycline, where the categories have been sorted in descending order according to their current (rate of translation). The blue and the red line correspond to the mean current for each category in the presence of doxycycline, without (blue line) and with (red line) the feedback mechanism.

Without the feedback, the proportion of each type of GO-Slim protein in the simulation is markedly affected. This can be seen by the curve not being strictly decreasing and having strong fluctuations. The different categories respond differently due to CAG codon content. For example, category 16 (cytoplasmic translation), comprised of highly codon biased mRNAs, has only 0.03% of CAG codons equating to 0.04 CAG codons per mRNA on average. Consequently, its current is the least affected by doxycycline. In contrast, category 66 (regulation transport, with poor codon bias) has 0.9% of CAG codons, corresponding to 8.6 codons per mRNA on average and is the most affected by doxycycline (Supplementary Materials). With the negative feedback loop now incorporated, the curve (Fig. 5I; red line) is much smoother, maintaining the descending order in the current almost perfectly. Therefore, the proportion of different types of proteins in the cell is maintained despite Gln4 depletion, and the inclusion of feedback restores the composition of the cell at the functional level.

Importantly however, the model incorporating feedback makes a clear, non-intuitive prediction: The charging levels of the glutamine tRNAs stay almost constant, decreasing only slightly as Gln4p is depleted (Fig. 5F, G). This is in contrast to the prediction made by the model without feedback. Intuitively, this is because with feedback, the initiation rate decreases as we deplete the system of Gln4p. As a consequence, the current of ribosomes along mRNAs decreases, as does the demand for glutamine tRNAs. The charged level of glutamine tRNAs thus only decreases slightly with doxycycline treatment. Hence, the feedback mechanism homeostatically restores the balance between the rate at which glutamine tRNAs are charged and the rate at which they are used.

This slight decrease in the charging level of glutamine tRNAs is enough to cause an increase in the charging level of non-glutamine tRNAs: As we have seen, the decrease in the Gln-synthetase levels causes a drop of global translation rate. As a consequence, the demand for non-Gln tRNAs also decreases. However, the fact that the *V_max_* of non-Gln synthetases are unaffected by doxycycline leads to an increase in the charged levels of non-Gln tRNAs. Note that the feedback mechanism is only effective in the presence of doxycyline, i.e. as the number of Gln4p molecules is overexpressed (i.e, in the absence of doxycycline), the two models coincide, see (Fig. 5D-G).

In summary, the simulation results including the negative feedback loop are consistent with the lack of ribosomal queuing observed in the polysome analysis (Fig. 4), as well as the impact that depletion of Gln4p has on the growth rate. The model also makes a clear, non-intuitive prediction, namely that the charging level of the glutamine tRNAs remains approximately constant across a range of doxycycline concentrations, since a fewer number of ribosomes start translation, thereby limiting the demand for charged glutaminyl tRNAs. All other tRNAs become over-charged due to an excess charging capacity. We therefore set out to validate this prediction experimentally, below.

### The proportion of glutamine-charged tRNAs is not affected by glutamine synthetase depletion

The mathematical model makes the clear, non-intuitive prediction that the charging level of a given tRNA stays almost constant as its tRNA synthetase is depleted from the system, whereas the charging level of all the other tRNAs increases. To validate these predictions, tRNAs were extracted either from *tetO-GLN4* (glutaminyl-tRNA synthetase shut-off) or *tetO-KRS1* (lysyl-tRNA synthetase shut-off) cells grown in a range of doxycycline concentrations. tRNAs were resolved on denaturing polyacrylamide gel, blotted and probed with biotin-labelled oligonucleotides specific for either the tRNA whose cognate synthetase is being depleted, or a control tRNA not directly affected by this depletion.

In the case of the lysyl-tRNA synthetase shut-off, the results clearly show that as the concentration of doxycycline increases, as the model predicts the proportion of charged lysine tRNA does not reduce, even increasing somewhat (Fig. 7A, B). The control, arginyl-tRNA charging levels also remained constant across a range of doxycycline concentrations and decreasing growth rates (Fig. 7A, B). In the case of the glutaminyl tRNA synthetase shut-off, there was also no reduction in the ratio of charged to un-charged tRNA in response to doxycycline treatment (Fig. 7C, D). In parallel, the control tRNA, in this case lysyl-tRNA, also did not reduce in response to doxycycline, and gradually increased as the model predicts (Fig. 5E).

**Figure 7:**
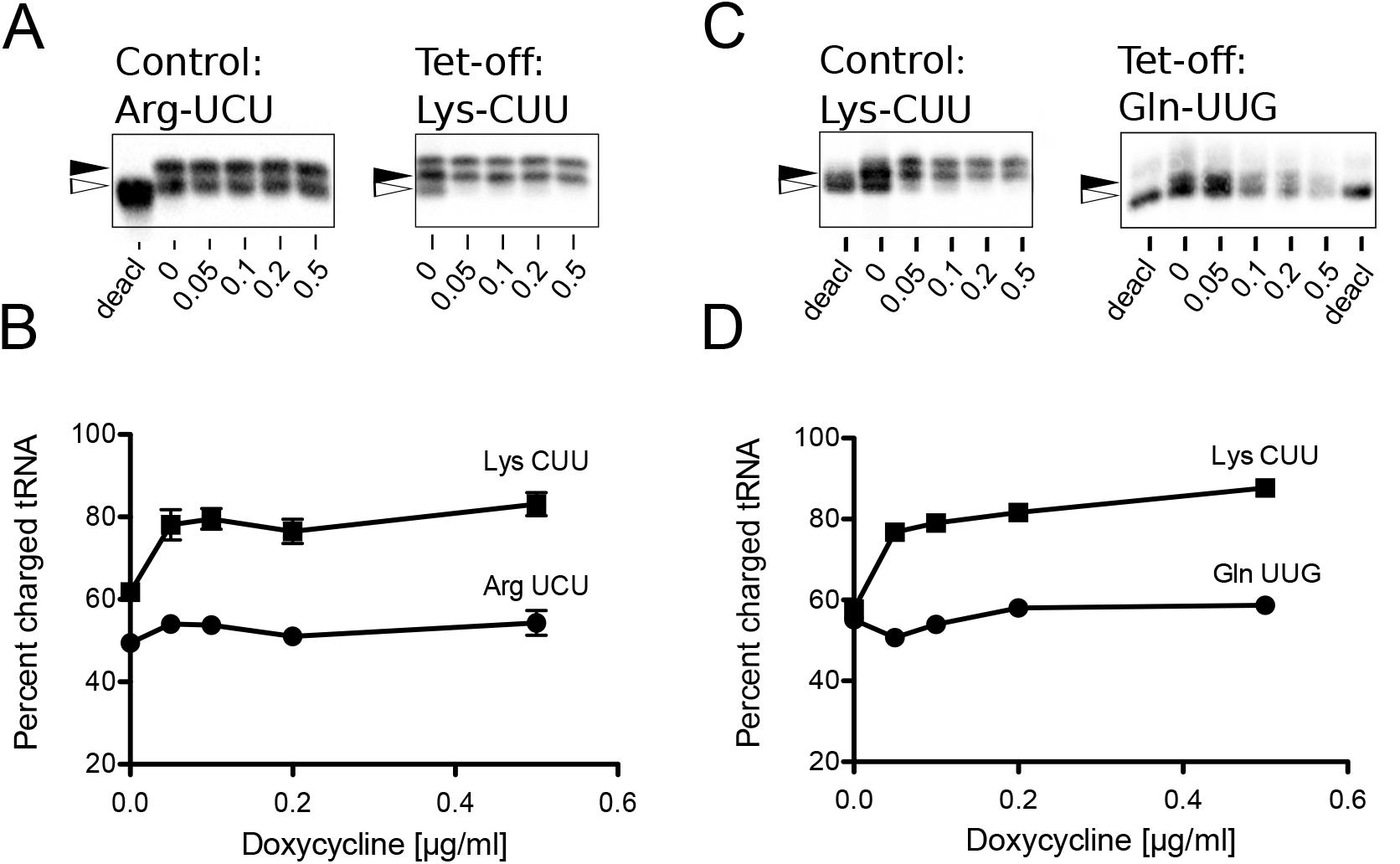
The proportion of charged tRNAs is not reduced by tRNA synthetase depletion. *Panel A:* Northern blots of control tRNA^Arg^_UCU_ and tet-off tRNA^Lys^_CUU_ isolated from a *tetO-KRS1* (lysyl – tRNA synthetase shut-off) strain grown in a range of doxycycline concentrations up to 0.5 μg ml^-1^. Blots include a control lane of pH 8-treated tRNA to provide a marker of deacylated tRNA (open arrow, ‘deacl’). The slower migration position of acylated tRNA is indicated (closed arrowhead). Results shown are typical of other biologically independent experimental replicates. There is a linear relationship between signal and image intensity. The images shown are of single blots and are not composite (2μg total RNAs loaded per lane). *Panel B:* Quantification of the Northern blot showing average % tRNA charging (UCU-Arg; closed circles) and CUU-Lys (closed squares). Means of three biological replicates on independent blots ± standard deviation (n=3). *Panel C:* Northern blots of control tRNA^Lys^_CUU_ and tet-off tRNA^Gln^_UUG_ isolated from a tetO-*GLN4* (glutaminyl-tRNA synthetase shut-off) strain grown in a range of doxycycline concentrations, and band intensities quantified (*panel D;* UUG-Gln; closed circles and CUU-Lys; closed squares; 2μg total RNAs loaded per lane).

The experimental result therefore validates the model predictions, and explains that as the activity of any individual tRNA synthetase decreases, the charging level of its target tRNA is paradoxically preserved, because the rate of translation initiation decreases as growth rate (and therefore ribosome content) also reduces. The translational demand for charged glutamine tRNAs is therefore matched with the capacity of the synthetase to supply them. Moreover, there is some experimental evidence that the charging level of the other tRNAs may increase as a consequence, as predicted by the model.

### Model simulation of uncharged tRNA sequestration by the tRNA synthetase network

The model prediction that the levels of uncharged glutamine tRNA change only slightly with increasing doxycycline is validated by the tRNA Northern blot experimental observation (Figs 5 and 7). These results are however apparently at odds with the earlier observations that treating the *tetO-GLN4* cells with doxycycline induces a *GCN4* amino acid starvation response, indicative of the accumulation of uncharged tRNA. To address this paradox, we noted two facts; (i) that in a cell undergoing active translation, a proportion of the uncharged tRNA existing at any moment in time is bound to the tRNA synthetase as the tRNA undergoes catalytic conversion, and (ii) there are thought to be approximately 20,000 Gln4p molecules per cell (67)(58). It is significant that there are approximately 100,000 tRNA^Gln^ in the cell (66)(68), of which measurements indicate 20% (i.e. 20,000) are uncharged in an actively growing cell. This suggests a hypothesis that under normal physiological conditions, some proportion of the uncharged tRNA population may be effectively sequestered by the tRNA synthetase.

We sought to test this hypothesis using a simple model (Fig. 8A), which assumes that uncharged tRNA population is divided between a *bound* state (*b*), where the tRNA is uncharged but bound to the corresponding synthetase enzyme and an *empty* (uncharged and unbound) state (*e*). The third state in figure 8A is the *charged* state (*c*) where the tRNA is charged with its corresponding amino acid. The total number 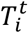 of tRNAs of type *i*. stays constant 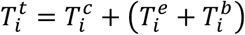 and the number of uncharged tRNA is then given by 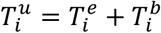.

**Figure 8.**
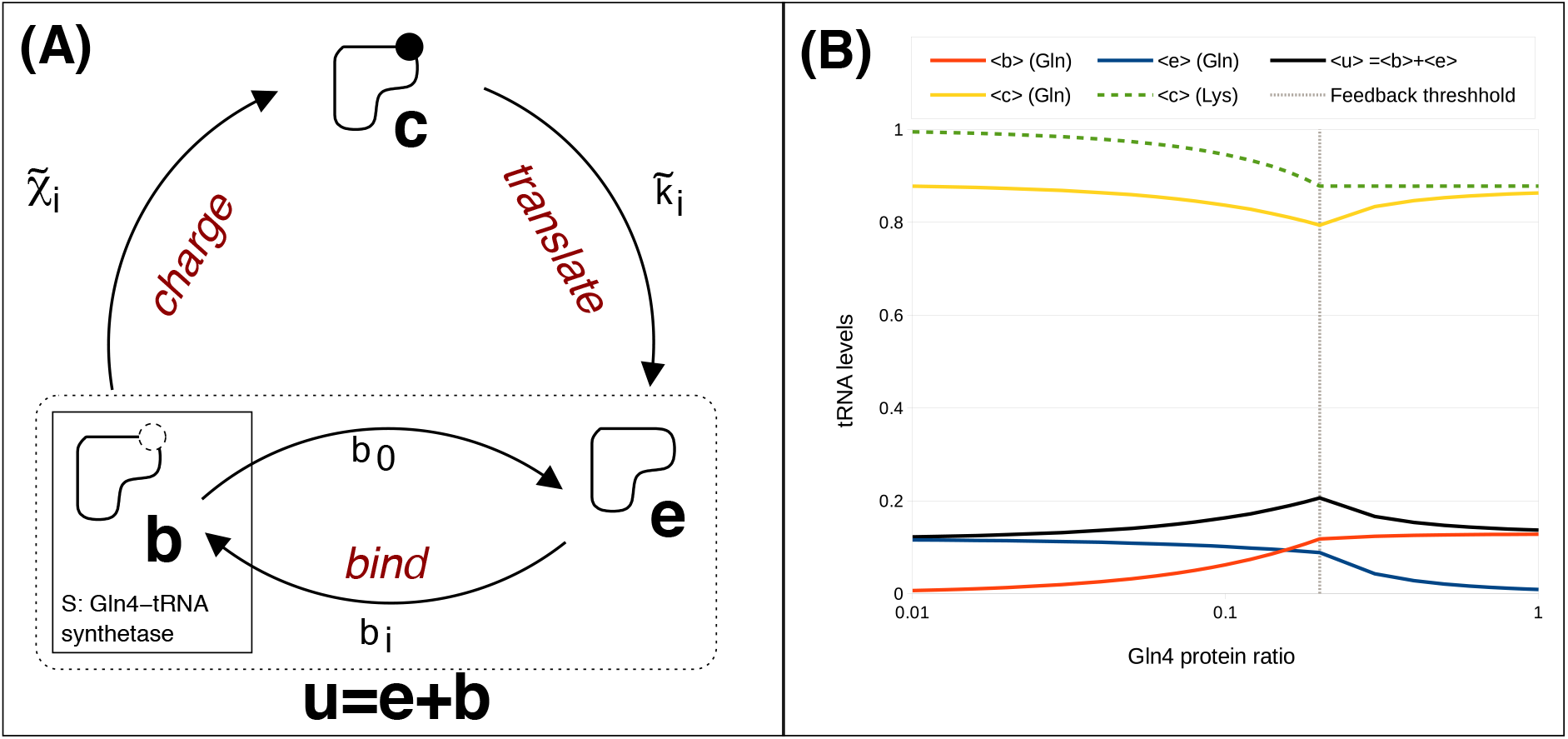
Analysis of the levels of free tRNA in a synthetase-depleted strain using model simulation. *Panel A;* Synthetase sequestration model for tRNAs: When we measure uncharged tRNA, some of it is synthetase-associated, captured mid-reaction, but still uncharged. Hence, two intermediate states are introduced: the “bound” state (*b*), and the “empty” state (*e);* the tRNA with amino acid is in the “charged” state (*c*). The transition from the empty state to the bound state, during which the synthetase associates with the tRNA at the binding rate *b_i_*, depends on the synthetase concentration [*E_i_*], and the number of “empty” tRNAs 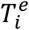 of amino acid of type *i*. The reverse transition *b*_0_describes the dissociation of the synthetase-tRNA complex and therefore depends on the number 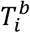 of “bound” tRNAs. The transition from “bound”(b) to “charged” (c) state occurs at rate 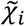, and it is affiliated with the catalytic rate *k_cat,i_* and it also depends on the number 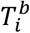 of “bound” tRNAs. The transition between the “charged” (c) and the “empty” (e) state is governed by the doxycycline sensitive hopping rate 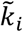, incorporating the feedback, and it depends on the number 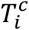 of “charged” tRNAs. *Panel B;* Resulting plot of the steady state charging levels of the glutamine tRNAs as a function of the Gln4 protein ratio; using the Gillespie algorithm, the 3-state-network of all 20 amino acids was simulated for different doxycycline factors. Yellow line: mean level 〈*c*〉 of charged glutamine tRNAs; Red line: mean level b) of bound glutamine tRNAs; Blue line: mean level 〈*e*〉 of empty glutamine tRNAs; Black line: mean level 〈*u*〉 = 〈*e*〉 + 〈*b*〉 of uncharged glutamine tRNAs; Green dashed line: mean level of charged lysine tRNAs. The grey dashed line indicates the feedback threshold *d* = 0.2: for Gln4 protein ratios smaller than 0.2 the usage rate decreases analogously to the growth rate.

The Synthetase Sequestration Model (SSM) considers explicitly two main steps in the process of aminoacylation: first, the tRNA is bound by the synthetase, and in a second step, the amino acid is attached to the tRNA. We refer to these two different states as to *bound* and *charged.* In the bound state, the tRNA is bound to the synthetase but uncharged, i.e., the tRNA is *sequestered* by the synthetase. Our model predicts how the balance between the three different tRNA states (*empty, bound* and *charged*) changes depending on Gln4p availability.

In the bound state, tRNA is not available to react with Gcn2 kinase, which detects uncharged tRNA and activates the *GCN4* amino acid starvation response (69). Therefore, when the synthetase concentration drops in response to doxycycline, there are fewer synthetase molecules available to bind to the tRNA, so that 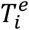 increases at the expense of 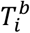. Crucially, the increased amount of *empty* (*e*) tRNA is now available to react with Gcn2, and trigger the *GCN4* response.

The results show that both the charged and uncharged levels of the glutamine tRNAs change only slightly, but the composition of the uncharged glutamine tRNAs changes strongly: without doxycycline, *d* = 1, the simulation reveals that almost all uncharged tRNAs are in the bound state (Fig. 8). In the presence of doxycycline, due to the reduced Gln4 synthetase concentration, the bound level lessens in favour of the empty level. Moreover, the mean level of charged lysine tRNAs increases to 100% as the Gln4 protein ratio decreases. The lysine tRNAs are representative for all non-glutamine tRNAs.

We assume that the transition between the empty and the bound state occurs at rate 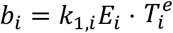, where *k*_1,*i*_ is the rate constant and the number of free synthetase molecules is given by 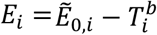, where 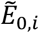 is the total amount of synthetase molecules of this type and 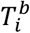 is the number of synthetase-tRNA complexes. Here, doxycycline reduces only the Gln4 protein, and the Gln4 protein ratio is denoted by 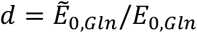, where 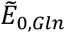 and 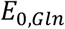 denote the total number of Gln4p in the presence and absence of doxycycline, respectively. Hence, 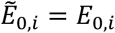 for all other amino acids. The reverse transition 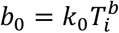 describes the dissociation of the synthetase-tRNA complex, with the rate constant *k*_0_.

The transition between the bound and the charged state is governed by the charging rate 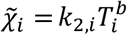, where the rate constants *k*_2,*i*_ are proportional to the catalytic rates *k_cat,i_* used in the Global Translation Model.

During the transition between the charged and empty state, the amino acid is transferred to the nascent polypeptide. Therefore, the transition rate between the charged and empty state is given by ribosomal current (translation rate). This can also be understood as the tRNA usage rate, and analogous to the Global Translation Model, we assume that this transition rate is given by 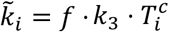, with the rate constant *k*_3_ and where *f* is a coupling feedback factor by which the usage rate of all tRNAs decreases as a function of the Gln4 concentration. Based on the experimental data presented in Fig. 1D on how the growth rate depends on the glutamine synthetase ratio, we define a stepwise linear function with *f* = 5*d* for *d* < 0.2 and *f* = 1 otherwise, where *d* denotes the glutamine synthetase ratio, as before. This reflects the fact that if the growth rate is not affected by doxycycline, then the charged tRNA usage rate should not be affected either. However, as the growth rate decreases, the initiation rate also decreases, thereby lowering the charged tRNA usage rate. The resulting synthetase sequestration model in figure 8A (simulated using a multiple-copy 3-state network; Supplementary Information) considers a charging cycle for each amino acid and a total number of tRNAs in multiples of the corresponding gene copy number.

Interestingly, due to the coupling feedback factor, this relatively simple model reproduces the predicted charging level of non-glutamine tRNAs in the Global Translation Model: the charging level (lysine is shown) increases in the presence of doxycycline (Fig. 8B). The coupling feedback factor reflects the decreasing growth rate, and translation rate of the cell in the presence of doxycycline. However, doxycycline only reduces Gln4p charging rate, therefore charging of non-glutamine amino acids then becomes the dominating form of transition in the process.

Importantly, the synthetase sequestration model predicts that in the absence of doxycycline, the greater proportion of uncharged tRNA is located in the *bound* state (*b*) (Fig. 8B). Adding doxycycline then increases the number of *empty* tRNAs in state (*e*) at the expense of tRNAs in the bound state (*b*). This result suggests that with the reduction in population of glutamine tRNA synthetases, although the total amount of uncharged tRNA remains more or less unchanged (as shown experimentally in Figure 7), doxycycline depletes Gln4p, causing a reduced capacity to sequester uncharged tRNA. The ratio of (*b*):(*e*) tRNAs is shifted in favour of a higher proportion of *empty* (*e*) tRNA, which can activate the Gcn2 kinase. This accounts for an activation of *GCN4* even while the proportion of uncharged tRNA is relatively stable.

## DISCUSSION

tRNA synthetases play a central role in the translation process, maintaining a supply of aminoacylated tRNAs sufficient to supply translation elongation. Underscoring their importance, there is a growing list of human genetic diseases linked to human tRNA synthetase genes. These cause a range of neurological and neurodevelopmental defects (70)(71)(16). To better understand the molecular aetiology of such disorders, we established a doxycycline-dependent glutamine tRNA synthetase shut-off system in which levels of tRNA synthetase could be controlled by doxycycline in a dose-dependent manner.

Our initial characterisation of synthetase depletion effects using quantitative Western blotting confirmed reductions in growth rate when the Gln4p cellular content drops below approximately 20,000 copies per cell, in close agreement to estimates for the physiological Gln4p abundance in yeast of 17,000 molecules per cell (58)(59). This indicates a tuned maintenance of tRNA synthetase levels in the cell sufficient to support the translational demands of the elongating ribosome population.

Intuitively, during active growth, a restriction in the charging capacity of tRNA synthetase might be expected to generate increased levels of uncharged tRNA, which activate Gcn2 kinase. Gcn2 can also be activated in a deacylated tRNA-independent manner in mouse (72) although it is not certain this happens in yeast. Gcn2 activation would act as a signal of amino acid starvation and, via phosphorylation of translation factor eIF2α, induce translation of *GCN4* transcription factor (62). This chain of events forms part of the so-called Integrated Stress Response (ISR), a series of sensors and responses that focus on regulation of protein synthesis in response to a range of imposed stresses (73). In fact, for both Gln- and Lys-tRNA synthetase shut-off events, our studies revealed just such an effect in response to doxycycline, with induction of a *lacZ* reporter gene under *GCN4* translational control (Fig. 2 and S7), and a cell-wide transcriptional response highly similar to a *GCN4* transcriptional response (66; Fig. 3). We considered additionally that reduced levels of charged tRNA might cause mistranslation, but tRNA synthetase depletion did not cause enhanced sensitivity to the translation error-inducing drug hygromycin B (Fig. S7).

Although increased levels of uncharged glutamine tRNAs should cause ribosomal queuing at CAA and CAG glutamine codons, in fact there was no evidence of shifts to larger polysomes in response to doxycycline (Fig. 4). We reasoned that the formation of queues may be limited by the *GCN4* response, which slows translation initiation via eIF2α phosphorylation, limiting the feed of ribosomes onto the transcriptome. However, ablating the *GCN2* kinase gene during doxycycline restriction of Gln4 did not reveal any shift to larger polysomes (Fig. 4). In order to understand why this might be so, we applied mathematical modelling to simulate the restriction in supply of charged tRNA to translation. The global model of translation used is to the best of our knowledge the first model that tracks the charged status of all tRNAs as they cycle through translation, become deacylated, and are then recharged by tRNA synthetases. The model tracks a fixed population of ribosomes as they initiate, elongate and terminate on a population of mRNAs with codon compositions matching those of the GO-Slim gene ontologies. It thus provides a fine-grained view of how translation responds to changes in the cellular Gln4p levels.

The model confirmed a series of experimental observations. First, simulating a reduction in the concentration of Gln4p produced slowed rates of translation (indicated in the model as a reduced current of ribosomes; Fig. 5D), matching experimental observation (Fig. 1). Slow growth will drive slower rates of initiation, via mass action principles. This was implemented in the model by the introduction of feedback; reductions in Gln4 levels drive reduced rates of translation initiation. In fact, this effect would also have been exacerbated by the *GCN4* response (Fig. 2, 3), through Gcn2 kinase phosphorylation of eIF2α. Simulations including the feedback effect mirrored experimental observation, namely that doxycycline restriction of Gln4 causes a global slow-down of translation initiation that prevents ribosomal queue formation (Fig. 5H). Essentially, mRNA translation becomes initiation limited (76) because of the limitation in tRNA charging capacity. Crucially however, the model made two testable predictions; first, that non-glutamine tRNA populations should become fully charged with amino acid during doxycycline treatment. Essentially, the entire translation system is regulated by the availability of Gln-tRNA^Gln^, and other tRNAs are utilised at slower rates, creating excess capacity in the population of 19, non-Gln synthetase species. The second prediction was that paradoxically, there should be only minor reductions in the levels of charged glutamine tRNAs, despite our detection of a *GCN4* response in the presence of doxycycline. Nevertheless, Northern blot analysis of tRNA charging confirmed both predictions (Fig. 7); the tRNA^Lys^ population became increasingly charged in a doxycycline-responsive manner during Gln tRNA synthetase shutoff, while the levels of charged glutamine tRNA were essentially unaltered across a range of doxycycline concentrations. The maintenance of tRNA charging, despite synthetase depletion was also seen the in case of the lysyl-tRNA in the second lysyl-tRNA synthetase shut-off strain, validating the model predictions.

To address the intriguing, apparent contradiction of a *GCN4* starvation response but with unaltered levels of tRNA^Gln^ charging, a minimal *synthetase sequestration* model simulated the effects of reducing cellular Gln4p synthetase, while levels of charged, and uncharged tRNA were monitored. Crucially, the latter species was further defined in terms of uncharged tRNA that is synthetase-bound (*b*) and that which is unbound, ‘empty’ (*e*) tRNA (Fig. 8). The model clearly showed that under normal physiological conditions of active translation, with 80% typical charging levels of tRNA, the 20% uncharged tRNA population is held in the bound state by the synthetase population, masking it from the Gcn2 kinase. However, in the presence of doxycycline, levels of Gln4p drop, and the ability of the Gln4p population to sequester uncharged tRNA during the charging process is diminished. Levels of unbound *‘e’* tRNA rise, allowing Gcn2 activation. This explains the clear signature of the *GCN4* response in response to doxycycline while tRNA charging levels are maintained (Fig. 2, 3).

Our study makes clear predictions concerning the molecular aetiology of the human tRNA synthetase gene mutations which cause a wide range of neurological, and neurodevelopmental defects (16). The model predicts potentially distinct effects on accumulation of unbound tRNA depending on whether a mutation alters the catalytic rate constant (*k_cat_*) of the tRNA synthetase, the affinity (*K_M_*) of the synthetase for tRNA or its expression level. The latter mimics the doxycycline shut-off model implemented in our study, which triggers a *GCN4* response. For example, valine synthetase (VARS) mutations that cause microcephaly and seizures can cause both loss of charging function, and exhibit much reduced expression (71). These would be predicted to trigger the ISR via Gcn2 activation due to a limited ability to sequester the cognate uncharged tRNA. Supporting the link between the ISR and neurological defects, mutations in translation initiation factor eIF2B, also cause a range of neurological defects including vanishing white matter (VWM) (74)(75). Some of these mutations compromise eIF2B’s guanine exchange factor (GEF) activity for eIF2, and are thus predicted to reduce the threshold for triggering the ISR/*GCN4* response.

Similar processes may operate in some types of tRNA synthetase mutation, where the synthetase sequestration model indicates unbound, uncharged tRNA is available to trigger a *GCN4* response. Overall the combined experimental and model simulation analysis explains how levels of charged tRNA are maintained despite the reductions in a given tRNA synthetase, because slowed growth leads to reductions in ribosome content and translational activity; this homeostatically matches demand for charged tRNA to its supply. However, as the synthetase sequestration model presented makes clear, depletion of Gln tRNA synthetase limits its capacity to sequester tRNA during the charging reaction. Thus, a greater proportion of the uncharged tRNA is available to react with Gcn2 kinase, triggering an amino acid starvation *GCN4* response. Our study reveals fundamental new insight into the role of the tRNA synthetase population in sequestering uncharged tRNAs, thus limiting the ability of the uncharged tRNA population to interact with the Gcn2 kinase, central to triggering a *GCN4* response. The study further identifies how the biochemical properties conferred by any given tRNA synthetase mutant may confer distinct molecular and thus disease phenotypes in human.

## ACKNOWLEDGEMENTS

The authors gratefully acknowledge the RNA sequencing and mass spectrometry support provided by the university of Aberdeen’s Centre for Genome Enabled Biology and Medicine, and Proteomics respectively. The Global Translation Model is deposited in the Biomodels database (76) with reference MODEL2001080004. MMcF carried out strain construction and characterisation, *GCN4* assays, RNA sequencing and mass spectrometry data analysis. IS conducted *GCN4* assays. HC carried out vector and strain construction and tRNA charging assays. Polysome profiling analysis was carried out by BC. SA carried out the ribosome quantification experiments. AR and MCR coded the original global translation model. CK and MCR extended the global translation model, CK coded and analysed the synthetase sequestration model, carried out the global translation model simulations and analysis. MCR and IS conceived the study and guided the research. IS, MCR, CK and MMcF co-wrote the manuscript.

## REFERENCES

1. Warner,J.R. (1999) The economics of ribosome biosynthesis in yeast. Trends Biochem. Sci., 24, 437–440.

2. Rodnina,M. V and Wintermeyer,W. (2009) Recent mechanistic insights into eukaryotic ribosomes. Curr. Opin. Cell Biol., 21, 435–443.

3. Rajendran,V., Kalita,P., Shukla,H., Kumar,A. and Tripathi,T. (2018) Aminoacyl-tRNA synthetases: Structure, function, and drug discovery. Int. J. Biol. Macromol., 111, 400–414.

4. Hinnebusch,A.G. (2005) Translational regulation of *GCN4* and the general amino acid control of yeast. Annu. Rev. Microbiol., 59, 407–450.

5. McNulty,D.E., Claffee,B.A., Huddleston,M.J. and Kane,J.F. (2003) Mistranslational errors associated with the rare arginine codon CGG in *Escherichia coli*. Protein Expr. Purif., 27, 365–374.

6. Belcourt,M.F. and Farabaugh,P.J. (1990) Ribosomal frameshifting in the yeast retrotransposon Ty: tRNAs induce slippage on a 7 nucleotide minimal site. Cell, 62, 339–352.

7. Kawakami,K., Pande,S., Faiola,B., Moore,D.P., Boeke,J.D., Farabaugh,P.J., Strathern,J.N., Nakamura,Y. and Garfinkel,D.J. (1993) A rare tRNA-Arg(CCU) that regulates Ty1 element ribosomal frameshifting is essential for Ty1 retrotransposition in *Saccharomyces cerevisiae*. Genetics, 135, 309–320.

8. Roche,E.D. and Sauer,R.T. (1999) SsrA-mediated peptide tagging caused by rare codons and tRNA scarcity. EMBO J., 18, 4579–4589.

9. Doma,M.K. and Parker,R. (2006) Endonucleolytic cleavage of eukaryotic mRNAs with stalls in translation elongation. Nature, 440, 561–564.

10. Matsuo,Y., Ikeuchi,K., Saeki,Y., Iwasaki,S., Schmidt,C., Udagawa,T., Sato,F., Tsuchiya,H., Becker,T., Tanaka,K., et al. (2017) Ubiquitination of stalled ribosome triggers ribosome-associated quality control. Nat. Commun., 8, 151–159.

11. Sitron,C.S., Park,J.H. and Brandman,O. (2017) Asc1, Hel2, and Slh1 couple translation arrest to nascent chain degradation. RNA, 23, 798–810.

12. Verma,R., Reichermeier,K.M., Burroughs,A.M., Oania,R.S., Reitsma,J.M., Aravind,L. and Deshaies,R.J. (2018) Vms1 and ANKZF1 peptidyl-tRNA hydrolases release nascent chains from stalled ribosomes. Nature, 557, 446–451.

13. Brandman,O., Stewart-Ornstein,J., Wong,D., Larson,A., Williams,C.C., Li,G.W., Zhou,S., King,D., Shen,P.S., Weibezahn,J., et al. (2012) A ribosome-bound quality control complex triggers degradation of nascent peptides and signals translation stress. Cell, 151, 1042–1054.

14. Defenouillere,Q., Zhang,E., Namane,A., Mouaikel,J., Jacquier,A. and Fromont-Racine,M. (2016) Rqc1 and Ltn1 Prevent C-terminal Alanine-Threonine Tail (CAT-tail)-induced Protein Aggregation by Efficient Recruitment of Cdc48 on Stalled 60S Subunits. J. Biol. Chem., 291, 12245–12253.

15. Shen,P.S., Park,J., Qin,Y., Li,X., Parsawar,K., Larson,M.H., Cox,J., Cheng,Y., Lambowitz,A.M., Weissman,J.S., et al. (2015) Protein synthesis. Rqc2p and 60S ribosomal subunits mediate mRNA-independent elongation of nascent chains. Science, 347, 75–78.

16. Antonellis,A. and Green,E.D. (2008) The role of aminoacyl-tRNA synthetases in genetic diseases. Annu. Rev. Genomics Hum. Genet., 9, 87–107.

17. Antonellis,A., Ellsworth,R.E., Sambuughin,N., Puls,I., Abel,A., Lee-Lin,S.Q., Jordanova,A., Kremensky,I., Christodoulou,K., Middleton,L.T., et al. (2003) Glycyl tRNA synthetase mutations in Charcot-Marie-Tooth disease type 2D and distal spinal muscular atrophy type V. Am. J. Hum. Genet., 72, 1293–1299.

18. Gonzalez,M., McLaughlin,H., Houlden,H., Guo,M., Yo-Tsen,L., Hadjivassilious,M., Speziani,F., Yang,X.L., Antonellis,A., Reilly,M.M., et al. (2013) Exome sequencing identifies a significant variant in methionyl-tRNA synthetase (MARS) in a family with late-onset CMT2. J. Neurol. Neurosurg. Psychiatry, 84, 1247–1249.

19. Jordanova,A., Irobi,J., Thomas,F.P., Dijck,P. Van, Meerschaert,K., Dewil,M., Dierick,I., Jacobs,A., Vriendt,E. De, Guergueltcheva,V., et al. (2006) Disrupted function and axonal distribution of mutant tyrosyl-tRNA synthetase in dominant intermediate Charcot-Marie-Tooth neuropathy. Nat. Genet., 38, 197–202.

20. McLaughlin,H.M., Sakaguchi,R., Liu,C., Igarashi,T., Pehlivan,D., Chu,K., Iyer,R., Cruz,P., Cherukuri,P.F., Hansen,N.F., et al. (2010) Compound heterozygosity for loss-of-function lysyl-tRNA synthetase mutations in a patient with peripheral neuropathy. Am. J. Hum. Genet., 87, 560–566.

21. Kodera,H., Osaka,H., Iai,M., Aida,N., Yamashita,A., Tsurusaki,Y., Nakashima,M., Miyake,N., Saitsu,H. and Matsumoto,N. (2015) Mutations in the glutaminyl-tRNA synthetase gene cause early-onset epileptic encephalopathy. J. Hum. Genet., 60, 97–101.

22. Zhang,X., Ling,J., Barcia,G., Jing,L., Wu,J., Barry,B.J., Mochida,G.H., Hill,R.S., Weimer,J.M., Stein,Q., et al. (2014) Mutations in QARS, encoding glutaminyl-tRNA synthetase, cause progressive microcephaly, cerebral-cerebellar atrophy, and intractable seizures. Am. J. Hum. Genet., 94, 547–558.

23. Vester,A., Velez-Ruiz,G., McLaughlin,H.M., Program,N.C.S., Lupski,J.R., Talbot,K., Vance,J.M., Zuchner,S., Roda,R.H., Fischbeck,K.H., et al. (2013) A loss-of-function variant in the human histidyl-tRNA synthetase (HARS) gene is neurotoxic in vivo. Hum. Mutat., 34, 191–199.

24. Motley,W.W., Seburn,K.L., Nawaz,M.H., Miers,K.E., Cheng,J., Antonellis,A., Green,E.D., Talbot,K., Yang,X.-L., Fischbeck,K.H., et al. (2011) Charcot-Marie-Tooth-linked mutant GARS is toxic to peripheral neurons independent of wild-type GARS levels. PLoS Genet., 7, e1002399.

25. Herzog,W., Muller,K., Huisken,J. and Stainier,D.Y.R. (2009) Genetic evidence for a noncanonical function of seryl-tRNA synthetase in vascular development. Circ. Res., 104, 1260–1266.

26. Han,J.M., Jeong,S.J., Park,M.C., Kim,G., Kwon,N.H., Kim,H.K., Ha,S.H., Ryu,S.H. and Kim,S. (2012) Leucyl-tRNA synthetase is an intracellular leucine sensor for the mTORC1-signaling pathway. Cell, 149, 410–424.

27. Belli,G., Gari,E., Piedrafita,L., Aldea,M. and Herrero,E. (1998) An activator/repressor dual system allows tight tetracycline-regulated gene expression in budding yeast. Nucleic Acids Res., 26, 942–947.

28. Gietz,R.D. and Woods,R.A. (2002) Transformation of yeast by lithium acetate/single-stranded carrier DNA/polyethylene glycol method. Methods Enzymol., 350, 87–96.

29. Laughery,M.F., Hunter,T., Brown,A., Hoopes,J., Ostbye,T., Shumaker,T. and Wyrick,J.J. (2015) New vectors for simple and streamlined CRISPR–Cas9 genome editing in *Saccharomyces cerevisiae*. Yeast, 32, 711–720.

30. Ban,N., Beckmann,R., Cate,J.H.D., Dinman,J.D., Dragon,F., Ellis,S.R., Lafontaine,D.L.J., Lindahl, L., Liljas,A., Lipton,J.M., et al. (2014) A new system for naming ribosomal proteins. Curr. Opin. Struct. Biol., 24, 165–169.

31. Mueller,P.P. and Hinnebusch,A.G. (1986) Multiple upstream AUG codons mediate translational control of GCN4. Cell, 45, 201–207.

32. Finkelstein,D.B. and Strausberg,S. (1983) Heat shock-regulated production of *Escherichia coli* beta-galactosidase in Saccharomyces cerevisiae. Mol. Cell. Biol., 3, 1625–1633.

33. von der Haar,T. (2007) Optimized protein extraction for quantitative proteomics of yeasts. PLoS One, 2, e1078.

34. Harlow,E. and Lane,D. (1988) Antibodies: a laboratory manual Cold Spring Harbor Laboratory Press, Cold Spring Harbor, N.Y.

35. Hill,D.E. and Struhl,K. (1986) A rapid method for determining tRNA charging levels in vivo: analysis of yeast mutants defective in the general control of amino acid biosynthesis. Nucleic Acids Res., 14, 10045–10051.

36. Varshney,U., Lee,C.P. and RajBhandary,U.L. (1991) Direct analysis of aminoacylation levels of tRNAs in vivo. Application to studying recognition of *Escherichia coli* initiator tRNA mutants by glutaminyl-tRNA synthetase. J. Biol. Chem., 266, 24712–24718.

37. Edgar,R., Domrachev,M. and Lash,A.E. (2002) Gene Expression Omnibus: NCBI gene expression and hybridization array data repository. Nucleic Acids Res., 30, 207–210.

38. Kim,D., Pertea,G., Trapnell,C., Pimentel,H., Kelley,R. and Salzberg,S.L. (2013) TopHat2: accurate alignment of transcriptomes in the presence of insertions, deletions and gene fusions. Genome Biol., 14, R36-2013-14-4-r36.

39. Trapnell,C., Roberts,A., Goff,L., Pertea,G., Kim,D., Kelley,D.R., Pimentel,H., Salzberg,S.L., Rinn,J.L. and Pachter,L. (2012) Differential gene and transcript expression analysis of RNA-seq experiments with TopHat and Cufflinks. Nat. Protoc., 7, 562–578.

40. de Godoy,L.M., Olsen,J. V, de Souza,G.A., Li,G., Mortensen,P. and Mann,M. (2006) Status of complete proteome analysis by mass spectrometry: SILAC labeled yeast as a model system. Genome Biol., 7, R50-2006-7-6-r50.

41. Gruhler,S. and Kratchmarova,I. (2008) Stable isotope labeling by amino acids in cell culture (SILAC). Methods Mol. Biol., 424, 101–111.

42. Stansfield,I., Grant,G.M., Akhmaloka and Tuite,M.F. (1992) Ribosomal association of the yeast SAL4 (SUP45) gene product: implications for its role in translation fidelity and termination. Mol. Microbiol., 6, 3469–3478.

43. Bonnin,P., Kern,N., Young,N.T., Stansfield,I. and Romano,M.C. (2017) Novel mRNA-specific effects of ribosome drop-off on translation rate and polysome profile. PLOS Comput. Biol., 13, e1005555.

44. Ingolia,N.T., Ghaemmaghami,S., Newman,J.R. and Weissman,J.S. (2009) Genome-wide analysis in vivo of translation with nucleotide resolution using ribosome profiling. Sci. (New York, NY), 324, 218–223.

45. Cook,L.J., Zia,R.K. and Schmittmann,B. (2009) Competition between multiple totally asymmetric simple exclusion processes for a finite pool of resources. Phys. Rev. Stat. nonlinear, soft matter Phys., 80, 31142.

46. Ciandrini,L., Stansfield,I. and Romano,M.C. (2013) Ribosome traffic on mRNAs maps to gene ontology: genome-wide quantification of translation initiation rates and polysome size regulation. PLoS Comput. Biol., 9, e1002866.

47. Miura,F., Kawaguchi,N., Yoshida,M., Uematsu,C., Kito,K., Sakaki,Y. and Ito,T. (2008) Absolute quantification of the budding yeast transcriptome by means of competitive PCR between genomic and complementary DNAs. BMC Genomics, 9, 574.

48. Brackley,C.A., Romano,M.C. and Thiel,M. (2011) The dynamics of supply and demand in mRNA translation. PLoS Comput. Biol., 7, e1002203.

49. Arava,Y., Wang,Y., Storey,J.D., Liu,C.L., Brown,P.O. and Herschlag,D. (2003) Genome-wide analysis of mRNA translation profiles in Saccharomyces cerevisiae. Proc. Natl. Acad. Sci. U. S. A., 100, 3889–3894.

50. Shah,A.A., Giddings,M.C., Parvaz,J.B., Gesteland,R.F., Atkins,J.F. and Ivanov,I.P. (2002) Computational identification of putative programmed translational frameshift sites. Bioinformatics, 18, 1046–1053.

51. Sorensen,M.A. and Pedersen,S. (1991) Absolute in vivo translation rates of individual codons in *Escherichia coli*. The two glutamic acid codons GAA and GAG are translated with a threefold difference in rate. J. Mol. Biol., 222, 265–280.

52. Dittmar,K.A., Sorensen,M.A., Elf,J., Ehrenberg,M. and Pan,T. (2005) Selective charging of tRNA isoacceptors induced by amino-acid starvation. EMBO Rep., 6, 151–157.

53. Sorensen,M.A. (2001) Charging levels of four tRNA species in Escherichia coli Rel(+) and Rel(-) strains during amino acid starvation: a simple model for the effect of ppGpp on translational accuracy. J. Mol. Biol., 307, 785–798.

54. Jakubowski,H. and Goldman,E. (1984) Quantities of individual aminoacyl-tRNA families and their turnover in Escherichia coli. J. Bacteriol., 158, 769 LP–776.

55. Kemp,A.J., Betney,R., Ciandrini,L., Schwenger,A.C.M., Romano,M.C. and Stansfield,I. (2013) A yeast tRNA mutant that causes pseudohyphal growth exhibits reduced rates of CAG codon translation. Mol. Microbiol., 87.

56. Gilchrist,M.A. and Wagner,A. (2006) A model of protein translation including codon bias, nonsense errors, and ribosome recycling. J. Theor. Biol., 239, 417–434.

57. Chu,D., Barnes,D.J. and von der Haar,T. (2011) The role of tRNA and ribosome competition in coupling the expression of different mRNAs in *Saccharomyces cerevisiae*. Nucleic Acids Res., 39, 6705–6714.

58. von der Haar,T. (2008) A quantitative estimation of the global translational activity in logarithmically growing yeast cells. BMC Syst. Biol., 2, 87.

59. Kulak,N.A., Pichler,G., Paron,I., Nagaraj,N. and Mann,M. (2014) Minimal, encapsulated proteomic-sample processing applied to copy-number estimation in eukaryotic cells. Nat. Methods, 11, 319.

60. Arita,T., Morimoto,M., Yamamoto,Y., Miyashita,H., Kitazawa,S., Hirayama,T., Sakamoto,S., Miyamoto,K., Adachi,R., Iwatani,M., et al. (2017) Prolyl-tRNA synthetase inhibition promotes cell death in SK-MEL-2 cells through GCN2-ATF4 pathway activation. Biochem. Biophys. Res. Commun., 488, 648–654.

61. Lanker,S., Bushman,J.L., Hinnebusch,A.G., Trachsel,H. and Mueller,P.P. (1992) Autoregulation of the yeast lysyl-tRNA synthetase gene GCD5/KRS1 by translational and transcriptional control mechanisms. Cell, 70, 647–657.

62. Natarajan,K., Meyer,M.R., Jackson,B.M., Slade,D., Roberts,C., Hinnebusch,A.G. and Marton,M.J. (2001) Transcriptional profiling shows that Gcn4p is a master regulator of gene expression during amino acid starvation in yeast. Mol. Cell. Biol., 21, 4347–4368.

63. Berriz,G.F., Beaver,J.E., Cenik,C., Tasan,M. and Roth,F.P. (2009) Next generation software for functional trend analysis. Bioinformatics, 25, 3043–3044.

64. Schaechter,M., MaalØe,O. and Kjeldgaard,N.O. (1958) Dependency on Medium and Temperature of Cell Size and Chemical Composition during Balanced Growth of Salmonella typhimurium. Microbiology, 19, 592–606.

65. Scott,M., Gunderson,C.W., Mateescu,E.M., Zhang,Z. and Hwa,T. (2010) Interdependence of Cell Growth and Gene Expression: Origins and Consequences. Science (80-.)., 330, 1099–1102.

66. Waldron,C. and Lacroute,F. (1975) Effect of growth rate on the amounts of ribosomal and transfer ribonucleic acids in yeast. J. Bacteriol., 122, 855–865.

67. Ghaemmaghami,S., Huh,W.K., Bower,K., Howson,R.W., Belle,A., Dephoure,N., O’Shea,E.K. and Weissman,J.S. (2003) Global analysis of protein expression in yeast. Nature, 425, 737–741.

68. Chan,P.P. and Lowe,T.M. (2009) GtRNAdb: a database of transfer RNA genes detected in genomic sequence. Nucleic Acids Res., 37, D93–7.

69. Hinnebusch,A.G. (1997) Translational regulation of yeast *GCN4*. A window on factors that control initiator-trna binding to the ribosome. J. Biol. Chem., 272, 21661–21664.

70. Steenweg,M.E., Ghezzi,D., Haack,T., Abbink,T.E., Martinelli,D., van Berkel,C.G., Bley,A., Diogo,L., Grillo,E., Te Water Naude,J., et al. (2012) Leukoencephalopathy with thalamus and brainstem involvement and high lactate ‘LTBL’ caused by EARS2 mutations. Brain, 135, 1387–1394.

71. Stephen,J., Nampoothiri,S., Banerjee,A., Tolman,N.J., Penninger,J.M., Elling,U., Agu,C.A., Burke,J.D., Devadathan,K., Kannan,R., et al. (2018) Loss of function mutations in VARS encoding cytoplasmic valyl-tRNA synthetase cause microcephaly, seizures, and progressive cerebral atrophy. Hum. Genet., 137, 293–303.

72. Ishimura,R., Nagy,G., Dotu,I., Chuang,J.H. and Ackerman,S.L. (2016) Activation of GCN2 kinase by ribosome stalling links translation elongation with translation initiation. Elife, 5, e14295.

73. Harding,H.P., Zhang,Y., Zeng,H., Novoa,I., Lu,P.D., Calfon,M., Sadri,N., Yun,C., Popko,B., Paules,R., et al. (2003) An Integrated Stress Response Regulates Amino Acid Metabolism and Resistance to Oxidative Stress. Mol. Cell, 11, 619–633.

74. Li,W., Wang,X., van der Knaap,M.S. and Proud,C.G. (2004) Mutations Linked to Leukoencephalopathy with Vanishing White Matter Impair the Function of the Eukaryotic Initiation Factor 2B Complex in Diverse Ways. Mol. Cell. Biol., 24, 3295–3306.

75. Richardson,J.P., Mohammad,S.S. and Pavitt,G.D. (2004) Mutations Causing Childhood Ataxia with Central Nervous System Hypomyelination Reduce Eukaryotic Initiation Factor 2B Complex Formation and Activity. Mol. Cell. Biol., 24, 2352–2363.

76. Chelliah, V. et al. BioModels: ten-year anniversary. Nucl. Acids Res. 2015, 43(Database issue):D542-8 Chelliah,V., Juty,N., Ajmera,I., Ali,R., Dumousseau,M., Glont,M., Hucka,M., Jalowicki,G., Keating,S., Knight-Schrijver,V., Lloret-Villas,A., Natarajan,K.N., Pettit,J.B., Rodriguez,N., Schubert,M., Wimalaratne,S.M., Zhao,Y., Hermjakob,H., Le Novère,N. and Laibe,C. (2015) BioModels: ten-year anniversary. Nucleic Acids Res., 43(Database issue):D542–8.

77. Vizcaíno,J.A., Deutsch,E.W., Wang,R., Csordas,A., Reisinger,F., Ríos,D., Dianes,J.A., Sun,Z., Farrah,T., Bandeira,N., Binz,P.A., Xenarios,I., Eisenacher,M., Mayer,G., Gatto,L., Campos,A., Chalkley,R.J., Kraus,H.J., Albar,J.P., Martinez-Bartolomé,S., Apweiler,R., Omenn,G.S., Martens,L., Jones,A.R. and Hermjakob,H. (2014). ProteomeXchange provides globally coordinated proteomics data submission and dissemination. Nature Biotechnol. 30, 223–226.

