## Supplementary information for "The molecular aetiology of tRNA synthetase depletion: induction of a *GCN4* amino acid starvation response despite homeostatic maintenance of charged tRNA levels"

**ABSTRACT**

During protein synthesis, charged tRNAs deliver amino acids to translating ribosomes, and are then re-charged by tRNA synthetases (aaRS). In humans, mutant aaRS cause a diversity of neurological disorders, but their molecular aetiologies are incompletely characterised. To understand system responses to aaRS depletion, the yeast glutamine aaRS gene (*GLN4*) was transcriptionally regulated using doxycycline by *tet-off* control. Depletion of Gln4p inhibited growth, and induced a *GCN4* amino acid starvation response, indicative of uncharged tRNA accumulation and Gcn2 kinase activation. Using a global model of translation that included aaRS recharging, Gln4p depletion was simulated, confirming slowed translation. Modelling also revealed that Gln4p depletion causes negative feedback that matches translational demand for Gln-tRNA$^{Gln}$ to aaRS recharging capacity. This maintains normal charged tRNA$^{Gln}$ levels despite Gln4p depletion, confirmed experimentally using tRNA Northern blotting. Model analysis resolves the paradox that Gln4p depletion triggers a *GCN4* response, despite maintenance of tRNA$^{Gln}$ charging levels, revealing that normally, the aaRS population can sequester free, uncharged tRNAs during aminoacylation. Gln4p depletion reduces this sequestration capacity, allowing uncharged tRNA$^{Gln}$ to interact with Gcn2 kinase. The study sheds new light on mutant aaRS disease aetiologies, and explains how aaRS sequestration of uncharged tRNAs can prevent *GCN4* activation under non-starvation conditions.

### Supplementary Materials

#### Stochastic simulations

##### *Global translation model:*

We use the Gillespie algorithm (1) to generate reactions taking place at random times according to their propensities. The Global Translation Model (GTM) is based on the Totally Asymmetric Simple Exclusion Process (TASEP) and has four main types of reactions: A ribosomal particle may i) enter the lattice, ii) translocate on the lattice, which uses up one tRNA, iii) leave the lattice by dropping off at any site of the lattice or terminating at the last lattice site, or iv) an uncharged tRNA may be recharged. Note that the ribosomal particles have a footprint of size $W$ codons. In the following, 'ribosome' means the translational site of the ribosome, i.e., the translocation from the initiation lattice site $i = 0$ to $i = 1$ is only possible if the lattice site $i = W + 1$ is empty.

We start at time $t = 0$ with $M$ empty mRNAs, each of which has $N_m$ lattice sites. Moreover, all tRNAs are charged, $T_i^t = T_i^c$. After a dwell time $\tau$, which is drawn from an exponential distribution determined by the propensities at that time point, the next reaction is randomly chosen.

Then, all propensities, numbers of species and state of the lattices are updated, and the procedure is repeated until the predefined end time $t_{max}$. We allow a certain transient time before we store the information of density profiles and the properties, in order to make sure that we reach the steady state, i.e., the total integration time considered to compute average quantities is $t = t_{max} - t_{trans}$.

##### *Specification of parameters; Global Translation Model*

A real *S. cerevisiae* cell has $10^7$ codons (derived using a knowledge of mRNA numbers and types per cell (2)), $3 \times 10^6$ tRNAs (3) and $2 \times 10^5$ ribosomes (4).

We chose a downscaling factor of 20 for our reduced system, corresponding to $5 \times 10^5$ codons and $10^5$ tRNAs. We include $1.5 \times 10^3$ individual mRNAs with individual length according to the gene ontology classification, see Table S11. All other parameters are downscaled accordingly: the downscaled system contains $10^4$ ribosomes, 1.5 x $10^5$ tRNAs (downscaled from total tRNAs/cell estimated at $3 \times 10^6$ (3)). The abundance of each tRNA species is assumed to be proportional to the tRNA gene copy number, with a factor of proportionality of $T_t$ = 526 for the 20-fold downscaled cell (see Table S3; (57)).

Note that the number of ribosomes in the reservoir changes in time. The fraction of ribosomes in the cell which are engaged in translation has been calculated for our setup to 75%, which is close to the value reported in (6). Then, the number of free ribosomes in the cytoplasm is given by 25%, i.e., in our downscaled cell, we have on average 2500 ribosomes available for initiation in steady state. Note that the simulation starts with empty mRNAs, i.e., initially, all ribosomes are free.

The initial initiation rates are listed in Table S2 for the downscaled cell, with the average initiation rate amounting to $\langle\alpha\rangle = 0.15/s$.

The maximum charging rate for the tRNA-synthetases $V_{max,i}$ is estimated by dividing the total number of each amino acids incorporated into proteins during the cell lifetime by the abundance of the corresponding tRNA-synthetases (7)(8). Interestingly, the $V_{max,i}$ values highly correlate with the tRNA gene copy number. The values of the Michaelis-Menten constants $K_{M,i}$ are not known for all synthetases, but the fact that the charging level of tRNAs under physiological conditions has been estimated to be 80% across different tRNA species (9-13) implies that the ratio $\xi = V_{max,i}/K_{M,i}$ is similar for all different synthetases (see Table S4). We use the 80% charging level to estimate this ratio in the simulations (we use the charging capacity of $\xi = V_{max,i}/K_{M,i} = 5/s$). Note that this ratio is proportional to the aminoacylation efficiency $k_{cat,i}/K_{M,i}$.

Considering wobble base pairing via the wobble factor $w(j)$ for each codon $j$, we reduce the hopping rates of the codons which use the G-U wobble by 39% compared to their G-C wobble counterparts, and those of codons using the I-G wobble by 36% relative to their I-U counterparts, based on the results in (14). In addition to these choices, a 60% reduction has been introduced for the so-called 'missing tRNAs' which are non-perfect matches and which do not have a supplement (15). All wobble examples are listed together with the initial hopping rates in Table S3 for the downscaled cell.

The hopping rate $k(j)$ for each codon $j$ is proportional to the number $T_j^c$ of charged tRNAs for this type of codon, with the proportionality factor $r$. Considering the wobble base pairing, the hopping rate becomes $k(j) = rT_j^c w(j)$. In order to determine the proportionality factor $r$, we use the fact that the average hopping rate has been estimated to be $\langle k(j)\rangle = 10/s$ (6)(16)(17). Starting with the assumption of a homogeneous charging level of 80% in the cell, the number of charged tRNAs is estimated as $T_j^c = 0.8\, T_j^t = 0.8\, Tt\, GCN(j)$. Then, the average hopping rate over all codons is given by $\langle k(j)\rangle = r0.8Tt \sum w\,(j)GCN\,(j)/N$ where the sum is over all codons $j = 1, \dots, N$. For a given system size, we can now retrieve the factor of proportionality $r$ , and for the considered downscaled system we obtain $r = 3.696 \times 10^{-3}$. Note that although the average value does not depend on the system size and does not change throughout the simulation, the hopping rate in every single Gillespie step depends on the number of charged tRNAs of the corresponding type.

Furthermore, we use the drop-off rate $\gamma = 5.6 \times 10^{-3}$/s as estimated (18) and the termination rate $\beta = 38.81$/s (19).

The average codon length of mRNA for each GO-Slim class was calculated. The ratio between total number of codons within this category and the number of mRNAs defines the average length of the single representative mRNA of this category. Then we determined the average number of each codon on the representative mRNA by dividing the number of codons of that type by the number of mRNAs within this category. Note that doing so, we neglected codons due to rounding effects, but the loss is less than 0.5%.

The code for the Global Translation Model is available through https://www.ebi.ac.uk/biomodels/ (23) with the following submission identifier following submission identifier: MODEL2001080004. The code is written in C language and adapts some of the structures developed in (22) to access the codons of the mRNA lattices.

##### *Synthetase Sequestration Model*

The synthetase sequestration model includes four main transitions for each amino acid type *i*: the binding rate $b_i$, the charging rate $\tilde{X}_i$, the usage rate $\tilde{k}_i$, and the synthetase-tRNA-complex dissociation rate $b_0$. We start at time $t = 0$ with 20 amino acids, with a starting configuration where all tRNAs are *empty*, i.e., the number of empty tRNAs is $T_i^e = T_i^t$ and the number of bound and charged tRNAs is zero. To ensure steady state results, we allow the transient time $t_{trans} = 5 \times 10^8\ steps$ to pass before we start to record the number of *charged*, *bound* and *empty* tRNAs of each amino acid type. The average charging level $\langle c_i \rangle$ is equal to the number of charged tRNAs of type *i*, weighted by the time the individual tRNAs of type *i* stay in the charged state, divided by the total number of tRNAs of this type and the total run time. Likewise, we obtain the bound level $\langle b_i \rangle$ and the empty level $\langle e_i \rangle$.

The synthetase sequestration simulation is carried out for the whole cell, i.e., we use the values from Table S4 for the number $E_{0,i}$ of synthetase molecules and the catalytic rate $k_{cat,i}$. The number of synthetase molecules reflects the total amount of synthetase in the cell, i.e, the free synthetase molecules and the ones captured within the synthetase-tRNA-complexes. However, the binding rates depend on the number of free synthetase molecules, i.e, $E_i = \tilde{E}_{0,i} - T_i^b$. Note that it is the total amount which is affected by doxycycline for the glutamine case: $\tilde{E}_{0,Gln} = dE_{0,Gln}$ with the Gln4 protein ratio $d$.

The code for the Synthetase Sequestration Model is available through the BioModels database https://www.ebi.ac.uk/biomodels/ (23) with the following submission identifier: MODEL2001080005

***Specification of parameters; Synthetase Sequestration Model***

The usage rate constant $k_3 = 0.1/s$ is estimated to represent the average translation rate. The charging rate constants $k_{2,i} = ak_{cat,i}$ are fitted to comply an average charging level of 80%, which leads to the factor of proportionality of $a = 0.2$. The $k_{cat,i}$ and the enzymatic concentrations of the synthetases $E_{0,i}$ are chosen analogously to the global translation model (Table S4). Note that as a matter of consistency, we adjust the number of glutamine synthetase molecules to the construct, $E_{0,Gln} \approx 10^5$.

The rate constants $k_0 = 43/s$ and $k_{1,Gln} = 0.007/s$ are taken from Uter (20). Note that we have converted the units of the binding rate constant. Using the quasi equilibrium approximation $K_{M,i} = k_0/k_{1,i}$, the binding rate constants of the other amino acids are derived by the relation $V_{max,i}/K_{M,i} = E_{0,i}k_{cat,i}/(k_0/k_{1,i})$. Corresponding to the global translation model, the Michaelis-Menten constant $K_{M,i} \propto V_{max,i}$ is proportional to the charging velocity $V_{max,i} = E_{0,i}k_{cat,i}$ with the factor of proportionality $\xi = 15$, which is thought to be the same for all tRNAs, and estimated by using $K_{M,Gln} = 5414$ molecules per cell, corresponding to the value given in Uter (20). Note that in the stochastic simulation we reach a steady state in the bound level, since we never run out of substrate, as the tRNAs are replenished throughout the cycle. Additionally, the dissociation rate constant $k_0 \gg k_{2,i}$ is much larger than the charging rate constants, and therefore, the conditions are fulfilled for the approximation to be appropriate.

***General expression for the usage rate***

For each tRNA the $usagerate = \sum M_u \, n_{u,i} J_u$ is proportional to the ribosomal current $J_u$ along the mRNA of type $u$, the number $M_u$ of copies of mRNA of type $u$, the number $n_{u,i}$ of codons on the mRNA of type $u$ decoded by tRNA of type $i$.

Every time a ribosome hops with rate $k_j$ from one codon to the next, a charged tRNA is used. Together with the ribosomal density $\rho_j$, the usage rate can be expressed by

$usagerate = \sum M_u \sum k_{p(j),u} \, \rho_{p(j),u}\left(1 - \rho_{p(j)+1,u}\right)$ with the position $p_{(j),u}$ of the $j^{th}$ codon which is decoded by tRNA $i$ on mRNA of type $u$.

***Autogenous Feedback***

For the Global Translation Model (GTM) autogenous feedback we analysed different effects that reduced growth rate has on the initiation rate. In the GTM, the initiation rate is given by $\alpha = \alpha_0 R_{free}$, where $R_{free}$ denotes the number of free ribosomes in the cytoplasm (not engaged in translation), and

$\alpha_0$ is a proportionality constant that comprises other effects on the initiation, such as the availability of initiation factors or secondary structures on the 5' UTR. In principle, in response to stress we expect that both $\alpha_0$ and $R_{free}$ are reduced ($\alpha_0$ decreases because the number of initiation factors can be assumed to decrease with the growth rate, and $R_{free}$ decreases because ribosomes are coupled to the growth rate). Given the experimental data on how the number of ribosomes decreases with Gln4p (Fig. 6), we tested what are the effects of the reduction of $\alpha_0$ and $R_{free}$ on the simulation results.

Using the data in Figure 6 for calibration of the ribosomal reduction, we obtain the simulation results shown in Fig. S5: when comparing these results to the ones obtained with no feedback (blue solid line), we see that the effect of ribosome reduction is very minor. However, the reduction in the initiation factors availability and therefore $\alpha_0$ (assumed to decrease proportionally to the growth rate) has a marked effect (shown as a solid red line in Fig. S5), especially on the charging level of the Glutamine codons. Therefore, for the sake of simplicity, we have now only considered the effect of the reduction in $\alpha_0$.

Moreover, we analysed the effects of tRNA reduction with decreasing growth rate, since in principle not only ribosomes but also tRNAs are downregulated under stress, e.g., through the TOR pathway (21). In order to assess the effect of downregulation of tRNA on our results, we have tested the effects of a tRNA feedback in the GTM, by reducing the amount of tRNA proportionally to the experimentally obtained reduction in ribosomal content. The results are shown in Fig. S5. The green dashed line shows the results of the model incorporating the tRNA feedback. By comparing these results with the ones obtained with the GTM with no feedback (blue solid line), we see that the reduction of the number of tRNAs under stress has only a very minor effect. Therefore, we have neglected it for the sake of simplicity and do not consider tRNA reduction in the rest of the model simulations.

***Correlation analysis of the current and the glutamine content***

To address the question whether the mRNAs with a high glutamine codon density are more sensitive to increased doxycycline concentration, we did a correlation analysis of the ratio $J_a(+doxy)/J_a(-doxy)$ of the current with and without doxycycline of the GO-Slim category a with the glutamine codon content per mRNA within this GO-Slim category.

In the GTM with no feedback, there is a strong correlation between $J_a(+doxy)/J_a(-doxy)$ and the CAG codon content (Spearman's correlation coefficient of -0.88). Depletion of charged Gln tRNAs causes elongation arrests, and therefore, a reduction in the current.

With autogenous feedback, on the other hand, there is a strong reduction in the correlation with the glutamine content (Spearman's correlation coefficient of 0.22), which is not surprising, because here, the balance between supply and demand is restored; translation initiation rate is decreased so that the charged Gln tRNA available can keep up with the demand.

Note that there is a weaker correlation between reduction in current and the non-rare CAA codon content per mRNA (Spearman's correlation coefficients -0.76 without feedback and 0.18 with feedback).

Figure S3 (panel A) shows the correlation between the CAG content of a mRNA within GO-Slim category a and the ratio $J_a(+doxy)/J_a(-doxy)$ with (red line) and without (blue line) feedback. In panel B, the ratio $J_a(+doxy)/J_a(-doxy)$ with (red line) and without (blue line) feedback together with the CAG content (green line) of a mRNA within GO Slim category a is shown for each GO-Slim category a, ranked by the doxycycline influence on the ratio $J_a(+doxy)/J_a(-doxy)$ without feedback.

***Synthetase sequestration at different charging levels***

As a matter of consistency with the literature, we used the average 80% tRNA charging level as data to parameterise our model. If we instead use 60% tRNA charging level, as suggested by the experimental results in Fig. 7, the results remain qualitatively the same as shown in Fig. S1A.

The glutamine charging level stays more or less constant whereas the empty level increases with increasing doxycycline concentration (decreasing Gln4 protein ratio) at the expense of the bound level.

Note that a smaller global charging level of the cell leads to a smaller global current. Therefore, the response on doxycycline evolves slightly differently. We use the GTM to inform the SSM how the threshold for the doxycycline sensitive response of the usage rate is shifted; for 80% charging the feedback for the SSM is apparent at Gln4 protein ratio 0.2 which corresponds to the threshold current $J_{thresh} = 0.03/s$ (see Fig. S1B). For 60% charging, however, this value of the current is already reached at the Gln4 protein ratio of 0.35, as shown by the red arrows in Fig. S1B.

***Density profiles***

Figure S1 shows the profile of the ribosomal density along an mRNA from GO Slim category 66, 'regulation of transport' (bottom), which has a large abundance of slow codons (19).

Comparison to the density profile shown in the main part of the manuscript (Fig. 5H) reveals a similar behaviour, although the two categories are quite different: category 16 has 299 mRNAs in the downscaled cell with an average length of 158 codons and on average 0.03 CAG codons on each mRNA. In contrast, category 66 has only two mRNAs in the downscaled cell, but with an average length of 920 codons per mRNA and 8.6 CAG codons on each mRNA, see Table S10, below.

| category | 16 | 66 |
|---|---|---|
| CAA | 33403 | 1514 |
| CAG | 258 | 412 |
| glutamines | 33661 | 1926 |
| all codons | 942334 | 43881 |
| mRNAs | 5946 | 48 |
| CAA/all codons [%] | 3.5 | 3.5 |
| CAG/all codons [%] | 0.03 | 0.9 |
| CAA/ glutamines [%] | 99.2 | 78.6 |
| CAG/ glutamines [%] | 0.8 | 21.4 |
| CAA/ mRNAs | 5.62 | 31.5 |
| CAG/ mRNAs | 0.04 | 8.6 |
| glutamines/mRNAs | 5.66 | 40.1 |

**Table S10:** A list of number of, CAG, CAA and CAG+CAA (glutamine) codons, number of all codons and mRNAs from the GO-Slim categories 16 and 66. The proportion of the CAG, CAA and CAG+CAA codons compared to all codons in the corresponding category, proportion of the CAG and CAA codons with respect to all glutamine codons within the category and the average number of CAG, CAA and CAG+CAA codons on an mRNA within the category.

The position of the glutamine codons on the mRNA in (Fig. S4) is highlighted by the grey dashed bars. The black line represents the situation without doxycycline. Both, the blue and the red lines represent the situation in the presence of doxycycline, corresponding to a Gln4 protein ratio of 5%, with (red line) and without (blue line) feedback loop.

Without feedback and in the presence of doxycycline, the ribosome density profile in (Fig. S4) is highly inhomogeneous (blue line); ribosomal queues build up behind glutamine codons. When the feedback mechanism is switched on, the profile is much smoother and the queues vanish (red dashed line).

In category 16, the ribosome density along the mRNA in the case with doxycycline is large compared to the case without doxycycline (see figure 6H), whereas in category 66, with doxycycline the ribosomal density is also large at the entrance of the lattice but decreases rapidly along the mRNA, due to the bottleneck created by the accumulation of rare codons.

In contrast, when the autogenous feedback mechanism is activated, ribosomes are equally distributed in a low density-like regime all over the mRNA, no queueing is visible for any of the two categories. Note that with feedback, the average density in both categories decreases considerably in the presence of doxycycline, which is also reflected in the protein production rate, see (Fig. 6D) in the main text.

Note that the ribosomal particles in the simulation have a footprint of $W = 9$, which leads to the plateau peaks obtained in the case without feedback. Note further that here, the translational site of the ribosome is on the left side, which produces the overhang peaks.

#### Figures

***Figure S1*****:** Panel A: Resulting plot of the steady state charging levels of the glutamine tRNAs as a function of the Gln4 protein ratio (an average of 60% tRNA charging level was used to parameterise our model). Yellow line: mean level $\langle c \rangle$ of charged glutamine tRNAs; Red line: mean level $\langle b \rangle$ of bound glutamine tRNAs; Blue line: mean level $\langle e \rangle$ of empty glutamine tRNAs. The grey line indicates the feedback threshold d = 0.35: for Gln4 protein ratios smaller than 0.35 the usage rate decreases proportionally to the growth rate.

Panel B: Global current J with autogenous feedback of the GTM for two different global charging levels: solid blue line corresponds to the global current for the global charging level at 80% as in Fig.6D of the main text, the dashed blue line corresponds to the global charging level of 60%. The red arrows are a guide for the eye to show how the shift in the feedback threshold for the usage rate in the SSM is obtained.

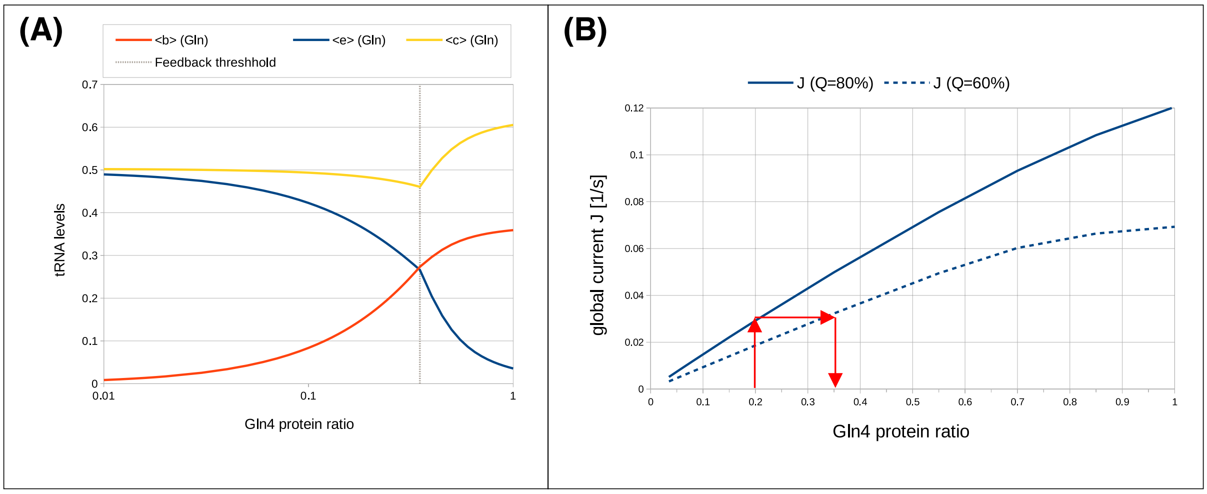


***Figure S2;*** **Depletion of the *KRS1* lysyl-tRNA synthetase using tet-off regulation causes translational induction of GCN4**

The effect of *KRS1* tRNA synthetase shut-off, using doxycycline, on uncharged tRNA accumulation and thus *GCN4* activation was measured using the *GCN4*-lacZ reporter plasmids in a tetO-*KRS1* strain. Reporter gene expression was measured in three independent biological replicates; error bars represent ± 1 standard error of the mean, n=4. Plasmid p180 measures a *GCN4* response, relative to the negative (p226) and positive (p227) controls.

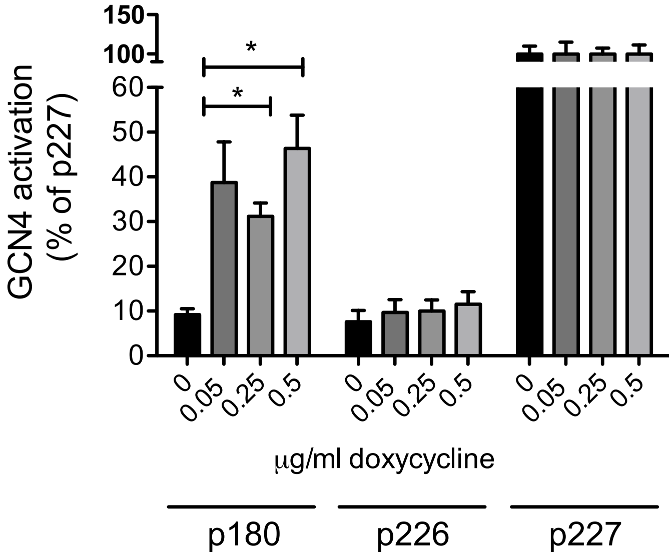

***Figure S3*:** **Phosphorylation of eIF2α in a Gln4 tRNA synthetase tet-off yeast strain in response to doxycycline.** Western blots of total cell lysates were probed with either [panel A] an anti-phospho-eIF2α antibody (phospho-Sui2p [34.7 kDa]; panel A), or a control anti-phosphoglycerate kinase (Pgk1p [44.7 kDa]; panel B) antibody to normalise for lane loading. Lanes 1-3 contain negative control samples, 3-fold overloaded to confirm the absence of phospho-eIF2α (Sui2p); wild-type yeast (lane 1); *GLN4* tet-off Δ*gcn2* (lane 2); *GLN4* tet-off Δ*gcn2*, doxycycline-treated (lane 3). Lanes 4-7 contain *GLN4* tet-off grown in the presence of increasing doxycycline concentrations. Band intensity of phospho-Sui2p was quantified using ImageJ (https://imageJ.nih.gov/ij/index.html), employing Pgk1p to normalise for lane loading variation (panel C).

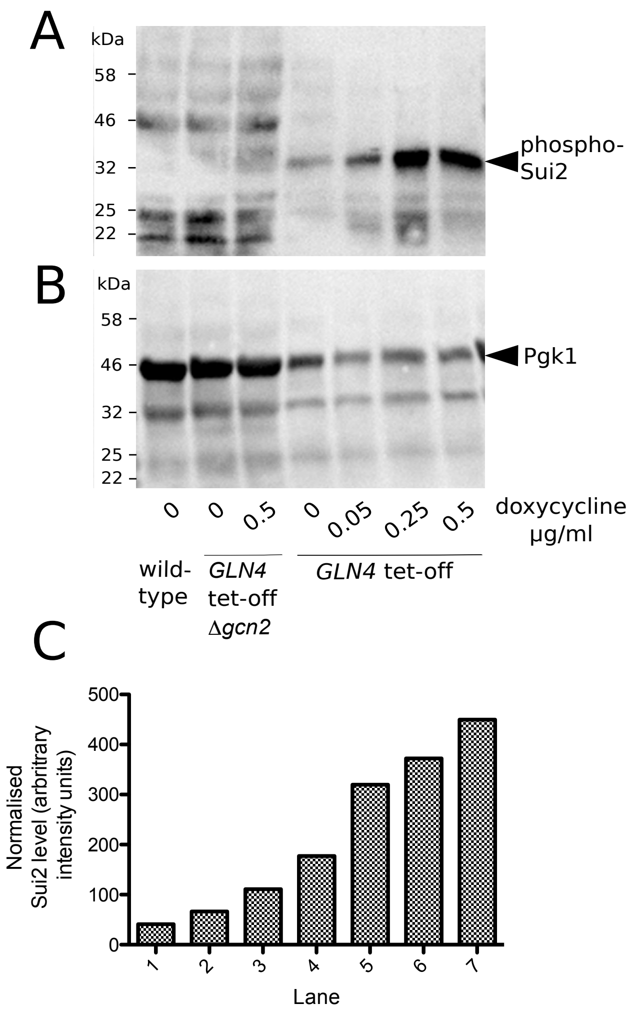

***Figure S4*: Ribosomal density profile along a representative mRNA from GO Slim category 66**

The 'regulation of transport', linear axes (top) and semi-logarithmic axes (bottom) are shown. The black line represents the situation without doxycycline and the blue and red lines represent the situation in the presence of doxycycline, without (blue line) and with (red, dashed line) feedback loop. The positions of the glutamine codons on each mRNA are indicated by the grey, dashed bars.

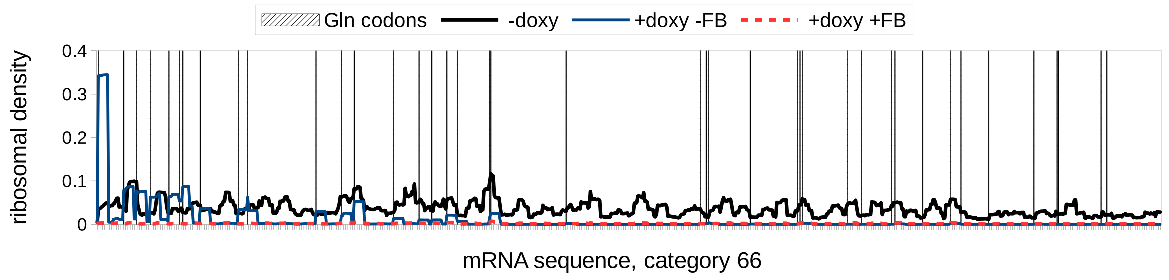


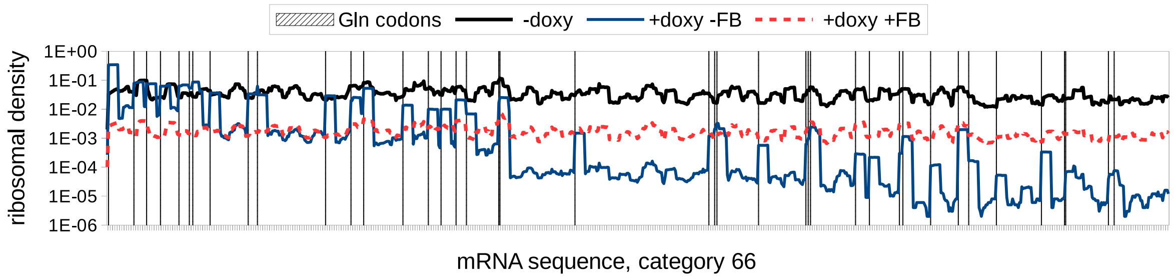

***Figure S5:*** **Different aspects of the autogenous feedback**

No feedback (blue lines) as reference, feedback on the number of ribosomes (green lines), feedback on the number of tRNAs (green dashed lines) and on the initiation factors (red lines). Upper panel: The global current (left) and the global charging level (right) without glutamine tRNAs as functions of the Gln4 protein ratio. Lower panel: The mean charging level of the glutamine tRNAs, CUG (left) and UUG (right). All results have been simulated using a cell size of 100 mRNAs.

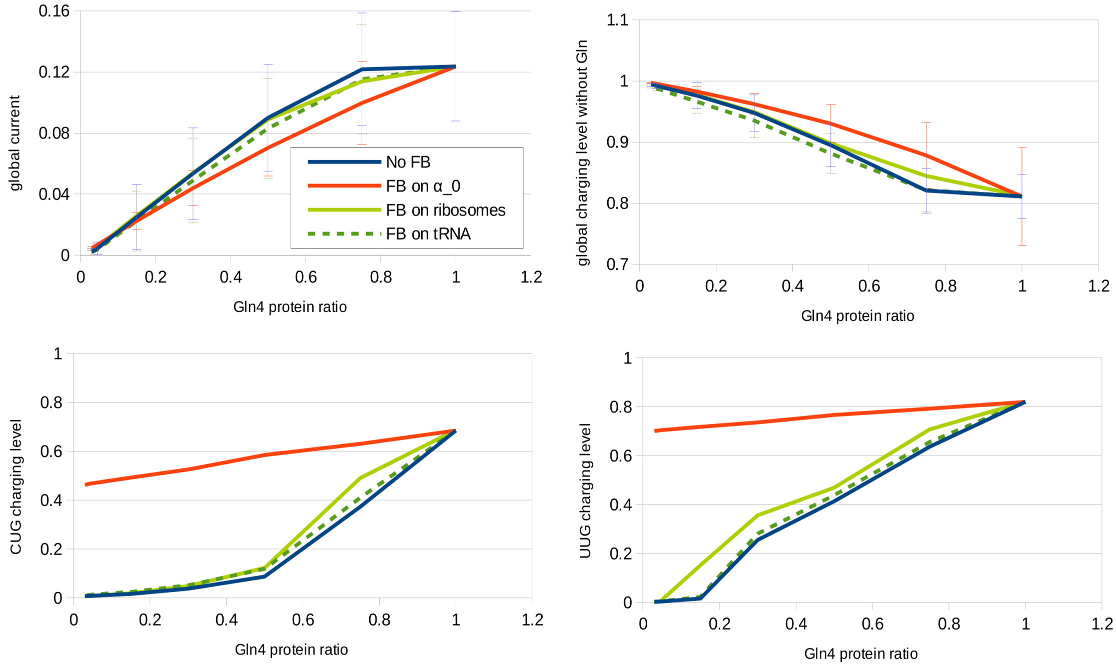


***Figure S6***: Panel A: correlation between the CAG content of a mRNA within GO Slim category $a$ and the ratio $J_a\,(+doxy)/J_a\,(-doxy)$ with (red line) and without (blue line) feedback. Panel B: the ratio $J_a\,(+doxy)/J_a\,(-doxy)$ with (red line) and without (blue line) feedback together with the CAG content (green line) of a mRNA within GO Slim category $a$ is shown for each GO-Slim category $a$, ranked by the doxycycline influence on the ratio without feedback.

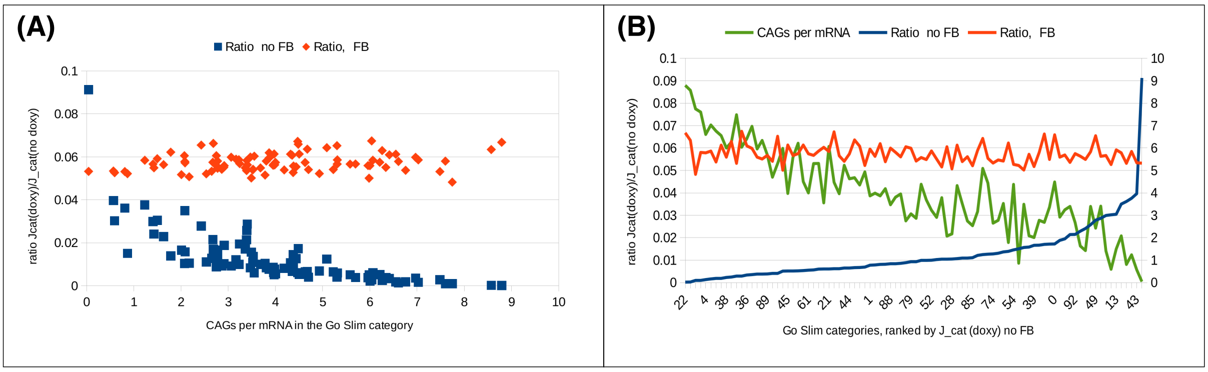

***Figure S7;* Hygromycin resistance of a wild-type strain and the Gln tRNA synthetase tet-off strain.** *Panel A-C:* the resistance to hygromycin of the *GLN4* tet-off strain was compared to its progenitor wild-type strain BGY2 by measuring mid-log phase growth rates in cells growing in growing in YPD medium containing doxycycline at either 0, 0.04 and 0.08 μg/ml. *Panel D:*, this data was processed for each concentration of doxycycline to show the growth rates as a percentage of those obtained in 0 μg/ml hygromycin. Cells were grown at 30°C, 400 rpm, in 96-well plates in a LabTech International Omega plate reader. Expressing mid-log phase growth rates in the presence, or absence of hygromycin indicated that depletion of the Gln4p tRNA synthetase did not render the cells more sensitive to hygromycin relative to the effect of hygromycin on wild-type cells (panel D).

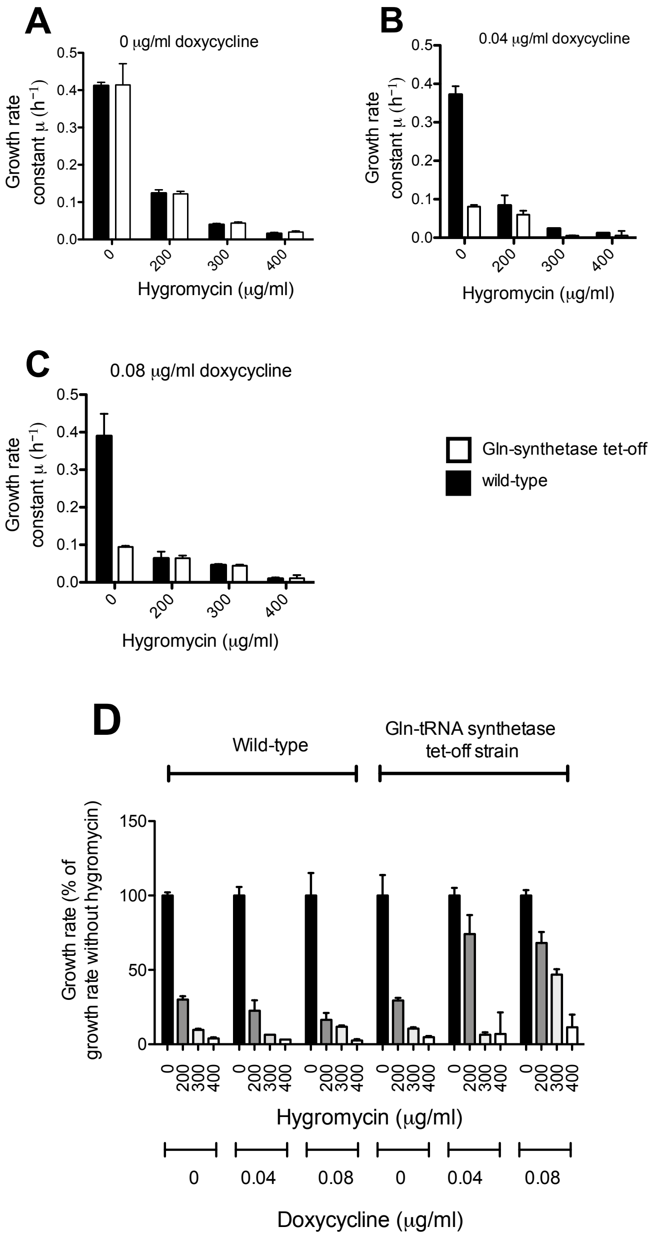

#### TABLES

***Table S1:* Oligonucleotides used in this study**

| ID | Name | Sequence 5'->3' |
|---|---|---|
| A1 | ptetO GLN4 F | AGGGATTTGATGCTTGTTTTAATGAGAAAAAAATATCAGGAGTATCAGCTGAAGCTT CGTACGC |
| A2 | ptetO GLN4 R | AGGGATTTGATGCTTGTTTTAATGAGAAAAAAATATCAGGAGTATCAGCTGAAGCTT CGTACGC |
| A3 | tet GLN4 HA R | CCCAACCTGTGAAAACAGCTGAGTCAATTCTTCTACAGAAGAAGCATAATCAGGAAC ATCGTATGGATACATATAGGCCACTAGTGGATCTG |
| A4 | pET-GLN4 3' | GTTAGCAGCCGGATCTCACTTGGAAGTTGCGTCCTTC |
| A5 | pET GLN4 F | ACGACGACAAGCATATGTATCCATACGATGTTCCTGATTATGCTTCTTCTGTAGAAGA ATTGACTC |
| A6 | $tRNA_{UUG}$ probe (Gln) | TTGTCCGGATCAAAACC |
| A7 | $tRNA_{CUU}$ probe (Lys) | CCCCTAACCTTATGATTAAGAGTC |
| A8* | Plasmid repair DNA: *GCN2* deletion | GAAGTGAAAGTTGGTGCGCATGTTTCGGCGTTCGAAACTTCTCCGCAGTGAAAGAT AAATCAAATATCAGTTAATCTTTGGTTTTAGAGCTAGAAATAGCAAGTTAAAATAAG GCTAGTCCGTTATCAACTTGAAAAAGTGGCACCGAGTCGG |
| A9 | Genome repair DNA: *GCN2* deletion | GTCTTTTAAGATTTTATAAGCATTGATTTTTTTTTTCAATAATTTTCCGTTCCCCTTAAC ACATACTATGTATAAGCCTTTAAAGATCTGTGCGGCGCTATAACGCATCAGTTAAAG GTAAAGTATATACAATGTCTTAGATACTATATT |
| A10 | gcn2D F | GGAGGAAGCAGTCACCCATC |
| A11 | gcn2D R | GCCTCACACAACATACGCAC |
| A12 | GFP genome integration forward | GCTGACTACGATGCTTTGGACATTGCTAACAGAATCGGTTACATTATGTCTAAAGGT GAAGAATTATTC |
| A13 | GFP genome integration reverse | CATGTTATGAAAGATTTATTGTATATTTAAAGAAAAATTTAAAAATAATATTAAATTT ATTAATTAAATTATTTGTACAATTCATCCATAC |
| A14 | GFP integration gRNA | GAAGTGAAAGTTGGTGCGCATGTTTCGGCGTTCGAAACTTCTCCGCAGTGAAAGAT AAATGATC TCGGTTACATTTAATCTAAT GTTTTAGAGCTAG AAATAGCAAGTTAAAATAAGGCTAGTCCGTTATCAACTTGAAAAAGTGGCACCGAG TCGG |
| A15 | *KRS1* tet-off regulation: forward primer | TTACATACATTGATTTATTGCTCTGTCCTTCCTCCGAGAAATCCGCAGCTGAAGCTTC GTACGC |
| A16 | *KRS1* tet-off regulation: reverse primer | GTTAGCAACACCTTCAGCGGCGGCTTTGACATTATCTTGTTGAGAAGCATAATCAGG AACATCGTATGGATACATATAGGCCACTAGTGGATCTG |
| A17 | $tRNA_{UCU}$ probe (Arg) | CACTCACGATGGGGGTCGAACCCATAATCTTCTGATTAGAAGTCAGACGCGTTGCCA TTA CGCCACGCGAGC |
| A18 | GFP cloning forward | GTTAGCAGCCGGATCCTTATTTGTACAATTCATCCATACCA |
| A19 | GFP cloning reverse | CATATGCTCGAGGATATGTCTAAAGGTGAAGAATTATTCA |

*Red text – Indicates *GCN2* gRNA, Green text – Indicates 5' sgRNA

*Table S2:* **Parameters used for our downscaled system**

Number of mRNAs, number of codons and the undisturbed initiation rates per GO-Slim category $a$. In total, there are $n_c = 500124$ and $n_m = 1516$ mRNAs in the downscaled cell fraction and the average initiation rate amounts to $\langle\alpha\rangle = 0.15/s$.

| a (GO-Slim) | Number of mRNAs | Number of codons | Av. Length of 1 mRNA | Initiation rate α_a [1/s] |
|---|---|---|---|---|
| 0 | 2 | 1182 | 591 | 0.0851 |
| 1 | 75 | 25650 | 342 | 0.21 |
| 2 | 23 | 10718 | 466 | 0.21 |
| 3 | 4 | 2156 | 539 | 0.191 |
| 4 | 2 | 1052 | 526 | 0.151 |
| 5 | - | - | 588 | - |
| 6 | 21 | 8736 | 416 | 0.144 |
| 7 | 51 | 24633 | 483 | 0.166 |
| 8 | 23 | 9085 | 395 | 0.107 |
| 9 | 6 | 1752 | 292 | 0.156 |
| 10 | 10 | 3360 | 336 | 0.188 |
| 11 | 17 | 6086 | 358 | 0.18 |
| 12 | 3 | 1401 | 467 | 0.128 |
| 13 | 27 | 10773 | 399 | 0.195 |
| 14 | 3 | 1533 | 511 | 0.119 |
| 15 | 2 | 1014 | 507 | 0.113 |
| 16 | 299 | 47242 | 158 | 0.207 |
| 17 | 7 | 3360 | 480 | 0.124 |
| 18 | 4 | 1316 | 329 | 0.132 |
| 19 | 7 | 2492 | 356 | 0.162 |
| 20 | 3 | 1458 | 486 | 0.11 |
| 21 | 6 | 2640 | 440 | 0.11 |
| 22 | 4 | 2948 | 737 | 0.121 |
| 23 | 6 | 2460 | 410 | 0.101 |
| 24 | 1 | 494 | 494 | 0.13 |
| 25 | 20 | 7640 | 382 | 0.2 |
| 26 | 13 | 5759 | 443 | 0.112 |
| 27 | 3 | 948 | 316 | 0.206 |
| 28 | 4 | 1492 | 373 | 0.177 |
| 29 | 26 | 11258 | 433 | 0.129 |
| 30 | 36 | 16272 | 452 | 0.126 |
| 31 | 5 | 1970 | 394 | 0.15 |
| 32 | 7 | 2534 | 362 | 0.177 |

| a (GO-Slim) | Number of mRNAs | Number of codons | Av. Length of 1 mRNA | Initiation rate α_a [1/s] |
|---|---|---|---|---|
| 33 | 4 | 1724 | 431 | 0.161 |
| 34 | 10 | 2080 | 208 | 0.191 |
| 35 | 25 | 7400 | 296 | 0.171 |
| 36 | 7 | 3591 | 513 | 0.117 |
| 37 | 20 | 8920 | 446 | 0.201 |
| 38 | 5 | 2255 | 451 | 0.115 |
| 39 | 31 | 11315 | 365 | 0.18 |
| 40 | 70 | 25200 | 360 | 0.187 |
| 41 | 2 | 1108 | 554 | 0.128 |
| 42 | - | - | 834 | - |
| 43 | 35 | 7630 | 218 | 0.207 |
| 44 | 8 | 3800 | 475 | 0.138 |
| 45 | 3 | 1260 | 420 | 0.143 |
| 46 | 29 | 12557 | 433 | 0.133 |
| 47 | 11 | 3212 | 292 | 0.174 |
| 48 | 2 | 780 | 390 | 0.117 |
| 49 | 10 | 2740 | 274 | 0.182 |
| 50 | 15 | 6270 | 418 | 0.152 |
| 51 | 1 | 454 | 454 | 0.0879 |
| 52 | 20 | 8780 | 439 | 0.188 |
| 53 | 8 | 3352 | 419 | 0.132 |
| 54 | 2 | 818 | 409 | 0.102 |
| 55 | 4 | 1520 | 380 | 0.142 |
| 56 | 6 | 2412 | 402 | 0.207 |
| 57 | 5 | 2585 | 517 | 0.0828 |
| 58 | 19 | 7847 | 413 | 0.134 |
| 59 | 22 | 8470 | 385 | 0.133 |
| 60 | 2 | 1094 | 547 | 0.0839 |
| 61 | 6 | 2370 | 395 | 0.179 |
| 62 | 2 | 1016 | 508 | 0.103 |
| 63 | 6 | 2736 | 456 | 0.15 |
| 64 | 2 | 864 | 432 | 0.114 |
| 65 | 15 | 5280 | 352 | 0.189 |
| 66 | 2 | 1840 | 920 | 0.0935 |
| 67 | 37 | 12432 | 336 | 0.171 |
| 68 | 4 | 2132 | 533 | 0.128 |
| 69 | 4 | 1768 | 442 | 0.166 |
| 70 | 19 | 4275 | 225 | 0.209 |
| 71 | 3 | 1428 | 476 | 0.13 |

| a (GO-Slim) | Number of mRNAs | Number of codons | Av. Length of 1 mRNA | Initiation rate α_a [1/s] |
|---|---|---|---|---|
| 72 | 119 | 26537 | 223 | 0.209 |
| 73 | 6 | 2874 | 479 | 0.119 |
| 74 | 8 | 2840 | 355 | 0.146 |
| 75 | 4 | 1748 | 437 | 0.112 |
| 76 | 58 | 13166 | 227 | 0.201 |
| 77 | 12 | 5964 | 497 | 0.133 |
| 78 | 2 | 814 | 407 | 0.123 |
| 79 | 5 | 1550 | 310 | 0.213 |
| 80 | 4 | 1928 | 482 | 0.158 |
| 81 | 4 | 1564 | 391 | 0.123 |
| 82 | 17 | 6851 | 403 | 0.134 |
| 83 | 3 | 1077 | 359 | 0.138 |
| 84 | 23 | 12650 | 550 | 0.19 |
| 85 | 11 | 4246 | 386 | 0.152 |
| 86 | 23 | 9844 | 428 | 0.122 |
| 87 | - | - | 364 | - |
| 88 | 6 | 3840 | 640 | 0.132 |
| 89 | 4 | 1728 | 432 | 0.14 |
| 90 | 4 | 1884 | 471 | 0.16 |
| 91 | 4 | 1300 | 325 | 0.129 |
| 92 | 3 | 990 | 330 | 0.124 |
| 93 | | | | |
| **Sum** | **1516** | **500124** | **-** | **-** |
| **Average** | **-** | **-** | **430** | **0.15** |

***Table S3:* List of amino acids (aa) with corresponding tRNAs (index i) and codons (index j).**

Gene copy number for each tRNA, wobble base pairing for each codon and the maximal hopping rates $k_{max}(j)$ for the downscaled cell. Note that, in general, the hopping rate depends on the number of charged tRNAs, the values given in this table correspond to all tRNAs are 100% charged. Note further, that the wobble factor $w(i)$ used in the equations in the text is given by $w(i) = (1 - wobble)$.

| aa | tRNA index i | tRNA | codon index j | codon | GCN | wobble | $k_{max}(j)$ |
|---|---|---|---|---|---|---|---|
| Ala | 1 | IGC | 1 | GCU | 11 | 0 | 21.39 |
| Ala | 1 | IGC | 2 | GCC | 11 | 0.36 | 13.69 |
| Ala | 2 | UGC | 3 | GCG | 6 | 0.6 | 4.67 |
| Ala | 2 | UGC | 4 | GCA | 6 | 0 | 11.66 |
| Arg | 3 | ICG | 5 | CGU | 7 | 0 | 13.61 |
| Arg | 3 | ICG | 6 | CGC | 7 | 0.36 | 8.71 |
| Arg | 3 | ICG | 7 | CGA | 7 | 0.36 | 8.71 |
| Arg | 4 | CCG | 8 | CGG | 1 | 0 | 1.94 |
| Arg | 5 | CCU | 9 | AGG | 1 | 0 | 1.94 |
| Arg | 6 | UCU | 10 | AGA | 12 | 0 | 23.33 |
| Asn | 7 | GUU | 11 | AAU | 11 | 0.39 | 13.04 |
| Asn | 7 | GUU | 12 | AAC | 11 | 0 | 21.39 |
| Asp | 8 | GUC | 13 | GAU | 16 | 0.39 | 18.97 |
| Asp | 8 | GUC | 14 | GAC | 16 | 0 | 31.11 |
| Cys | 9 | GCA | 15 | UGU | 4 | 0.39 | 4.74 |
| Cys | 9 | GCA | 16 | UGC | 4 | 0 | 7.78 |
| Gln | 10 | CUG | 17 | CAG | 1 | 0 | 1.94 |
| Gln | 11 | UUG | 18 | CAA | 9 | 0 | 17.50 |
| Glu | 12 | CUC | 19 | GAG | 2 | 0 | 3.89 |
| Glu | 13 | UUC | 20 | GAA | 15 | 0 | 29.16 |
| Gly | 14 | GCC | 21 | GGU | 16 | 0.39 | 18.97 |
| Gly | 14 | GCC | 22 | GGC | 16 | 0 | 31.11 |
| Gly | 15 | CCC | 23 | GGG | 2 | 0 | 3.89 |
| Gly | 16 | UCC | 24 | GGA | 3 | 0 | 5.83 |
| His | 17 | GUG | 25 | CAU | 8 | 0.39 | 9.49 |
| His | 17 | GUG | 26 | CAC | 8 | 0 | 15.55 |
| Ile | 18 | AAU | 27 | AUU | 13 | 0 | 25.27 |
| Ile | 18 | AAU | 28 | AUC | 13 | 0.36 | 16.17 |
| Ile | 19 | UAU | 29 | AUA | 2 | 0 | 3.89 |
| Leu | 20 | GAG | 30 | CUU | 1 | 0.39 | 1.19 |
| Leu | 20 | GAG | 31 | CUC | 1 | 0 | 1.94 |

| aa | tRNA index i | tRNA | codon index j | codon | GCN | wobble | $k_{max}(j)$ |
|---|---|---|---|---|---|---|---|
| Leu | 21 | UAG | 32 | CUG | 3 | 0.6 | 2.33 |
| Leu | 21 | UAG | 33 | CUA | 3 | 0 | 5.83 |
| Leu | 22 | CAA | 34 | UUG | 10 | 0 | 19.44 |
| Leu | 23 | UAA | 35 | UUA | 7 | 0 | 13.61 |
| Lys | 24 | CUU | 36 | AAG | 14 | 0 | 27.22 |
| Lys | 25 | UUU | 37 | AAA | 8 | 0 | 15.55 |
| Met | 26 | CAU | 38 | AUG | 11 | 0 | 21.39 |
| Phe | 27 | GAA | 39 | UUU | 11 | 0.39 | 13.04 |
| Phe | 27 | GAA | 40 | UUC | 11 | 0 | 21.39 |
| Pro | 28 | IGG | 41 | CCC | 2 | 0.36 | 2.49 |
| Pro | 28 | IGG | 42 | CCU | 2 | 0 | 3.89 |
| Pro | 29 | UGG | 43 | CCG | 10 | 0.6 | 7.78 |
| Pro | 29 | UGG | 44 | CCA | 10 | 0 | 19.44 |
| Ser | 30 | IGA | 45 | UCU | 11 | 0 | 21.39 |
| Ser | 30 | IGA | 46 | UCC | 11 | 0.36 | 13.69 |
| Ser | 31 | CGA | 47 | UCG | 1 | 0 | 1.94 |
| Ser | 32 | UGA | 48 | UCA | 4 | 0 | 7.78 |
| Ser | 33 | GCU | 49 | AGU | 2 | 0.39 | 2.37 |
| Ser | 33 | GCU | 50 | AGC | 2 | 0 | 3.89 |
| Thr | 34 | IGU | 51 | ACU | 11 | 0 | 21.39 |
| Thr | 34 | IGU | 52 | ACC | 11 | 0.36 | 13.69 |
| Thr | 35 | CGU | 53 | ACG | 1 | 0 | 1.94 |
| Thr | 36 | UGU | 54 | ACA | 5 | 0 | 9.72 |
| Trp | 37 | CCA | 55 | UGG | 6 | 0 | 11.66 |
| Tyr | 38 | GUA | 56 | UAU | 8 | 0.39 | 9.49 |
| Tyr | 38 | GUA | 57 | UAC | 8 | 0 | 15.55 |
| Val | 39 | IAC | 58 | GUU | 14 | 0 | 27.22 |
| Val | 39 | IAC | 59 | GUC | 14 | 0.36 | 17.42 |
| Val | 40 | CAC | 60 | GUG | 2 | 0 | 3.89 |
| Val | 41 | UAC | 61 | GUA | 3 | 0 | 5.83 |

*Table S4:* **Parameters used for the simulation**

Calculation of the catalytic rate $k_{cat}$ for each amino acid type from the estimated number of amino acids added/cell/ second and the number of synthetase/cell. The last column shows the number of tRNA copies/amino acid type.

| aa | amino acids added/cell/s | synthetase/cell | kcat (reactions/s) | tRNA copies |
|---|---|---|---|---|
| Ala | 190467 | 22920 | 8.31 | 17 |
| Arg | 104064 | 17540 | 5.93 | 21 |
| Asn | 117783 | 5950 | 19.80 | 11 |
| Asp | 144229 | 17660 | 8.17 | 16 |
| Cys | 24576 | 12970 | 1.89 | 4 |
| Gln | 87125 | 25830 | 3.37 | 10 |
| Glu | 171562 | 46100 | 3.72 | 17 |
| Gly | 157047 | 51950 | 3.02 | 21 |
| His | 48063 | 16730 | 2.87 | 8 |
| Ile | 152007 | 19790 | 7.68 | 16 |
| Leu | 211848 | 80830 | 2.62 | 21 |
| Lys | 183511 | 26230 | 7.00 | 22 |
| Met | 49471 | 50790 | 0.97 | 11 |
| Phe | 97812 | 10270 | 9.52 | 11 |
| Pro | 102638 | 13480 | 7.61 | 12 |
| Ser | 176762 | 18440 | 9.59 | 18 |
| Thr | 141140 | 30520 | 4.62 | 17 |
| Trp | 23667 | 11250 | 2.10 | 6 |
| Tyr | 75072 | 9500 | 7.90 | 8 |
| Val | 164360 | 6750 | 24.35 | 19 |

***Table S5;*** **Gene ontologies significantly enriched in, and common to, gene sets upregulated in response to Gln4 depletion and 3-AT treatment**

| ***p* values** | | | |
|---|---|---|---|
| ***GLN4*-depletion:** | **3AT treatment** | **Gene ontology class** | **Description** |
| 4.0586E-22 | 4.786E-26 | GO:0008652 | cellular amino acid biosynthetic process |
| 7.5122E-20 | 4.3473E-25 | GO:0016053 | organic acid biosynthetic process |
| 7.5122E-20 | 4.3473E-25 | GO:0046394 | carboxylic acid biosynthetic process |
| 1.0886E-21 | 5.2633E-25 | GO:1901607 | alpha-amino acid biosynthetic process |
| 9.2335E-21 | 6.3954E-23 | GO:1901605 | alpha-amino acid metabolic process |
| 1.7127E-17 | 4.1551E-20 | GO:0044283 | small molecule biosynthetic process |
| 2.0423E-16 | 4.1081E-19 | GO:0006520 | cellular amino acid metabolic process |
| 2.7156E-16 | 4.2686E-19 | GO:0019752 | carboxylic acid metabolic process |
| 1.3871E-15 | 1.3062E-18 | GO:0043436 | oxoacid metabolic process |
| 1.6251E-15 | 1.5912E-18 | GO:0006082 | organic acid metabolic process |
| 1.2274E-11 | 5.454E-17 | GO:0044281 | small molecule metabolic process |
| 3.667E-10 | 1.2398E-15 | GO:0009067 | aspartate family amino acid biosynthetic process |
| 8.1829E-11 | 9.9498E-13 | GO:0009066 | aspartate family amino acid metabolic process |
| 3.4377E-09 | 1.2241E-10 | GO:0009086 | methionine biosynthetic process |
| 4.3125E-09 | 3.3538E-10 | GO:0006526 | arginine biosynthetic process |
| 8.1479E-09 | 5.7122E-10 | GO:0006555 | methionine metabolic process |
| 4.5527E-08 | 6.6122E-10 | GO:0000096 | sulfur amino acid metabolic process |
| 2.7031E-05 | 8.1427E-10 | GO:0055114 | oxidation-reduction process |
| 1.5637E-08 | 8.2756E-10 | GO:0000097 | sulfur amino acid biosynthetic process |
| 2.3565E-10 | 4.2276E-09 | GO:0006790 | sulfur compound metabolic process |
| 1.2359E-07 | 2.145E-08 | GO:0016829 | lyase activity |
| 1.4968E-05 | 5.3124E-08 | GO:0016491 | oxidoreductase activity |
| 3.049E-11 | 8.1241E-08 | GO:0044272 | sulfur compound biosynthetic process |
| 2.0438E-05 | 3.8434E-07 | GO:0048037 | cofactor binding |
| 4.9575E-05 | 4.6508E-07 | GO:0008483 | transaminase activity |
| 4.9575E-05 | 4.6508E-07 | GO:0016769 | transferase activity, transferring nitrogenous groups |
| 1.4196E-09 | 3.0789E-06 | GO:0006525 | arginine metabolic process |
| 2.3477E-05 | 5.5375E-06 | GO:0044282 | small molecule catabolic process |
| 1.8657E-08 | 1.4377E-05 | GO:0009084 | glutamine family amino acid biosynthetic process |
| 1.2642E-07 | 2.2184E-05 | GO:1901565 | organonitrogen compound catabolic process |
| 1.2189E-08 | 3.2749E-05 | GO:0009064 | glutamine family amino acid metabolic process |

***Table S6;* Gene ontologies significantly enriched in, and common to, gene sets downregulated in response to Gln4 depletion and 3-AT treatment**

| ***p* values** | | | |
|---|---|---|---|
| ***GLN4*-depletion:** | **3AT treatment** | **Gene ontology class** | **Description** |
| 1.5662E-15 | 8.347E-92 | GO:0043228 | non-membrane-bounded organelle |
| 1.5662E-15 | 8.347E-92 | GO:0043232 | intracellular non-membrane-bounded organelle |
| 1.2258E-16 | 4.1402E-91 | GO:0030529 | intracellular ribonucleoprotein complex |
| 1.2258E-16 | 4.1402E-91 | GO:1990904 | ribonucleoprotein complex |
| 1.767E-53 | 4.8838E-64 | GO:0002181 | cytoplasmic translation |
| 9.9036E-40 | 9.3358E-55 | GO:0005840 | ribosome |
| 1.1195E-46 | 1.5979E-54 | GO:0044445 | cytosolic part |
| 1.8043E-26 | 4.8827E-52 | GO:0006412 | translation |
| 4.7545E-26 | 3.1621E-51 | GO:0043043 | peptide biosynthetic process |
| 1.3219E-22 | 4.5489E-47 | GO:0043604 | amide biosynthetic process |
| 1.7481E-23 | 2.6637E-46 | GO:0006518 | peptide metabolic process |
| 4.7194E-37 | 2.2651E-45 | GO:0044391 | ribosomal subunit |
| 2.7668E-37 | 2.1065E-43 | GO:0003735 | structural constituent of ribosome |
| 1.1541E-19 | 6.86E-40 | GO:0043603 | cellular amide metabolic process |
| 4.1786E-05 | 4.526E-36 | GO:0000462 | maturation of SSU-rRNA from tricistronic rRNA transcript (SSU-rRNA, 5.8S rRNA, LSU-rRNA) |
| 7.866E-31 | 6.5514E-33 | GO:0022627 | cytosolic small ribosomal subunit |
| 4.7415E-27 | 6.8294E-32 | GO:0022625 | cytosolic large ribosomal subunit |
| 9.7592E-26 | 3.8026E-31 | GO:0005622 | intracellular |
| 1.2174E-23 | 6.4895E-30 | GO:1901566 | organonitrogen compound biosynthetic process |
| 3.3777E-28 | 1.1283E-26 | GO:0005198 | structural molecule activity |
| 7.1011E-24 | 1.4982E-25 | GO:1901564 | organonitrogen compound metabolic process |
| 7.1097E-18 | 1.5934E-25 | GO:0015934 | large ribosomal subunit |
| 4.5621E-20 | 2.7434E-20 | GO:0015935 | small ribosomal subunit |
| 4.7639E-11 | 4.9283E-19 | GO:0044271 | cellular nitrogen compound biosynthetic process |
| 3.6033E-10 | 4.0626E-16 | GO:0005737 | cytoplasm |
| 2.4835E-10 | 3.0074E-14 | GO:0006407 | rRNA export from nucleus |
| 2.4835E-10 | 3.0074E-14 | GO:0051029 | rRNA transport |
| 8.0048E-06 | 5.3341E-12 | GO:0044267 | cellular protein metabolic process |
| 3.0854E-09 | 3.2521E-09 | GO:0000786 | nucleosome |
| 5.9599E-10 | 1.673E-08 | GO:0000788 | nuclear nucleosome |
| 5.8362E-07 | 1.3338E-07 | GO:0044815 | DNA packaging complex |

***Table S7;* Gene ontologies significantly enriched in, and common to, proteins upregulated in response to Gln4 depletion (SILAC analysis) and genes upregulated by 3-AT treatment (transcript profile analysis)**

| ***p* values** | | | |
|---|---|---|---|
| ***GLN4*-depletion:** | **3AT treatment** | **Gene ontology class** | **Description** |
| 1.2379E-28 | 4.786E-26 | GO:0008652 | cellular amino acid biosynthetic process |
| 1.4123E-29 | 4.3473E-25 | GO:0016053 | organic acid biosynthetic process |
| 1.4123E-29 | 4.3473E-25 | GO:0046394 | carboxylic acid biosynthetic process |
| 2.91E-26 | 5.2633E-25 | GO:1901607 | alpha-amino acid biosynthetic process |
| 1.4848E-27 | 6.3954E-23 | GO:1901605 | alpha-amino acid metabolic process |
| 7.2333E-30 | 4.1551E-20 | GO:0044283 | small molecule biosynthetic process |
| 1.6504E-28 | 4.1081E-19 | GO:0006520 | cellular amino acid metabolic process |
| 1.2902E-35 | 4.2686E-19 | GO:0019752 | carboxylic acid metabolic process |
| 2.2993E-35 | 1.3062E-18 | GO:0043436 | oxoacid metabolic process |
| 2.7255E-35 | 1.5912E-18 | GO:0006082 | organic acid metabolic process |
| 1.9364E-44 | 5.454E-17 | GO:0044281 | small molecule metabolic process |
| 1.1372E-10 | 1.2398E-15 | GO:0009067 | aspartate family amino acid biosynthetic process |
| 5.5567E-10 | 9.9498E-13 | GO:0009066 | aspartate family amino acid metabolic process |
| 6.8429E-08 | 4.9081E-11 | GO:0009085 | lysine biosynthetic process |
| 2.0743E-05 | 1.2241E-10 | GO:0009086 | methionine biosynthetic process |
| 3.9311E-09 | 3.0807E-10 | GO:0006553 | lysine metabolic process |
| 5.2225E-09 | 3.3538E-10 | GO:0006526 | arginine biosynthetic process |
| 5.1918E-06 | 7.1395E-10 | GO:0006081 | cellular aldehyde metabolic process |
| 3.8622E-05 | 8.2756E-10 | GO:0000097 | sulfur amino acid biosynthetic process |
| 2.6898E-07 | 2.6861E-08 | GO:0019878 | lysine biosynthetic process via aminoadipic acid |
| 1.8848E-40 | 4.5317E-08 | GO:0003824 | catalytic activity |
| 3.3076E-07 | 4.6508E-07 | GO:0008483 | transaminase activity |
| 3.3076E-07 | 4.6508E-07 | GO:0016769 | transferase activity, transferring nitrogenous groups |
| 6.9018E-07 | 3.0789E-06 | GO:0006525 | arginine metabolic process |
| 2.7726E-06 | 3.2472E-06 | GO:0030170 | pyridoxal phosphate binding |
| 6.3369E-14 | 5.5375E-06 | GO:0044282 | small molecule catabolic process |
| 1.9819E-06 | 1.4377E-05 | GO:0009084 | glutamine family amino acid biosynthetic process |
| 1.2904E-09 | 3.2749E-05 | GO:0009064 | glutamine family amino acid metabolic process |
| 1.2379E-28 | 4.786E-26 | GO:0008652 | cellular amino acid biosynthetic process |
| 1.4123E-29 | 4.3473E-25 | GO:0016053 | organic acid biosynthetic process |
| 1.4123E-29 | 4.3473E-25 | GO:0046394 | carboxylic acid biosynthetic process |

***Table S8:*** **Proteins identified through SILAC as reduced in concentration in response to Gln tRNA synthetase shut-off (at 1.4-fold or greater reduction)**

| Systematic gene name | Gene | Description | Log fold ratio: |
|---|---|---|---|
| YGR159C | NSR1 | Nucleolar protein that binds nuclear localization sequences; required for pre-rRNA processing | -0.5073126 |
| YKL180W | RPL17A | Ribosomal Protein of the Large subunit | -0.5087329 |
| YGL028C | SCW11 | Soluble Cell Wall protein | -0.5091384 |
| YPL211W | NIP7 | Nuclear ImPort | -0.5105569 |
| YGR285C | ZUO1 | ZUOtin | -0.5120751 |
| YLR354C | TAL1 | TransALdolase | -0.5151069 |
| YGR085C | RPL11B | Ribosomal Protein of the Large subunit | -0.5180315 |
| YNL220W | ADE12 | ADEnine requiring | -0.5214528 |
| YGL189C | RPS26A | Ribosomal Protein of the Small subunit | -0.526169 |
| YJL080C | SCP160 | S. cerevisiae protein involved in the Control of Ploidy | -0.5308698 |
| YJL136C | RPS21B | Ribosomal Protein of the Small subunit | -0.5385382 |
| YGL234W | ADE5,7 | ADEnine requiring | -0.5430009 |
| YHL015W | RPS20 | Ribosomal Protein of the Small subunit | -0.5574836 |
| YBR025C | OLA1 | Obg-Like ATPase | -0.5574836 |
| YMR058W | FET3 | FErrous Transport | -0.5584636 |
| YCR073W-A | SOL2 | Suppressor Of Los1-1 | -0.5614972 |
| YDL229W | SSB1 | Stress-Seventy subfamily B | -0.5638415 |
| YGL147C | RPL9A | Ribosomal Protein of the Large subunit | -0.5644269 |
| YLR448W | RPL6B | Ribosomal Protein of the Large subunit | -0.5651097 |
| YOL120C | RPL18A | Ribosomal Protein of the Large subunit | -0.5651097 |
| YER043C | SAH1 | S-Adenosyl-l-Homocysteine hydrolase | -0.5667665 |
| YNL112W | DBP2 | Dead Box Protein | -0.5734715 |
| YNL069C | RPL16B | Ribosomal Protein of the Large subunit | -0.5861162 |
| YGR214W | RPS0A | Ribosomal Protein of the Small subunit | -0.5934972 |
| YHR128W | FUR1 | 5-FluoroURidine resistant | -0.6005552 |
| YDR399W | HPT1 | Hypoxanthine guanine PhosphoribosylTransferase | -0.6039764 |
| YBR191W | RPL21A | Ribosomal Protein of the Large subunit | -0.6047356 |
| YNL096C | RPS7B | Ribosomal Protein of the Small subunit | -0.605684 |
| YDR025W | RPS11A | Ribosomal Protein of the Small subunit | -0.6060631 |
| YGL031C | RPL24A | Ribosomal Protein of the Large subunit | -0.6070107 |
| YBR031W | RPL4A | Ribosomal Protein of the Large subunit | -0.6193661 |
| YPR132W | RPS23B | Ribosomal Protein of the Small subunit | -0.6231177 |
| YJR010W | MET3 | METhionine requiring | -0.6291001 |
| YDR012W | RPL4B | Ribosomal Protein of the Large subunit | -0.6315234 |
| YAR071W | PHO11 | PHOsphate metabolism | -0.6408985 |
| YNL209W | SSB2 | Stress-Seventy subfamily B | -0.6429326 |
| YER036C | ARB1 | ATP-binding cassette protein involved in Ribosome Biogenesis | -0.6436715 |
| YOR167C | RPS28A | Ribosomal Protein of the Small subunit | -0.6438562 |

| | | | |
|---|---|---|---|
| YER177W | BMH1 | Brain Modulosignalin Homolog | -0.647729 |
| YMR116C | ASC1 | Absence of growth Suppressor of Cyp1 | -0.6493855 |
| YML056C | IMD4 | IMP Dehydrogenase | -0.6501212 |
| YLR325C | RPL38 | Ribosomal Protein of the Large subunit | -0.6520505 |
| YKL001C | MET14 | METhionine requiring | -0.652693 |
| YPL127C | HHO1 | Histone H One | -0.6548938 |
| YKR057W | RPS21A | Ribosomal Protein of the Small subunit | -0.6570912 |
| YPL131W | RPL5 | Ribosomal Protein of the Large subunit | -0.6931412 |
| YPL105C | SYH1 | SmY2 Homolog | -0.7021256 |
| YOL109W | ZEO1 | ZEOcin resistance | -0.7077898 |
| YAL003W | EFB1 | Elongation Factor Beta | -0.7094671 |
| YFL037W | TUB2 | TUBulin | -0.7099082 |
| YOR293W | RPS10A | Ribosomal Protein of the Small subunit | -0.7136958 |
| YGR234W | YHB1 | Yeast flavoHemogloBin | -0.7277898 |
| YGL123W | RPS2 | Ribosomal Protein of the Small subunit | -0.7376002 |
| YLR432W | IMD3 | IMP Dehydrogenase | -0.7382923 |
| YJR123W | RPS5 | Ribosomal Protein of the Small subunit | -0.7416611 |
| YKL216W | URA1 | URAcil requiring | -0.7431272 |
| YER102W | RPS8B | Ribosomal Protein of the Small subunit | -0.7438165 |
| YOR133W | EFT1 | Elongation Factor Two | -0.7651952 |
| YJR145C | RPS4A | Ribosomal Protein of the Small subunit | -0.7683326 |
| YLR048W | RPS0B | Ribosomal Protein of the Small subunit | -0.784504 |
| YNL208W | | | -0.7864291 |
| YBR189W | RPS9B | Ribosomal Protein of the Small subunit | -0.7957669 |
| YAL059W | ECM1 | ExtraCellular Mutant | -0.7998333 |
| YOR095C | RKI1 | Ribose-5-phosphate Ketol-Isomerase | -0.8062828 |
| YOR063W | RPL3 | Ribosomal Protein of the Large subunit | -0.8127036 |
| YHL033C | RPL8A | Ribosomal Protein of the Large subunit | -0.8310665 |
| YDR502C | SAM2 | S-AdenosylMethionine requiring | -0.8318772 |
| YNL178W | RPS3 | Ribosomal Protein of the Small subunit | -0.8366513 |
| YPL061W | ALD6 | ALdehyde Dehydrogenase | -0.8396372 |
| YFL045C | SEC53 | SECretory | -0.8425364 |
| YKL054C | DEF1 | RNAPII DEgradation Factor | -0.8565475 |
| YPL090C | RPS6A | Ribosomal Protein of the Small subunit | -0.8610025 |
| YNL113W | RPC19 | RNA Polymerase C | -0.8742855 |
| YJR070C | LIA1 | Ligand of eIF5A | -0.8855744 |
| YLR441C | RPS1A | Ribosomal Protein of the Small subunit | -0.8939834 |
| YGR210C | | | -0.9280481 |
| YPL043W | NOP4 | NucleOlar Protein | -0.9486008 |
| YDR099W | BMH2 | Brain Modulosignalin Homolog | -0.9661351 |
| YCR084C | TUP1 | dTMP-UPtake | -0.9889936 |
| YLR300W | EXG1 | EXo-1,3-beta-Glucanase | -1.0554733 |
| YML063W | RPS1B | Ribosomal Protein of the Small subunit | -1.0614306 |

| | | | |
|---|---|---|---|
| YPL079W | RPL21B | Ribosomal Protein of the Large subunit | -1.0688084 |
| YJL177W | RPL17B | Ribosomal Protein of the Large subunit | -1.0727918 |
| YLR206W | ENT2 | Epsin N-Terminal homology | -1.0994299 |
| YLR342W | FKS1 | FK506 Sensitivity | -1.1277653 |
| YLR249W | YEF3 | Yeast Elongation Factor | -1.1402551 |
| YOR252W | TMA16 | Translation Machinery Associated | -1.1767696 |
| YDR172W | SUP35 | SUPpressor | -1.2705887 |
| YBR092C | PHO3 | PHOsphate metabolism | -1.2849286 |
| YOR340C | RPA43 | RNA Polymerase A | -1.3810064 |
| YHR057C | CPR2 | Cyclosporin A-sensitive Proline Rotamase | -1.6014588 |
| YGR148C | RPL24B | Ribosomal Protein of the Large subunit | -1.6188494 |
| YFL007W | BLM10 | BLeoMycin resistance | -1.6273734 |
| YHR216W | IMD2 | IMP Dehydrogenase | -1.7110543 |
| YDR101C | ARX1 | Associated with Ribosomal eXport complex | -1.7300965 |
| YLR074C | BUD20 | BUD site selection | -1.7991702 |
| YNL141W | AAH1 | Adenine AminoHydrolase | -1.9590282 |
| YJR004C | SAG1 | Sexual AGglutination | -2.035448 |
| YML131W | | | -3.7443764 |
| YOR168W | GLN4 | GLutamiNe metabolism | -5.2097653 |

***Table S9:*** **Transcriptional response of translation initiation factor genes significantly repressed in the GLN4 tet-off strain in response to doxycycline treatment**

***Notes*** (a)*;* Significantly repressed (green shading, bold font) and induced genes (orange shading) are indicated. (b); A false discovery rate (FDR) of 0.05 was indicated the significance or otherwise of an adjusted *q* value (Benjamini-Hochberg).

| **Systematic gene name** | **Gene name** | **Initiation factor** | **Log$_2$ mRNA ratio (+ doxycycline/control)** | ***q* value** |
|---|---|---|---|---|
| YOL139C | *CDC33* | eIF4E | **-0.544** | 0.000128 |
| YGL049C | *TIF4632* | eIF4G | **-0.303** | 0.019159 |
| YGR162W | *TIF4631* | eIF4G | **-0.931** | 0.000128 |
| YKR059W | *TIF1* | eIF4A | **-0.932** | 0.000128 |
| YJL138C | *TIF2* | eIF4A | **-0.808** | 0.000128 |
| YPR163C | *TIF3* | eIF4B | **-1.207** | 0.000128 |
| YBR079C | *RPG1* | eIF3a | **-0.732** | 0.000128 |

| **Systematic gene name** | **Gene name** | **Initiation factor** | **$Log_2$ mRNA ratio (+ doxycycline/control)** | ***q* value** |
|---|---|---|---|---|
| YOR361C | *PRT1* | eIF3b | **-0.371** | 0.013003 |
| YMR309C | *NIP1* | eIF3c | **-0.321** | 0.0315117 |
| YDR429C | *TIF35* | eIF3g | **-1.09737** | 0.000128 |
| YMR146C | *TIF34* | eIF3i | **-1.25764** | 0.000128 |
| YLR192C | *HCR1* | eIF3j | **-0.503** | 0.000128 |
| YMR260C | *TIF11* | eIF1A | **-0.96252** | 0.000128 |
| YNL244C | *SUI1* | eIF1 | **-0.72058** | 0.000128 |
| YJR007w | *SUI2* | eIF2$\alpha$ | 0.195772 | 0.18319 |
| YPL237W | *SUI3* | eIF2$\beta$ | **-0.35716** | 0.025653 |
| YER025W | *GCD11* | eIF2$\gamma$ | **-0.76006** | 0.000128 |
| YKR026c | *GCN3* | eIF2B$\alpha$ | 0.362512 | 0.003837 |
| YLR291c | *GCD7* | eIF2B$\beta$ | 0.179204 | 0.181697 |
| YOR260w | *GCD1* | eIF2B$\gamma$ | **-0.44972** | 0.000592 |
| YGR083c | *GCD2* | eIF2B$\delta$ | **-0.28306** | 0.045961 |
| YDR211w | *GCD6* | eIF2B$\varepsilon$ | **-0.30185** | 0.029318 |
| YPR041w | *TIF5* | eIF5 | **-0.64725** | 0.000128 |
| YAL035w | *FUN12* | eIF5B | **-0.6244** | 0.000128 |

***Table S11:*** **Summary of the gene ontology categories assigned to the categorical number** a **used in the text.**

| a | GO-slim category | category |
|---|---|---|
| 0 | amino acid transport | GO:0006865 |
| 1 | biological process | GO:0008150 |
| 2 | carbohydrate metabolic process | GO:0005975 |
| 3 | carbohydrate transport | GO:0008643 |
| 4 | cell budding | GO:0007114 |
| 5 | cell morphogenesis | GO:0000902 |
| 6 | cell wall organization or biogenesis | GO:0071554 |
| 7 | cellular amino acid metabolic process | GO:0006520 |
| 8 | cellular ion homeostasis | GO:0006873 |
| 9 | cellular respiration | GO:0045333 |
| 10 | cellular response to DNA damage stimulus | GO:0006974 |
| 11 | chromatin organization | GO:0006325 |
| 12 | chromosome segregation | GO:0007059 |
| 13 | cofactor metabolic process | GO:0051186 |
| 14 | conjugation | GO:0000746 |
| 15 | cytokinesis | GO:0000910 |
| 16 | cytoplasmic translation | GO:0002181 |
| 17 | cytoskeleton organization | GO:0007010 |
| 18 | DNA recombination | GO:0006310 |
| 19 | DNA repair | GO:0006281 |
| 20 | DNA replication | GO:0006260 |
| 21 | DNA-templated transcription, initiation | GO:0006352 |
| 22 | DNA-templated transcription, termination | GO:0006353 |
| 22 | DNA-templated transcription, elongation | GO:0006354 |
| 22 | endocytosis | GO:0006897 |
| 23 | endosomal transport | GO:0016197 |
| 24 | exocytosis | GO:0006887 |
| 25 | generation of precursor metabolites and energy | GO:0006091 |
| 26 | Golgi vesicle transport | GO:0048193 |
| 27 | histone modification | GO:0016570 |
| 28 | invasive growth in response to glucose limitation | GO:0001403 |

| a | GO-slim category | category |
|---|---|---|
| 29 | ion transport | GO:0006811 |
| 30 | lipid metabolic process | GO:0006629 |
| 31 | lipid transport | GO:0006869 |
| 32 | meiotic cell cycle | GO:0051321 |
| 33 | membrane fusion | GO:0061025 |
| 34 | mitochondrial translation | GO:0032543 |
| 35 | mitochondrion organization | GO:0007005 |
| 36 | mitotic cell cycle | GO:0000278 |
| 37 | monocarboxylic acid metabolic process | GO:0032787 |
| 38 | mRNA processing | GO:0006397 |
| 39 | nuclear transport | GO:0051169 |
| 40 | nucleobase-containing compound transport | GO:0015931 |
| 41 | nucleobase-containing small molecule metabolic process | GO:0055086 |
| 41 | nucleus organization | GO:0006997 |
| 42 | oligosaccharide metabolic process | GO:0009311 |
| 43 | organelle assembly | GO:0070925 |
| 44 | organelle fusion | GO:0048284 |
| 45 | organelle fission | GO:0048285 |
| 45 | organelle inheritance | GO:0048308 |
| 46 | other | - |
| 47 | peptidyl-amino acid modification | GO:0018193 |
| 48 | peroxisome organization | GO:0007031 |
| 49 | protein alkylation and acylation | GO:0008213 |
| 50 | protein acylation | GO:0043543 |
| 50 | protein complex biogenesis | GO:0070271 |
| 51 | protein dephosphorylation | GO:0006470 |
| 52 | protein folding | GO:0006457 |
| 53 | protein glycosylation | GO:0006486 |
| 54 | protein lipidation | GO:0006497 |
| 55 | protein maturation | GO:0051604 |
| 56 | protein modification by small protein conjugation or removal | GO:0070647 |
| 57 | protein phosphorylation | GO:0006468 |
| 58 | protein targeting | GO:0006605 |

| a | GO-slim category | category |
|---|---|---|
| 59 | proteolysis involved in cellular protein catabolic process | GO:0051603 |
| 60 | pseudohyphal growth | GO:0007124 |
| 61 | regulation of cell cycle | GO:0051726 |
| 62 | regulation of DNA metabolic process | GO:0051052 |
| 63 | regulation of organelle organization | GO:0033043 |
| 64 | regulation of protein modification process | GO:0031399 |
| 65 | regulation of translation | GO:0006417 |
| 66 | regulation of transport | GO:0051049 |
| 67 | response to chemical | GO:0042221 |
| 68 | response to heat | GO:0009408 |
| 69 | response to osmotic stress | GO:0006970 |
| 70 | response to oxidative stress | GO:0006979 |
| 71 | response to starvation | GO:0042594 |
| 72 | ribosomal subunit export from nucleus | GO:0000054 |
| 73 | ribosome assembly | GO:0042255 |
| 73 | ribosomal large subunit biogenesis | GO:0042273 |
| 73 | ribosomal small subunit biogenesis | GO:0042274 |
| 73 | RNA catabolic process | GO:0006401 |
| 74 | RNA modification | GO:0009451 |
| 75 | RNA splicing | GO:0008380 |
| 76 | rRNA processing | GO:0006364 |
| 77 | signaling | GO:0023052 |
| 78 | snoRNA processing | GO:0043144 |
| 79 | sporulation | GO:0043934 |
| 80 | telomere organization | GO:0032200 |
| 81 | transcription from RNA polymerase I promoter | GO:0006360 |
| 82 | transcription from RNA polymerase II promoter | GO:0006366 |
| 83 | transcription from RNA polymerase III promoter | GO:0006383 |
| 84 | translational elongation | GO:0006414 |
| 85 | translational initiation | GO:0006413 |
| 86 | transmembrane transport | GO:0055085 |
| 87 | transposition | GO:0032196 |
| 88 | tRNA aminoacylation for protein translation | GO:0006418 |

| a | GO-slim category | category |
|---|---|---|
| 89 | tRNA processing | GO:0008033 |
| 90 | vacuole organization | GO:0007033 |
| 91 | vesicle organization | GO:0016050 |
| 92 | vitamin metabolic process | GO:0006766 |

#### REFERENCES


1. Gillespie,D.T. (1977) Exact stochastic simulation of coupled chemical reactions. *J. Phys. Chem.*, **81**, 2340–2361.

2. Miura,F., Kawaguchi,N., Yoshida,M., Uematsu,C., Kito,K., Sakaki,Y. and Ito,T. (2008) Absolute quantification of the budding yeast transcriptome by means of competitive PCR between genomic and complementary DNAs. *BMC Genomics*, **9**, 574.

3. Waldron,C. and Lacroute,F. (1975) Effect of growth rate on the amounts of ribosomal and transfer ribonucleic acids in yeast. *J. Bacteriol.*, **122**, 855–865.

4. Warner,J.R. (1999) The economics of ribosome biosynthesis in yeast. *Trends Biochem. Sci.*, **24**, 437–440.

5. Chan,P.P. and Lowe,T.M. (2009) GtRNAdb: a database of transfer RNA genes detected in genomic sequence. *Nucleic Acids Res.*, **37**, D93-7.

6. Arava,Y., Wang,Y., Storey,J.D., Liu,C.L., Brown,P.O. and Herschlag,D. (2003) Genome-wide analysis of mRNA translation profiles in *Saccharomyces cerevisiae*. *Proc. Natl. Acad. Sci. U. S. A.*, **100**, 3889–3894.

7. von der Haar,T. (2008) A quantitative estimation of the global translational activity in logarithmically growing yeast cells. *BMC Syst. Biol.*, **2**, 87.

8. Ghaemmaghami,S., Huh,W.K., Bower,K., Howson,R.W., Belle,A., Dephoure,N., O'Shea,E.K. and Weissman,J.S. (2003) Global analysis of protein expression in yeast. *Nature*, **425**, 737–741.

9. Dittmar,K.A., Sorensen,M.A., Elf,J., Ehrenberg,M. and Pan,T. (2005) Selective charging of tRNA isoacceptors induced by amino-acid starvation. *EMBO Rep.*, **6**, 151–157.

10. Varshney,U., Lee,C.P. and RajBhandary,U.L. (1991) Direct analysis of aminoacylation levels of tRNAs in vivo. Application to studying recognition of *Escherichia coli* initiator tRNA mutants by glutaminyl-tRNA synthetase. *J. Biol. Chem.*, **266**, 24712–24718.

11. Sorensen,M.A. (2001) Charging levels of four tRNA species in *Escherichia coli* Rel(+) and Rel(-) strains during amino acid starvation: a simple model for the effect of ppGpp on translational accuracy. *J. Mol. Biol.*, **307**, 785–798.

12. Jakubowski,H. and Goldman,E. (1984) Quantities of individual aminoacyl-tRNA families and their turnover in *Escherichia coli*. *J. Bacteriol.*, **158**, 769 LP – 776.

13. Kemp,A.J., Betney,R., Ciandrini,L., Schwenger,A.C.M., Romano,M.C. and Stansfield,I. (2013) A yeast tRNA mutant that causes pseudohyphal growth exhibits reduced rates of CAG codon translation. *Mol. Microbiol.*, **87**.

14. Gilchrist,M.A. and Wagner,A. (2006) A model of protein translation including codon bias, nonsense errors, and ribosome recycling. *J. Theor. Biol.*, **239**, 417–434.

15. Bjork,G.R., Huang,B., Persson,O.P. and Bystrom,A.S. (2007) A conserved modified wobble nucleoside (mcm5s2U) in lysyl-tRNA is required for viability in yeast. *RNA*, **13**, 1245–1255.

16. Sorensen,M.A. and Pedersen,S. (1991) Absolute in vivo translation rates of individual codons in *Escherichia coli.* The two glutamic acid codons GAA and GAG are translated with a threefold difference in rate. *J. Mol. Biol.*, **222**, 265–280.

17. Shah,P., Ding,Y., Niemczyk,M., Kudla,G. and Plotkin,J.B. (2013) Rate-limiting steps in yeast protein translation. *Cell*, **153**, 1589–1601.

18. Sin,C., Chiarugi,D. and Valleriani,A. (2016) Quantitative assessment of ribosome drop-off in E. coli. *Nucleic Acids Res.*, **44**, 2528–2537.

19. Ciandrini,L., Stansfield,I. and Romano,M.C. (2013) Ribosome traffic on mRNAs maps to gene ontology: genome-wide quantification of translation initiation rates and polysome size regulation. *PLoS Comput. Biol.*, **9**, e1002866.

20. Uter,N.T., Gruic-Sovulj,I. and Perona,J.J. (2005) Amino Acid-dependent Transfer RNA Affinity in a Class I Aminoacyl-tRNA Synthetase. *J. Biol. Chem.*, **280**, 23966–23977.

21. Huynh,L.-N., Thangavel,M., Chen,T., Cottrell,R., Mitchell,J.M. and Prætorius-Ibba,M. (2010) Linking tRNA localization with activation of nutritional stress responses. *Cell Cycle*, **9**, 3184–3190.

22. W. von Bloh. (2008) Sequential and Parallel Implementation of Networks. In Graben, P., Zhou, C., Thiel, M., and Kurths J., *Lectures in Supercomputational Neuroscience; Understanding Complex Systems*. Springer: Complexity.

23. Chelliah, V. et al. BioModels: ten-year anniversary. Nucl. Acids Res. 2015, 43(Database issue):D542-8 Chelliah,V., Juty,N., Ajmera,I., Ali,R., Dumousseau,M., Glont,M., Hucka,M., Jalowicki,G., Keating,S., Knight-Schrijver,V., Lloret-Villas,A., Natarajan,K.N., Pettit,J,B,, Rodriguez,N., Schubert,M,. Wimalaratne,S.M., Zhao,Y., Hermjakob,H., Le Novère,N. and Laibe,C. (2015) BioModels: ten-year anniversary. *Nucleic Acids Res*., **43** (Database issue):D542-8.
